# Convergence of orphan quality control pathways at a ubiquitin chain-elongating ligase

**DOI:** 10.1101/2024.08.07.607117

**Authors:** Sara Carrillo Roas, Yuichi Yagita, Paul Murphy, Robert Kurzbauer, Tim Clausen, Eszter Zavodszky, Ramanujan S. Hegde

## Abstract

Unassembled and partially assembled subunits of multi-protein complexes have emerged as major quality control clients, particularly under conditions of imbalanced gene expression such as stress, aging, and aneuploidy. The factors and mechanisms that eliminate such orphan subunits to maintain protein homeostasis are incompletely defined. Here, we show that the UBR4-KCMF1 ubiquitin ligase complex is required for efficient degradation of multiple unrelated orphan subunits from the chaperonin, proteasome cap, proteasome core, and a protein targeting complex. Epistasis analysis in cells and reconstitution studies *in vitro* show that the UBR4-KCMF1 complex acts downstream of a priming ubiquitin ligase that first mono-ubiquitinates orphans. UBR4 recognizes both the orphan and its mono-ubiquitin and builds a K48-linked poly-ubiquitin degradation signal. The discovery of a convergence point for multiple quality control pathways may explain why aneuploid cells are especially sensitive to loss of UBR4 or KCMF1 and identifies a potential vulnerability across many cancers.

## Introduction

Excess subunits of multimeric complexes are eliminated by quality control pathways to maintain cellular proteostasis and organism health^1–3^. Such ‘orphaned’ subunits arise from noise in gene expression, generating imbalances in subunit stoichiometry^4–6^. Subunit imbalances can be exaggerated by conditions that markedly re-wire gene expression pathways, such as environmental stresses, cancers, and possibly even aging^7–10^. Subunits of multimeric complexes are among the most abundant rapidly-degraded nascent proteins in both cancer and normal cells, indicating that orphans comprise the majority of quality control substrates in metazoans^11,12^. Thus, assembly quality control (AQC) pathways are emerging as central to proteostasis, making their identification of major importance.

Although AQC operates in many cellular compartments, most major protein complexes assemble partially or completely in the cytosol^13–16^. Several cytosolic AQC factors of the ubiquitin-proteasome system have been identified and dissected to various mechanistic depths. Two paradigms have emerged for orphan recognition by E3 ligases^2,3,12,17–21^. First, the ligase can directly recognize the orphan via region(s) that are occluded by a targeting factor, assembly chaperone, another subunit or intramolecular folding. Second, ligase recognition can be indirect and involve an assembly chaperone or other adaptor that provides substrate specificity.

Ligases that can operate by direct recognition during AQC include HUWE1 and UBE2O, both of which can ubiquitinate at least some of their targets in a purified system^17–19,21^. By contrast, HERC1 and HERC2 are ubiquitin ligases that operate by an adaptor-mediated mechanism^12,20^. HERC1 recognition of orphaned PSMC5, a 19S proteasome subunit, requires its dedicated assembly factor PAAF1. Analogously, HERC2 recognizes most subunits of the cytosolic chaperonin (CCT) via the adapter protein ZNRD2. Notably, each of these recognition mechanisms seems to be highly specific to a small subset of structurally related AQC clients, indicating that different types of orphans require different recognition factors. Thus, the AQC factors for many types of orphan subunits probably remain unidentified.

Defining the complete set of AQC pathways is likely to be of considerable value because impaired AQC can lead to disease, particularly in the nervous system^9,12,22–26^. Furthermore, AQC pathways may be attractive targets in aneuploid cancers. This is because a large body of literature indicates that aneuploidy markedly increases imbalances in subunit stoichiometry, imposes a major proteostatic burden, and leads to reduced fitness at the cell and organism levels^27–31^. The ability of aneuploid cancers to overcome this fitness cost^32^ hints at the possibility that they are highly dependent on AQC.

We therefore searched for additional AQC factors, leading us to a ubiquitin ligase complex comprised of UBR4 and KCMF1 that we find operates downstream of multiple known AQC ligases including HERC1, HERC2 and HUWE1. Mechanistic analyses indicate that the UBR4-KCMF1 complex recognizes orphans that are first mono-ubiquitinated by a ‘priming’ ligase, then commits orphans for degradation via assembly of a K48-linked poly-ubiquitin chain. The convergence of multiple AQC pathways at the UBR4-KCMF1 complex may explain why it is preferentially important for growth of cancer cells with a high aneuploidy index^33,34^.

## Results

### Orphaned 20S proteasome subunits are subject to quality control

Earlier analyses indicated imbalanced expression of subunits of both the 20S core proteasome and its 19S regulatory particle^12^. Whereas HERC1 was identified as an AQC factor for a subset of 19S subunits (e.g., PSMC5), it did not mediate degradation of orphaned PSMB4, a subunit of the 20S β-ring (Fig. S1A). To monitor orphaned β-ring subunit degradation, we generated fluorescent reporters of each subunit (PSMB1 through PSMB7) fused to GFP at either the N- or C-terminus (Fig. 1A). Relative to an inline RFP control separated by a P2A ribosome skipping sequence, each PSMB reporter was degraded when overexpressed in cultured cells (Fig. 1B,C; Fig. S1B). Degradation efficiency varied across subunits and was partially influenced by the GFP tag’s position. Because PSMB4 was among the most effectively degraded orphan subunits, we focused on this for further characterization.

**Figure 1.**
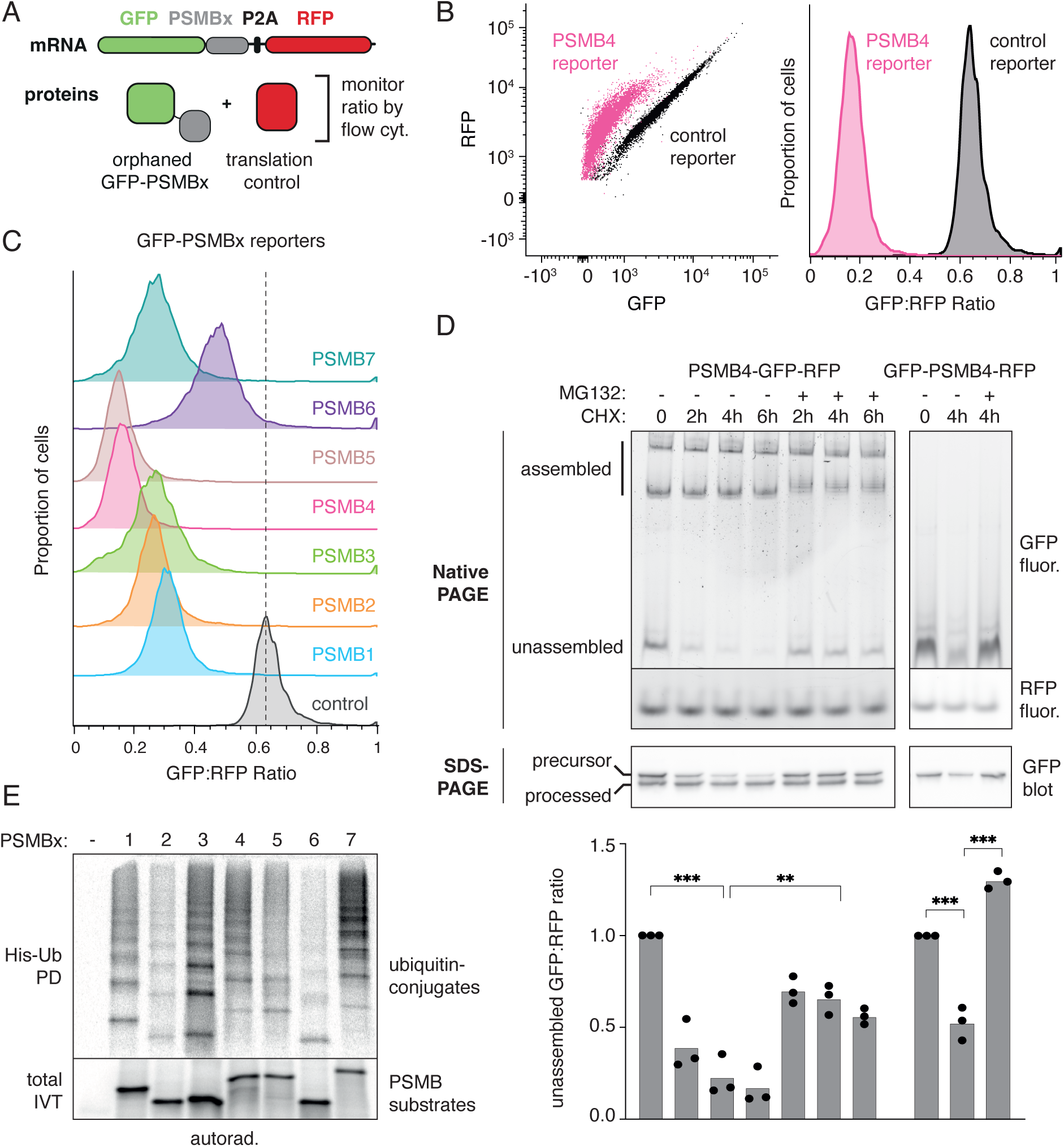
Orphaned PSMB subunits are targeted for degradation. **(A)** Diagram of the dual-color reporter mRNA and the expected protein products. GFP-tagged PSMB subunits and RFP are translated from the same mRNA but as separate products due to a viral P2A ribosomal skipping sequence. Relative GFP-tagged PSMB stability is measured by flow cytometry using the GFP:RFP fluorescence ratio. **(B)** Expression of the GFP-PSMB4 fluorescent reporter (pink) or control GFP reporter lacking PSMB4 (black) stably integrated into inducible HEK293 Flp-in TREx cells was induced for 48 h with 10 ng/pL doxycycline (dox), then analyzed by flow cytometry. Overlaid scatter plots of individual cells (left) and the corresponding histograms of the GFP:RFP ratio (right) are shown. **(C)** Expression of the seven GFP-PSMBx reporters compared to the control reporter was induced and analyzed as in B. Histograms of GFP:RFP ratio are shown, with the dotted line indicating the peak of the control histogram. The PSMB4 histogram from B is shown again for reference. **(D)** Expression of the PSMB4-GFP or GFP-PSMB4 reporters was induced for 18 h with dox. After removal of dox, cells were treated with 100 pg/mL cycloheximide (CHX) and 20 pM MG132 as indicated. Cell lysates were analyzed by Native PAGE and in-gel fluorescence to detect GFP and RFP, or denaturing SDS-PAGE followed by immunoblot for GFP. The ratio of unassembled GFP to RFP from native in-gel fluorescence was quantified from three independent experiments, normalized to the untreated sample (lane 1) and plotted below the gels. Statistical comparisons between key samples by Student’s t-test are indicated: ** indicates p<0.01 and *** indicates p<0.001. **(E)** PSMB subunits fused to a TwinStrep tag (TST) were translated in rabbit reticulocyte lysate (RRL) with ^35^S-methione and His-tagged ubiquitin (His-Ub). The samples were analyzed by SDS-PAGE and autoradiography either directly (total IVT) or following a denaturing His-Ub pulldown (His-Ub PD). See also Fig. SI.

Analysis by native gel electrophoresis showed that C-terminally tagged PSMB4-GFP assembled into complexes that correspond to 20S and 26S proteasome particles, whereas N-terminally tagged GFP-PSMB4 was mostly unassembled (Fig. 1D). Consistent with this interpretation, denaturing gels showed that assembly-dependent processing of an N-terminal pro-peptide occurred for PSMB4-GFP, but not GFP-PSMB4. In both cases, a substantial unassembled population was seen on native gels, and this population selectively disappeared upon translation inhibition with cycloheximide (CHX). Proteasome inhibition (with MG132) during CHX treatment inhibited unassembled PSMB4 degradation. Denaturing gels showed that the unprocessed product of PSMB4-GFP was selectively degraded during CHX-chase and stabilized by MG132 (Fig. 1D).

Each of the seven tagged PSMB subunits were ubiquitinated when translated *in vitro* without other PSMB subunits in rabbit reticulocyte lysate (Fig. 1E). Native affinity purification of PSMB4 from the translation reaction showed that it co-purified with E3 ubiquitin ligase(s), as demonstrated by additional ubiquitination upon addition of E1, E2, ubiquitin, and ATP (Fig S1C). These results indicate that orphaned PSMB subunits are subjected to quality control and degraded in a proteasome-dependent manner in cells and poly-ubiquitinated *in vitro*. Furthermore, PSMB4 interactions with E3 ligases *in vitro* are evidently sufficiently stable to permit co-purification, providing a route to their subsequent identification.

### The UBR4-KCMF1 complex engages orphaned PSMBs for degradation

Candidate AQC factors for orphaned PSMB subunits were identified from native affinity purified *in vitro*-translated subunits subjected to mass spectrometry (Fig. 2A, S2; Table S1). As expected, and consistent with analogous experiments with orphaned CCT and 19S subunits^12,20^, some partial assembly seems to occur, as evidenced by recovery of various proteasome subunits and assembly chaperones. In addition, several components of the ubiquitination machinery were recovered with most or all PSMBs. The recovery of Cullin1 and Skp1 indicated a role for the SCF ubiquitin ligase complex, consistent with partial stabilization of orphan PSMBs upon inhibition of Cullin-type ligases via a NEDDylation inhibitor (Fig. S3A). However, no effect was observed on orphaned GFP-PSMB4 upon knockdown (KD) of individual Cullin adaptors, perhaps indicating redundancy (Fig. S3B).

**Figure 2.**
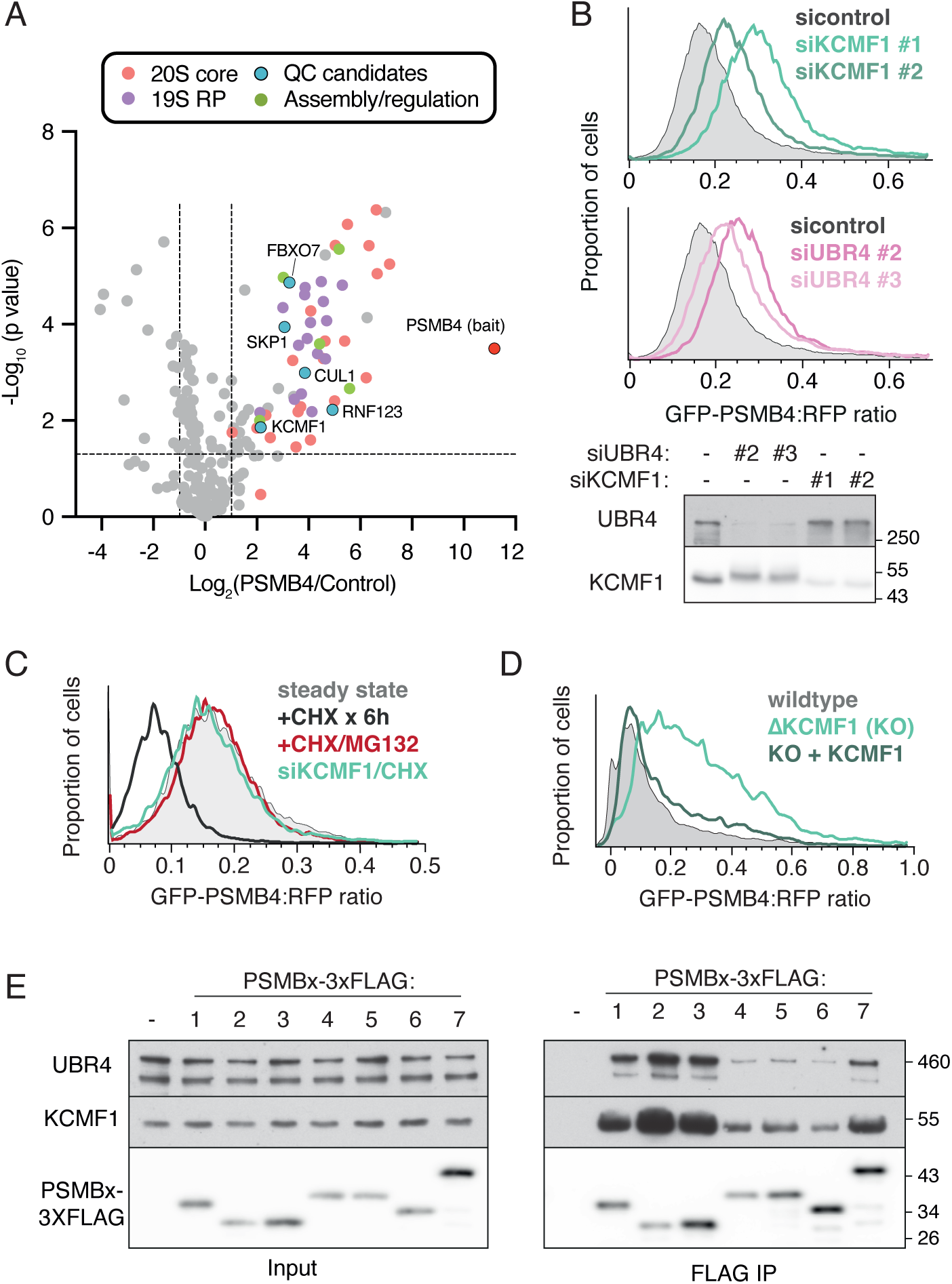
The UBR4-KCMF1 complex is required for the degradation of orphaned PS MB subunits. **(A)** PSMB4-TST was translated in RRL, affinity-purified under native conditions and analyzed by label-free quantitative mass spectrometry. The volcano plot shows proteins enriched in the PSMB4 pulldown compared to a mock translation (negative control). Different classes of proteins are indicated in differently colored dots (RP is regulatory particle), with PSMB4 and quality control candidates labeled. **(B)** Stable-inducible cells containing the GFP-PSMB4 reporter were transfected with the indicated siRNAs and grown for 72Һ. Expression of the GFP-PSMB4 reporter was induced with dox for the final 18h before analysis by flow cytometry (top) or immunoblot (bottom). **(C)** GFP-PSMB4 reporter cell lines were transfected with non-targeting control or KCMF1 siRNAs for a total of 72Һ. GFP-PSMB4 reporter expression was induced with dox for 18h, and removed prior to treatment with 100 pg/mL CHX and 20 pM MG132 for the final 6Һ, where indicated. Cells were then analyzed by flow cytometry. **(D)** Wildtype and KCMF1 KO (AKCMF1) cells were transiently co-transfected with the GFP-PSMB4 reporter and KCMFl-3xFLAG constructs, as indicated, for 48Һ. Cells were then analyzed by flow cytometry. (E) PSMB subunits C-terminally tagged with a 3xFLAG tag were translated in RRL and immunoprecipitated (FLAG IP) under native conditions. Input and IP samples were analyzed by immunoblot for the indicated proteins. See also Fig. S2-S4, and Table SI.

Among the other interacting ubiquitination factors, a small steady-state stabilization of GFP-PSMB4 was seen with RNF123 KD (Fig S3B), and a more substantive stabilization with KCMF1 KD (Fig. 2B). Using the more sensitive CHX-chase format, we observed that the extent of GFP-PSMB4 stabilization with KCMF1 KD was comparable to proteasome inhibition with MG132 in both native gel (Fig S3C) and flow cytometry assays (Fig. 2C). GFP-PSMB4 stabilization was also seen in KCMF1 knockout (KO) cells, where reporter degradation was restored by exogenous KCMF1 (Fig. 2D). Thus, KCMF1 plays a crucial and non-redundant role in orphaned PSMB4 degradation.

Consistent with KCMF1 forming a complex with UBR4, a giant E3 ligase^35,36^, the two endogenous proteins co-sedimented through a sucrose gradient (Fig. S4A). Furthermore, UBR4 co-immunoprecipitated exogenous tagged KCMF1 (Fig. S4B), and UBR4 KD led to GFP-PSMB4 stabilization, as seen with KCMF1 KD (Fig. 2B). The extent of GFP-PSMB4 stabilization with UBR4 KD was comparable to proteasome inhibition with MG132, and no additive effect was seen with UBR4 and KCMF1 double KD (Fig S3C). These results indicate that the UBR4-KCMF1 complex, and not either alone, mediates GFP-PSMB4 degradation. Most other PSMB subunit reporters were also stabilized, albeit to varying degrees, by UBR4 or KCMF1 KD (Fig. S4C). Both endogenous KCMF1 and endogenous UBR4 were co-immunoprecipitated by FLAG-tagged PSMB subunits translated *in vitro* or overexpressed in cultured cells (Fig. 2E, S4D). Thus, the UBR4-KCMF1 complex physically interacts with orphaned PSMB subunits and is required for their efficient degradation.

### Multiple AQC pathways involve the UBR4-KCMF1 complex

Many AQC factors show a high degree of specificity towards their substrates and do not seem to show compensation or overlap. For example, clients of HERC1 and HERC2 (PSMC5 and CCT4, respectively) cannot engage or use the non-cognate pathway^12,20^. Surprisingly however, orphaned CCT4-GFP degradation was strongly impacted by either UBR4 or KCMF1 KD to an extent comparable to that seen with HERC2 KD (Fig 3A). Similarly, UBR4 and KCMF1 were also needed for degradation of orphaned UBL4A (an established HUWE1 client^37^ that is normally part of the BAG6 complex^38^), and orphaned PSMC5 (an established HERC1 client^12^) (Fig 3A; Fig S5A,B). By contrast, NCOA4, a non-orphan HERC2 substrate that is mostly unstructured and is recognized by a different mechanism^39^, was not dependent on UBR4 or KCMF1 (Fig. 3A). Thus, multiple previously established AQC pathways, but not all degradation pathways, rely on the UBR4-KCMF1 complex.

**Figure 3.**
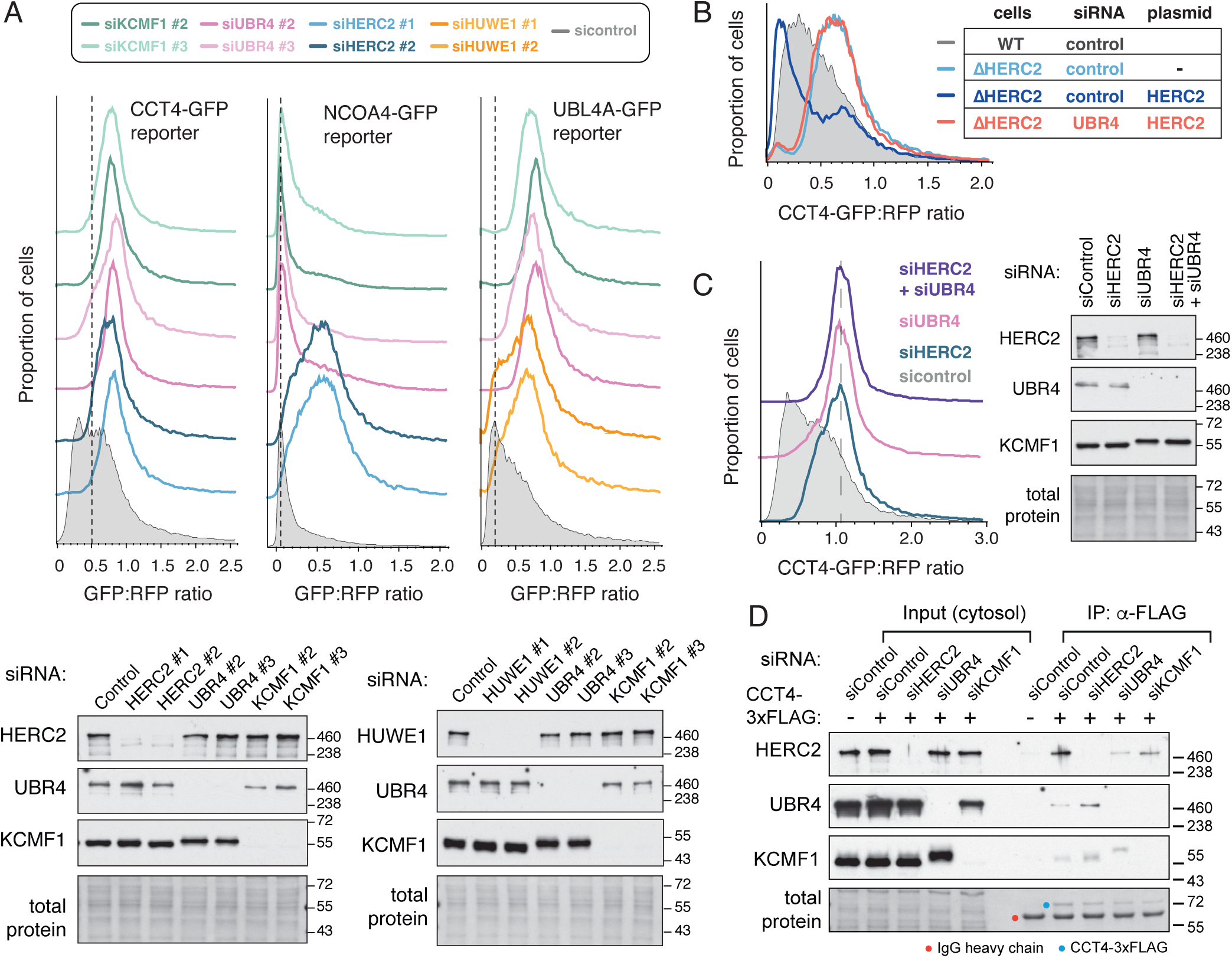
Multiple types of orphaned proteins depend on the UBR4-KCMF1 complex for efficient degradation. **(A)** HEK293T cells treated with control, KCMF1, UBR4, HERC2 or HUWE1-targeting siRNAs were transiently transfected with CCT4, NC0A4 or UBL4A fluorescent reporters, as indicated. Cells were then analyzed by flow cytometry (top) or immunoblot (bottom). **(B)** Wildtype and AHERC2 cells treated with control or UBR4-targeting siRNAs were co-transfected with the CCT4-GFP reporter and HERC2, as indicated, and analyzed by flow cytometry. **(C)** Cells treated with control, HERC2 or UBR4-targeting siRNAs were transfected with the CCT4-GFP reporter and analyzed by flow cytometry (left) or immunoblot (right). **(D)** HEK293T cells treated with control, HERC2, KCMF1 or UBR4-targeting siRNA were transfected with CCT4-3xFLAG, then subjected to anti-FLAG IP under native conditions. Input and IP samples were analyzed by immunoblot. See also Fig. S5.

To understand the relationship between the UBR4-KCMF1 complex and AQC, we focused on the CCT4-HERC2 system because it is understood in the greatest mechanistic depth^20^. CCT4-GFP dependence on the UBR4-KCMF1 complex was not due to the loss of either HERC2 or its adaptor ZNRD2 (Fig. 3A; Fig. S5C). Orphaned CCT4-GFP degradation could be restored to UBR4 or KCMF1 KO cells upon re-expression of exogenous UBR4 or KCMF1, respectively (Fig. S5D). As seen previously^20^, HERC2 overexpression in HERC2 KO cells drives CCT4-GFP degradation even beyond that seen in WT cells (Fig. 3B). Strikingly however, silencing UBR4 rendered HERC2 overexpression completely inert toward CCT4-GFP. Thus, the ability of the HERC2-ZNRD2 complex to degrade CCT4-GFP is strictly dependent on the UBR4-KCMF1 complex, arguing that they act sequentially in the same pathway. Consistent with this conclusion, depleting both HERC2 and UBR4 did not stabilize CCT4-GFP any more than depleting either factor alone (Fig. 3C).

Co-IP experiments showed that the UBR4-KCMF1 complex interacts with most overexpressed FLAG-tagged CCT subunits (Fig. S5E). CCT8, which is not recognized effectively as an orphan and hence not degraded^20^, interacted minimally with the UBR4-KCMF1 complex. Neither HERC2 KD nor UBR4 KD prevented CCT4-GFP from interacting with the KCMF1-UBR4 complex or HERC2, respectively (Fig. 3D). Yet, the remaining ligase, despite interacting with orphaned CCT4-GFP, was incapable of mediating its degradation. Considered together, the data indicate that the HERC2-ZNRD2 complex and the UBR4-KCMF1 complex can each interact with CCT4-GFP mostly independently of each other, yet act together in the same pathway to mediate CCT4-GFP degradation.

### Sequential ubiquitination by HERC2 and the UBR4-KCMF1 complex

A multi-ligase requirement for substrate degradation could suggest that each ligase contributes different parts of the final poly-ubiquitin degradation signal. Earlier reconstitution experiments with purified factors showed that the HERC2-ZNRD2 complex directly recognizes CCT4 and appends between one to three mono ubiquitins^20^. Because this low level of multi-mono-ubiquitination is typically insufficient to mediate proteasome degradation^40^, we tested multi-mono-ubiquitinated CCT4 for further modification by the UBR4-KCMF1 complex.

In this sequential ubiquitination experiment (Fig. 4A), radiolabelled CCT4 produced using purified *E. coli* translation factors and ribosomes (the so-called PURE system^41^) was incubated with recombinant HERC2, the E2 enzyme UBE2D1, E1 enzyme, His-tagged ubiquitin, and ATP. Multi-mono-ubiquitinated CCT4 was natively purified via His-ubiquitin and used as a substrate for recombinant UBR4-KCMF1 complex, its cognate E2 enzyme UBE2A^36^, E1 enzyme, FLAG-tagged ubiquitin, and ATP. Recovery of FLAG-ubiquitin products showed that multi-mono-ubiquitinated CCT4 became poly-ubiquitinated by the UBR4-KCMF1 complex as evidenced by the appearance of high molecular weight products (Fig. 4B, lane 8).

**Figure 4.**
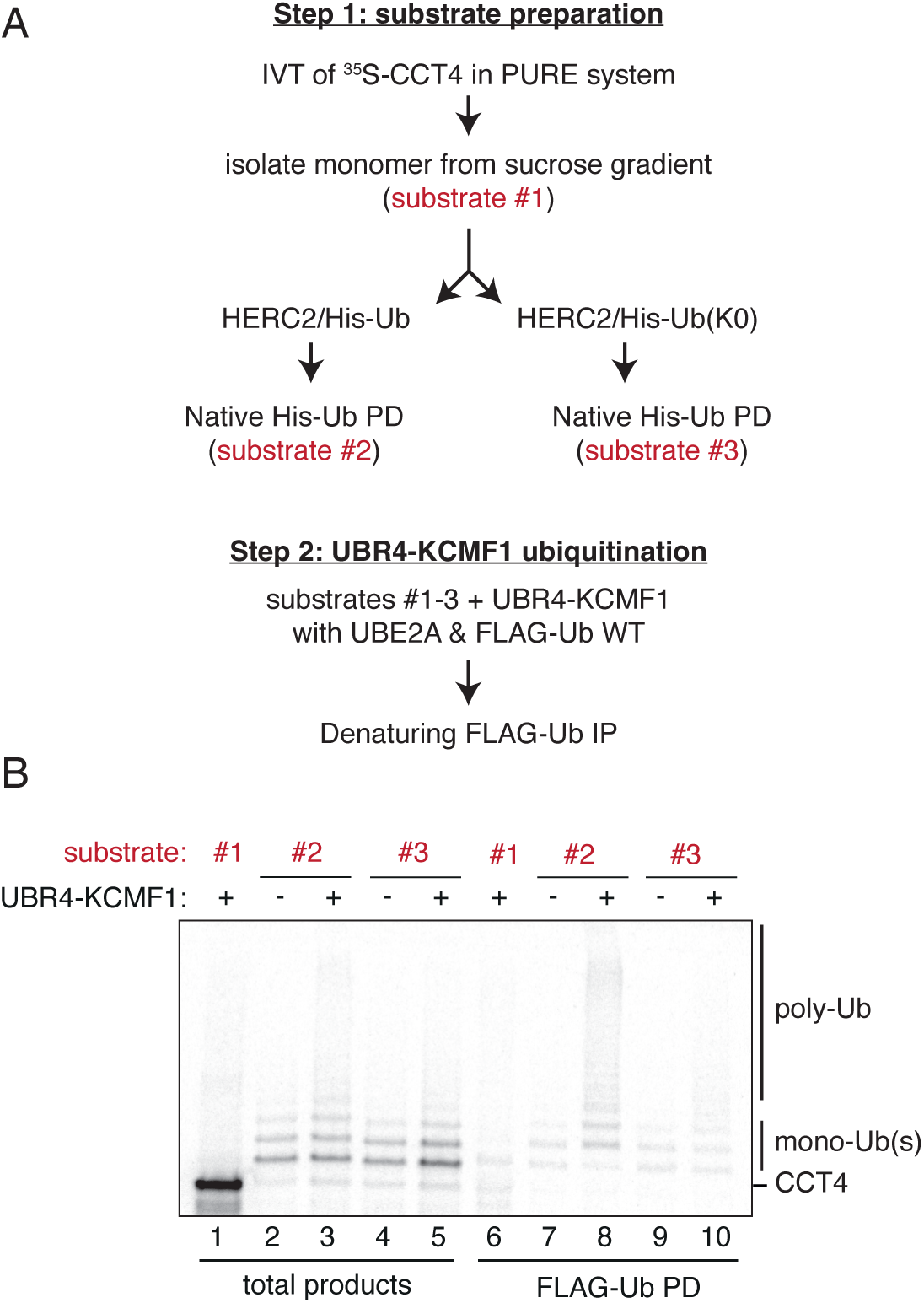
The UBR4-KCMF1 complex poly-ubiquitinates mono-ubiquitinated CCT4. **(A)** Schematic of experimental approach, indicating the steps from which the three ^35^S-methionine-labelled substrates for ubiquitination assays were taken. **(B)** ^35^S-methionine-labelled CCT4 substrates produced as outlined in A were incubated with El, E2 (UBE2A), wild type FLAG-Ub and recombinant UBR4 and KCMF1. The samples were then analyzed by autoradiography either directly (total products) or after a denaturing FLAG-Ub pulldown.

The finding that unmodified CCT4 was not an effective target for the UBR4-KCMF1 complex (lane 6) indicates that pre-ubiquitination by HERC2 strongly facilitates ubiquitination by the UBR4-KCMF1 complex. Importantly, the use of lysine-free (K0) ubiquitin in the HERC2 reaction did not appreciably impair HERC2-mediated multi-mono-ubiquitination (lane 2 vs. 4), but sharply reduced poly-ubiquitination by the UBR4-KCMF1 complex supplied with wild type ubiquitin (lane 8 vs. 10). Thus, the UBR4-KCMF1 complex selectively ubiquitinates mono-ubiquitinated CCT4 by adding chains on the pre-existing ubiquitin(s).

These reconstitution experiments suggest that a poly-ubiquitin chain, the most common proteasome degradation signal, cannot be formed on CCT4 by either the UBR4-KCMF1 complex or the HERC2-ZNRD2 complex alone. Only through the sequential action of these factors can CCT4 be poly-ubiquitinated, explaining why both are needed for efficient degradation of orphaned CCT4 in cells. A further implication of the findings is that mono-ubiquitin may be involved in substrate recognition by the UBR4-KCMF1 complex, allowing this substrate-linked ubiquitin to be positioned appropriately for chain formation, which evidently cannot occur effectively on any of CCT4’s numerous surface lysine residues. To begin exploring these mechanistic hypotheses, we turned to structural modelling.

### Structural modelling of the UBR4-KCMF1 complex

Using AlphaFold^42–44^, we generated a high-confidence structural model of a complex containing UBR4, KCMF1, UBE2A, and calmodulin (CALM), recently defined as the SIFI complex^35^ (Fig. S6A-C and Fig. 5A,B). As depicted in the schematic (Fig. 5B, right), the vast majority of UBR4 consists of a long snake-like alpha-helical scaffold consisting of extended armadillo repeats (grey) whose N-terminal and C-terminal ‘arms’ are likely to be flexible relative to the central part of the scaffold (Fig. 5B, left). UBE2A (cyan) is cradled by UBR4’s C-arm via a hemi-RING ligase domain^45^, CALM (brown) interacts with a flexibly-attached helix of the UBR4 scaffold, and KCMF1 (green) makes a bi-partite interaction with UBR4 via two separate domains (Fig. 5A,B).

**Figure 5.**
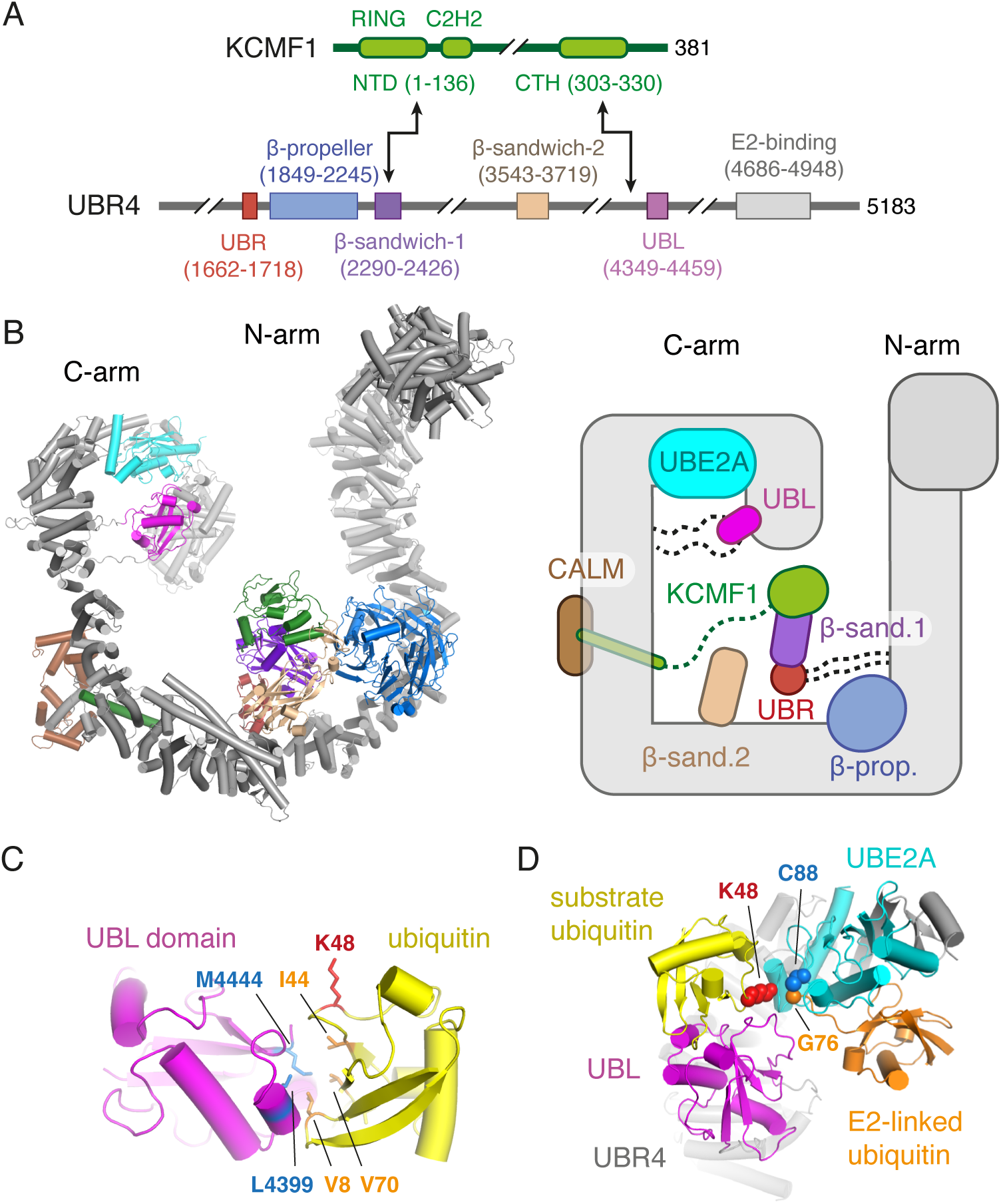
Structural model of ubiquitin recognition by the UBR4-KCMF1 complex. **(A)** Domain architecture of KCMF1 (top) and UBR4 (bottom) with boundaries based on structural modelling (see Fig. S6). Arrows indicate domain interactions between KCMF1 and UBR4. The diagrams are not to scale. **(B)** A composite AlphaFold3 structural model (left; see Fig. S6 for details) and schematic (right) of a complex comprising UBR4, KCMF1, CALM and UBE2A. Long unstructured loops of low confidence are not displayed in the structural model, but some of these are indicated by dotted lines in the schematic. The positions of the N-arm and C-arm are arbitrary, as they seem to be flexible and predicted in different relative positions in different models, hinging at roughly the region preceding the beta-propeller domain and downstream of the site of CALM binding. **(C)** Close-up view of the high-confidence predicted interaction between ubiquitin (yellow) and the UBL domain of UBR4 (magenta). Key hydrophobic residues in UBR4 (blue) that abut the hydrophobic patch of ubiquitin (orange), as well as lysine-48 (K48) of ubiquitin (red) are indicated. (D) Structural model of ubiquitin chain elongation by UBR4. The relative positions of the UBL domain of UBR4 (magenta), its bound ubiquitin (yellow), UBE2A with a charged ubiquitin, and a region of the UBR4 C-arm are shown. In this configuration, K48 (red spheres) of the UBL-bound ubiquitin is close to C88 of UBE2A (blue spheres) onto which ubiquitin is charged via its terminal glycine (G76). See also Fig. S6.

The alpha-helical scaffold of UBR4 is interspersed with five well-defined non-scaffold domains, some of which are attached via flexible linkers: UBR (red; C1662-A1718), β-propeller (blue; E1849-N2245), β-sandwich1 (purple; D2290-T2426), β-sandwich2 (tan; C3541-P3719), and UBL (magenta; F4349-L4459). The UBR and β-sandwich1 domains of UBR4 are confidently predicted to form a complex with each other and with the N-terminal domain (NTD) of KCMF1 (which contains the RING ligase domain) (Fig. S6B). This module is flexibly tethered to the UBR4 scaffold and is likely to be mobile. The UBL and β-sandwich2 domains are also flexibly tethered to the UBR4 scaffold (e.g., Fig. S6C), whereas the β-propeller seems to be more stably bound to the scaffold via an extensive conserved interface.

Because UBR4 prefers ubiquitination of pre-ubiquitinated substrates, we posited that it may contain a ubiquitin-interacting domain. Consistent with this idea, a single high-confidence ubiquitin binding site is predicted on UBR4 at its UBL domain (Fig. S6D; Fig. 5C). The UBL domain engages this putative substrate-linked ubiquitin via ubiquitin’s hydrophobic patch (L8, I44, V70), as is typical for ubiquitin-binding domains^46,47^. The long flexible linkers to which the UBL domain is attached (Fig. S6C) would allow it to readily reach UBE2A, bringing the substrate’s ubiquitin in proximity to the ubiquitin charged on UBE2A.

In the configurations predicted by AlphaFold3, K48 of the UBL-bound ubiquitin points toward C88 on UBE2A, onto which a second ubiquitin is charged (Fig. 5D). Notably, the ubiquitin that would be charged onto UBE2A (modelled using AF3 and consistent with various E2∼Ub structures) occupies a position that does not clash with any part of UBR4 or the substrate-ubiquitin bound to the UBL domain. The configuration modelled in Fig. 5D illustrates a plausible mechanism by which E2-charged ubiquitin is transferred to K48 of substrate ubiquitin. Although other configurations may be possible given the UBL domain’s flexible linkers, later experiments below indicate that K48 is indeed the predominant linkage type generated on orphaned clients by the UBR4-UBE2A complex.

### Structure-guided domain analysis of the UBR4-KCMF1 complex

The predicted structures provided a foundation for identifying key functional domains of the UBR4-KCMF1 complex. Focusing first on KCMF1, we unexpectedly found that the NTD (including the RING domain) and downstream linker region were completely dispensable for orphan CCT4-GFP degradation in cells (Fig. 6A). By contrast, the C-terminal domain (CTD), which contains a highly conserved helix (CTH: E303-L330) embedded in UBR4’s structural scaffold, was necessary and sufficient for restoring CCT4-GFP degradation in KCMF1 KO cells (Fig. 6A,B). Consistent with the predicted bi-partite interaction of KCMF1 with UBR4, all of these deletion mutants retained an interaction with UBR4 by co-IP, with only the ΔCTD showing a partial loss of interaction (Fig. S7A).

**Figure 6.**
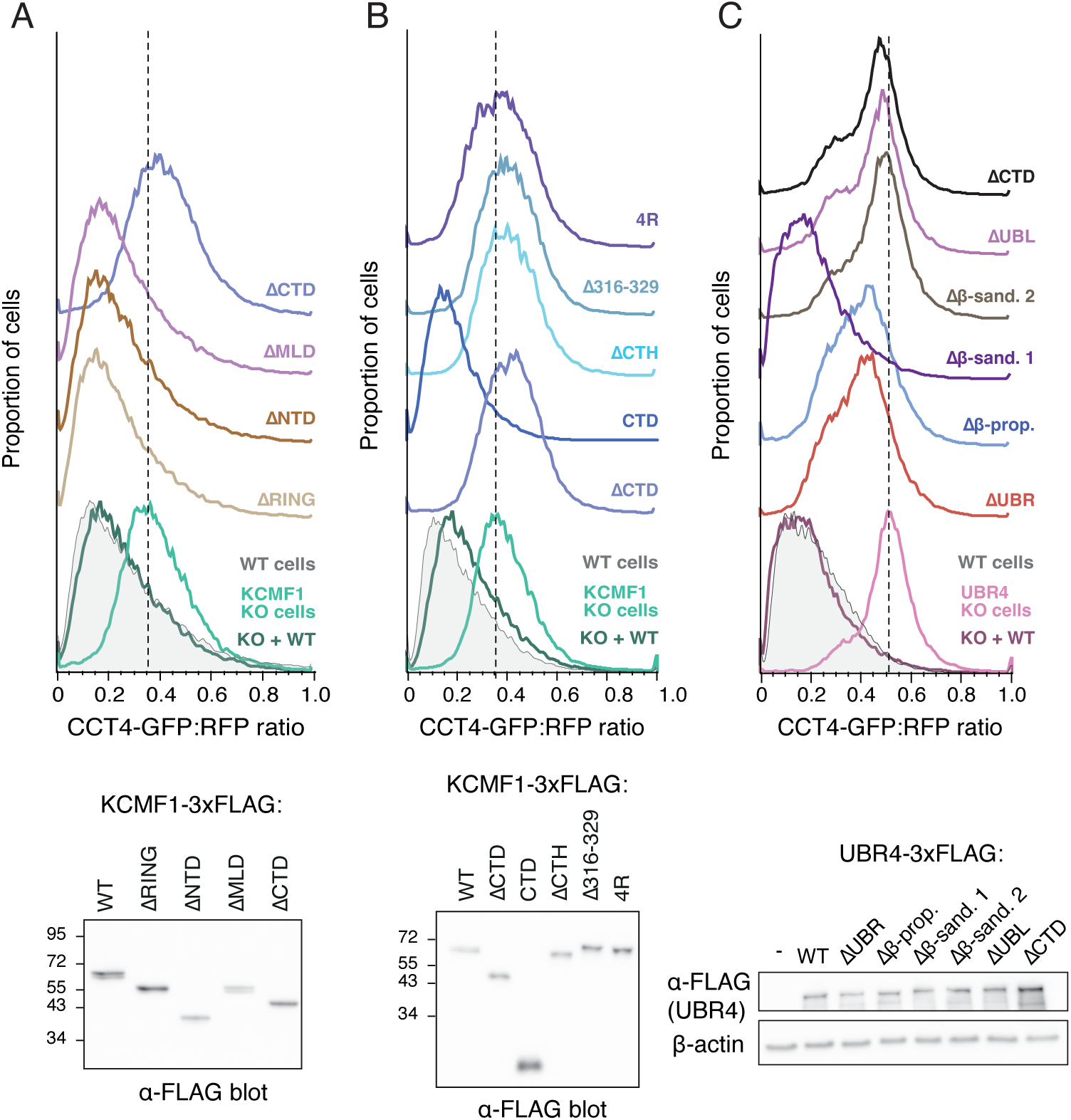
Identification of functional domains of the UBR4-KCMF1 complex. **(A)** Wildtype and AKCMF1 cells were transiently co-transfected with the CCT4 reporter and KCMF1 constructs (as indicated) for 48Һ and analyzed by flow cytometry (top) or immunoblot (bottom). NTD (N-terminal domain), MLD (middle-linker domain), CTD (C-terminal domain). **(B)** Wildtype and AKCMF1 cells were transiently co-transfected with the CCT4 reporter and KCMF1 constructs (as indicated) for 48Һ and analyzed by flow cytometry (top) or immunoblot (bottom). CTH (C-terminal helix), 4R (F319R/L323R/L324R/L325R). **(C)** Wildtype and AUBR4 cells were transiently co-transfected with the CCT4 reporter and UBR4 constructs (as indicated) for 48Һ and analyzed by flow cytometry (top) or immunoblot (bottom). See also Fig. S7.

Deletion of only the conserved CTH that complements UBR4’s scaffold was sufficient to completely inactivate KCMF1 without influencing expression of either KCMF1 or UBR4 (Fig. 6B; Fig. S7B). A similar effect was seen by mutating to arginine four conserved hydrophobic residues in this helix predicted to form key interactions with UBR4 (4R mutant; Fig. 6B). Thus, with respect to orphaned CCT4-GFP degradation, the critical role for KCMF1 is due entirely to a small structural element, the CTH, that buffets the UBR4 scaffold. Because deletion of the CTH does not impact UBR4 stability, its role may be to ensure appropriate positioning of the UBE2A-containing C-arm of UBR4 relative to other UBR4 modules that may participate in substrate engagement.

Consistent with no role for the NTD of KCMF1, deletion of β-sandwich1 on UBR4, with which it interacts, had no effect on CCT4-GFP degradation (Fig. 6C). By contrast, deletion of the UBE2A-interacting C-terminal domain (CTD) of UBR4 or the ubiquitin-interacting UBL domain completely abolished UBR4 function without an effect on expression levels. Deletion of β-sandwich2 also abolished CCT4-GFP degradation, whereas partial function was seen with deletion of the β-propeller or UBR domains, again without an effect on expression (Fig. 6C). A critical role for the UBL domain in orphaned CCT4 degradation, together with its predicted ubiquitin-binding capacity, suggests that ubiquitin recognition by the UBR4-KCMF1 complex plays a central role in its targeting of substrates for degradation.

### The UBR4-KCMF1 complex builds K48-linked poly-ubiquitin

To examine the role of substrate ubiquitin, we returned to the PURE *in vitro* system and used target proteins to which a non-cleavable ubiquitin (with the G76V mutation) was fused inline to the substrate’s N-terminus, thereby mimicking mono-ubiquitinated substrates without having to use a priming ligase (Fig. S8A). As was seen with HERC2-ubiquitinated CCT4, the Ub-CCT4 fusion protein isolated from the PURE system was preferentially poly-ubiquitinated by the UBR4-KCMF1 complex, whereas CCT4 was not (Fig. 7A). Similar results were seen for Ub-PSMB4 and even for Ub-GFP (Fig. 7A). That PSMB4 is not a target for ubiquitination by the UBR4-KCMF1 complex *in vitro* but is dependent on UBR4 and KCMF1 in cells suggests that a yet-unidentified priming ligase must first act on PSMB4. The Ub-GFP result indicates that although GFP is not an orphaned protein, the attached mono-ubiquitin is sufficient to target it for poly-ubiquitination by the UBR4-KCMF1 complex *in vitro*.

**Figure 7.**
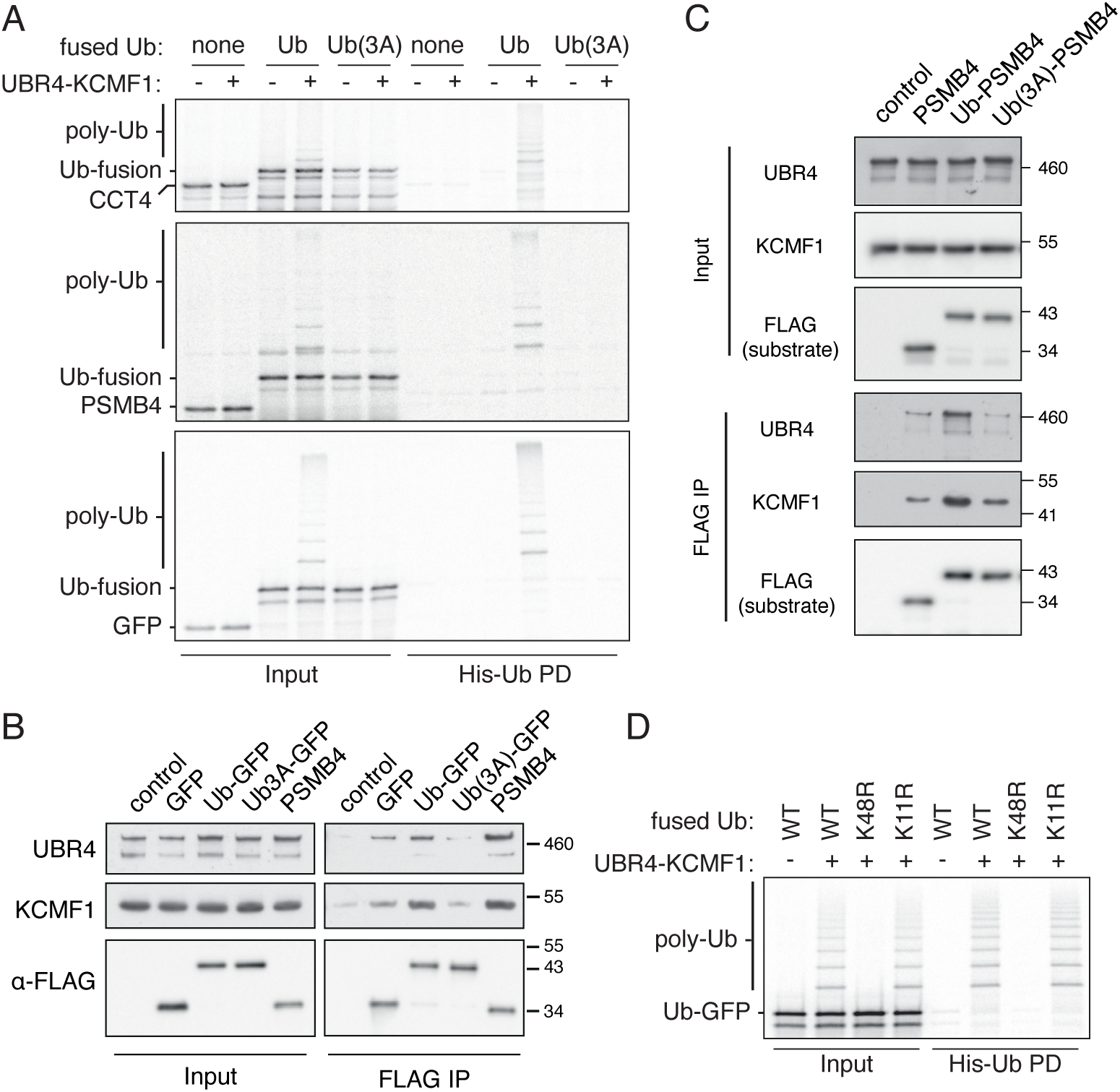
Mechanism of K48-polyubiquitination by the UBR4-KCMF1 complex. **(A)** ^35^S-methionine-labelled PSMB4, CCT4 and GFP were translated in the PURE system with or without N-terminal fusions of either ubiquitin (Ub) or ubiquitin containing a mutated hydrophobic patch [Ub(3A)]. The G76V mutation in ubiquitin was used in the fusion. Soluble substrates (see Fig. S8A) were then incubated with El, E2 (UBE2A), His-Ub, ATP and recombinant UBR4 and KCMF1. The samples were then analyzed by autoradiography either directly (Input) or following a His-Ub PD under denaturing conditions. **(B)** FLAG-tagged GFP, Ub-GFP, Ub(3A)-GFP and PSMB4 were translated in RRL and subjected to FLAG IP under native conditions. Input and IP samples were then analyzed by immunoblot. **(C)** FLAG-tagged PSMB4, Ub-PSMB4 and Ub(3A)-PSMB4 were translated in RRL in the presence of 20 pM El inhibitor (TAK-243) to prevent substrate ubiquitination and subjected to FLAG IP under native conditions. Input and IP samples were then analyzed by immunoblot. **(D)** ^35^S-methionine-labelled Ub-GFP, Ub(K48R)-GFP and Ub(Kl 1R)-GFP translated in the PURE system were incubated with El, E2 (UBE2A), His-Ub, ATP and recombinant UBR4 and KCMF1. The samples were then analyzed by autoradiography either directly (Input) of after His-Ub PD. See also Fig. S8.

Consistent with the structural model for substrate ubiquitin recognition, mutation of the hydrophobic patch (L8A, I44A, V70A) on the PURE system-produced Ub-fusion substrates completely abolished poly-ubiquitination by the UBR4-KCMF1 complex (Fig. 7A). Co-IP experiments using Ub-GFP translated in reticulocyte lysate showed that it interacts with endogenous UBR4-KCMF1 complex, and this interaction relies partially on the hydrophobic patch of ubiquitin (Fig. 7B). In the case of a bona fide orphan (e.g., PSMB4), an interaction comparable to Ub-GFP was seen even without a fused ubiquitin (Fig. 7B), although enhancement of the interaction was seen with Ub-PSMB4 (Fig. 7C). Despite an interaction similar to Ub-GFP, unmodified PMSB4 is not ubiquitinated by UBR4-KCMF1 (Fig. 7A). Instead, the substrate-linked ubiquitin seems to be crucial, evidently to provide a target site for poly-ubiquitination. Consistent with this idea, CCT4 engages UBR4-KCMF1 as effectively as Ub-CCT4 (Fig. S8B), even though only the latter is subject to poly-ubiquitination (Fig. 7A).

Using Ub-GFP produced in the PURE system, we found that the major site of poly-ubiquitination is K48, since mutation of only this lysine to arginine in Ub-GFP completely abolished all ubiquitination by the UBR4-KCMF1 complex (Fig. 7D). When ubiquitin(K48R) was used in the reaction with wild type Ub-GFP, the UBR4-KCMF1 complex could add the first ubiquitin effectively, but poly-ubiquitination was markedly reduced to the same levels as K0 ubiquitin (Fig. S8C). No other lysine mutant of ubiquitin showed this effect, indicating that in this system, the poly-ubiquitin generated by the UBR4-KCMF1 complex is essentially all K48-linked. Similarly, use of ubiquitin(K48R) in reactions with Ub-CCT4 and Ub-PSMB4 permitted the attachment of the first ubiquitin, but not effective elongation of poly-ubiquitin chains (Fig. S8D,E).

## Discussion

We have discovered that multiple unrelated orphaned proteins rely on the UBR4-KCMF1 complex. Focusing primarily on the orphaned CCT subunit pathway involving the HERC2-ZNRD2 complex, we revealed a two-step mechanism for generating a proteasome degradation signal on the orphan. The first step involves ‘priming’ with one or more mono-ubiquitins. For CCT4, PSMC5 and UBL4A, the priming ligases are likely to be HERC2, HERC1, and HUWE1, respectively^12,20,37^, whereas the priming ligase for PSMB4 remains to be identified. The second step involves substrate- and ubiquitin recognition by the UBR4-KCMF1 complex. This step builds a K48-linked poly-ubiquitin chain onto K48 of substrate ubiquitin, resulting in a suitable signal for proteasome targeting^40^.

The mechanistic basis for ubiquitin recognition seems to involve the UBL domain of UBR4. Because the UBL domain and the C-arm where UBE2A resides are both flexible, various positions of ubiquitin on substrates of different sizes and shapes could be accommodated. Although the site(s) of substrate binding on the UBR4-KCMF1 complex are currently unknown, they may involve the UBR, β-sandwich or β-propeller domains. Each of these are typical protein-protein interaction elements, and having multiple such domains could allow the UBR4-KCMF1 complex to interact with a range of orphaned substrates. Notably, the β-sandwiches and UBR domain seem to be flexibly tethered to the UBR4 scaffold, which would help accommodate diversity in substrate shape and size. One prediction of this model is that different substrates would rely on different domains, an idea that merits future work.

The capacity of the UBR4-KCMF1 complex to accommodate different unrelated mono-ubiquitinated substrates means that multiple AQC pathways converge at this ligase complex for efficient degradation. Given the high orphan burden in cancer cells, points of convergence such as this may represent particularly potent vulnerabilities. Consistent with this idea, both KCMF1 and UBR4 were required for the survival of highly aneuploid cancer cell lines^33^, with UBR4 essentiality correlating strongly with copy number imbalances among protein complex subunits^34^. Thus, a major role of the UBR4-KCMF1 complex is in maintaining cellular proteostasis via degradation of orphans, a large source of quality control substrates in cells.

For orphaned CCT4-GFP, a role for the putative ligase activity of KCMF1 was dispensable, with only a C-terminal structural helix being essential. It is possible that other substrates, perhaps those whose shape or size cannot access the UBR4 C-arm effectively, would rely instead on KCMF1 for ubiquitination. Alternatively, some substrates may rely on branched ubiquitin chains for effective degradation^48^. In this case, it is possible that having two ligases would allow for branched chain formation, as has been suggested for the UBR4-KCMF1 complex^49^. Exploration of this idea will first require identifying a substrate whose degradation relies critically on both the UBR4 and KCMF1 ligase domains.

As with KCMF1, the role of CALM, if any, for orphaned protein degradation remains poorly understood. An earlier suggestion that KCMF1’s ligase domain and CALM are critical for function of the SIFI complex^35^ should be interpreted with some caution because the deletion mutants intended to disrupt CALM and KCMF1 binding included key structural elements of the UBR4 scaffold (Fig. S6E). If CALM were to be involved, it may serve a structural role analogous to that provided by the C-terminal helix of KCMF1. CALM seems unlikely to recruit substrate given that CALM’s key substrate-interacting regions are already engaged with UBR4. An improved structural model, CALM inhibitors, and a recombinant *in vitro* system for orphan ubiquitination should help address this issue.

The relationship between our findings and a recently proposed role for the UBR4-KCMF1 complex in dampening the integrated stress response (ISR) after mitochondrial stress remains to be resolved^35^. The targets of ISR silencing are the ISR-initiating ligase HRI, the mitochondrial stress transducer DELE1, and non-imported mitochondrial precursors. Whether any or all of these substrates require separate priming and elongation steps, as observed here for orphaned CCT4, is unknown. However, a two-step process seems plausible given that the UBR4-KCMF1 complex uses its UBL domain to position a pre-existing substrate ubiquitin for K48 modification, facilitating chain formation.

In this context, it is noteworthy that non-imported ER and mitochondrial precursors engage ligase-associating QC factors that seem to add only one or a few ubiquitins^50–52^. HRI and DELE1 may similarly be primed by a yet-unknown ligase. A parsimonious model would be one where stress silencing, AQC, and perhaps other pathways converge at the UBR4-KCMF1 complex for creating a poly-ubiquitin signal that commits substrates for degradation. Such a model would explain the multi-faceted proteostasis phenotypes of losing this ligase complex, including high sensitivity to aneuploidy^33^, prolonged stress signalling^35^, and reduced efficiencies of seemingly unrelated degradation pathways like ERAD^53^ and muscle protein degradation^54^.

A two-step mechanism for AQC and stress silencing may allow for reversibility and regulation. For example, a priming ubiquitin may be a provisional mark subject to reversal, thereby allowing greater time for protein complex assembly, stress signalling, or mitochondrial import. Commitment to degradation would only occur after a poly-ubiquitin signal has been built and has engaged downstream poly-ubiquitin-binding receptors such as the p97 ATPase^55,56^ or the proteasome cap^57^. Such a multistep process would allow the threshold for AQC or stress silencing to be set differently under different conditions or in different cell types.

## Key resources table

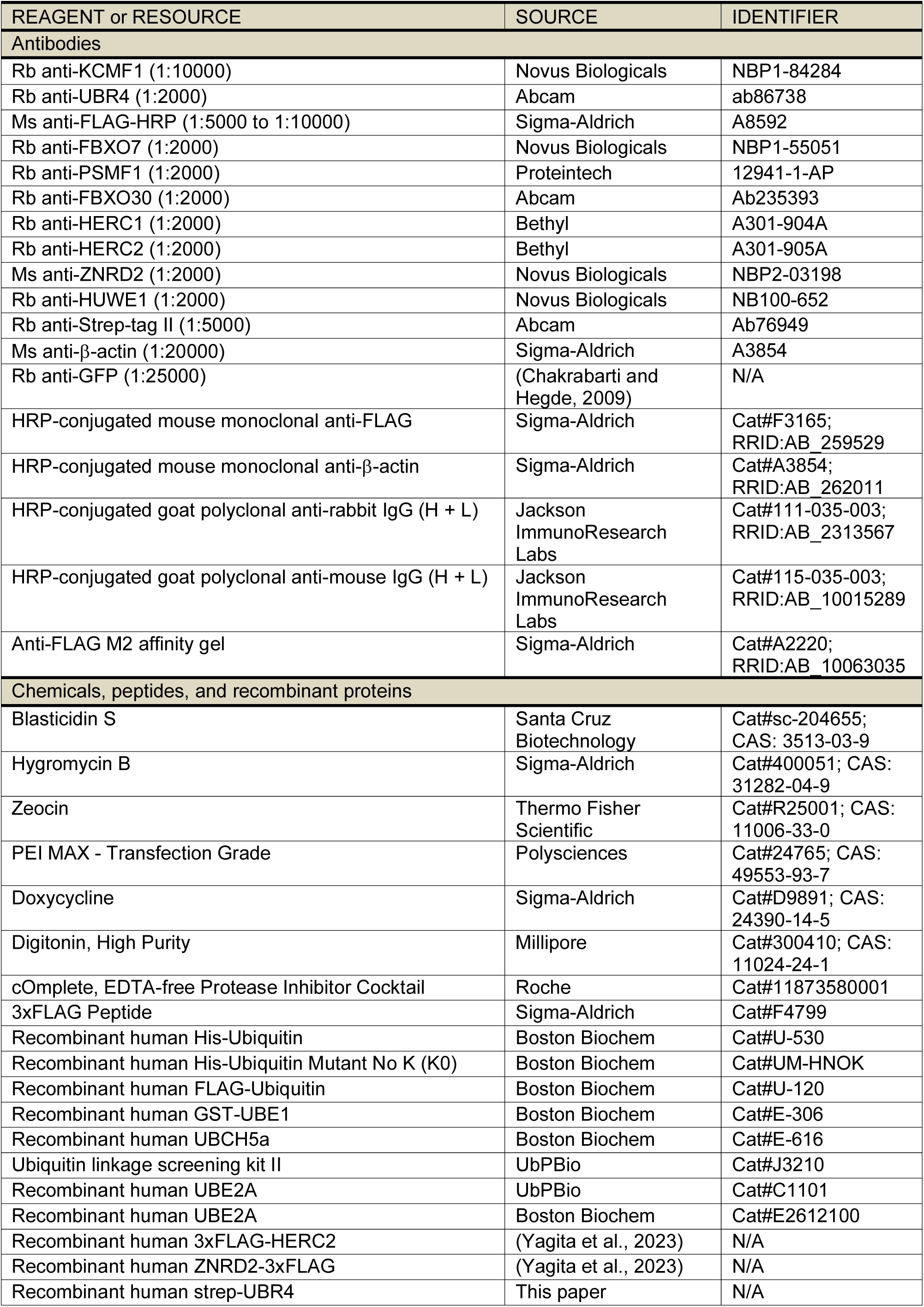
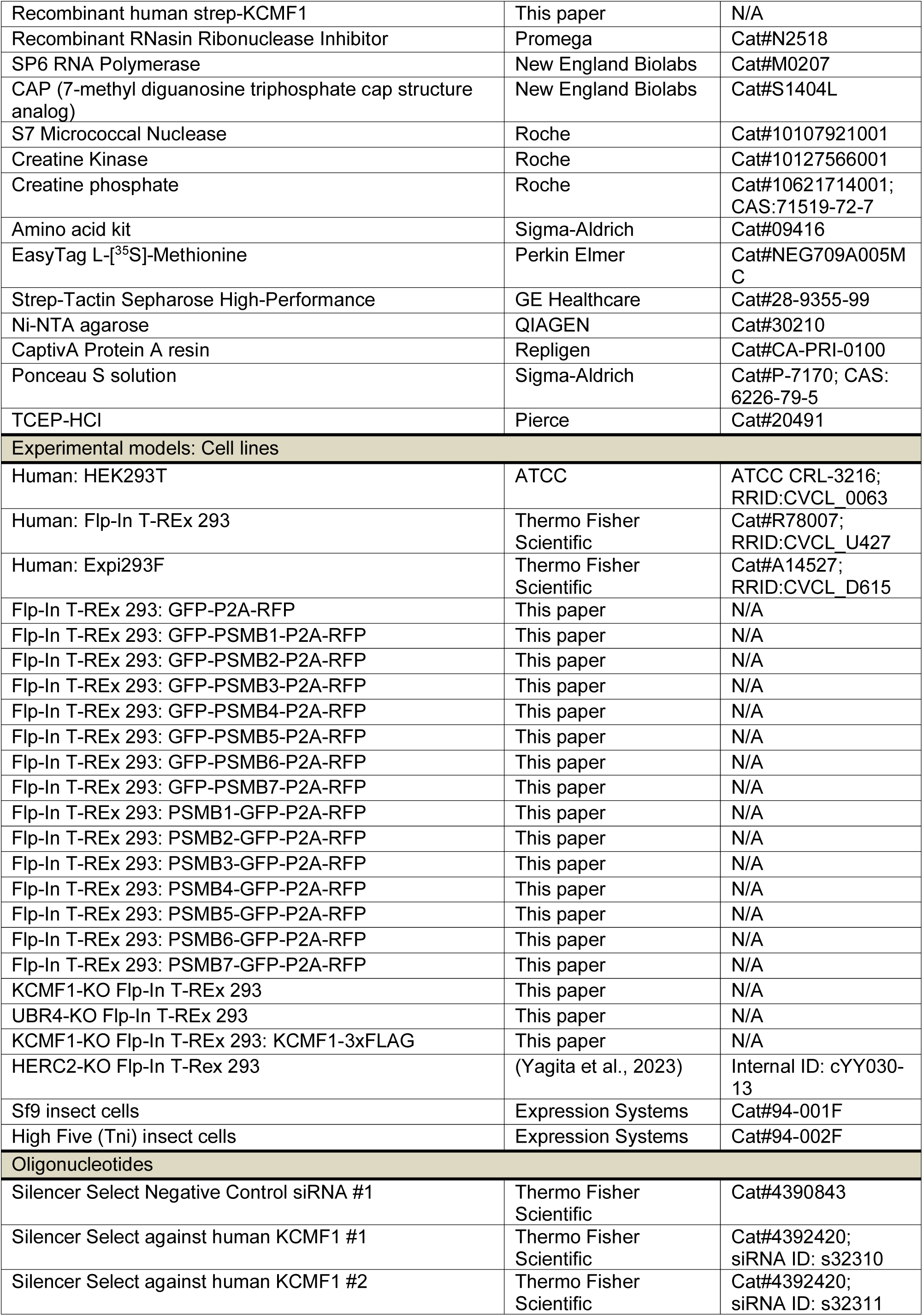
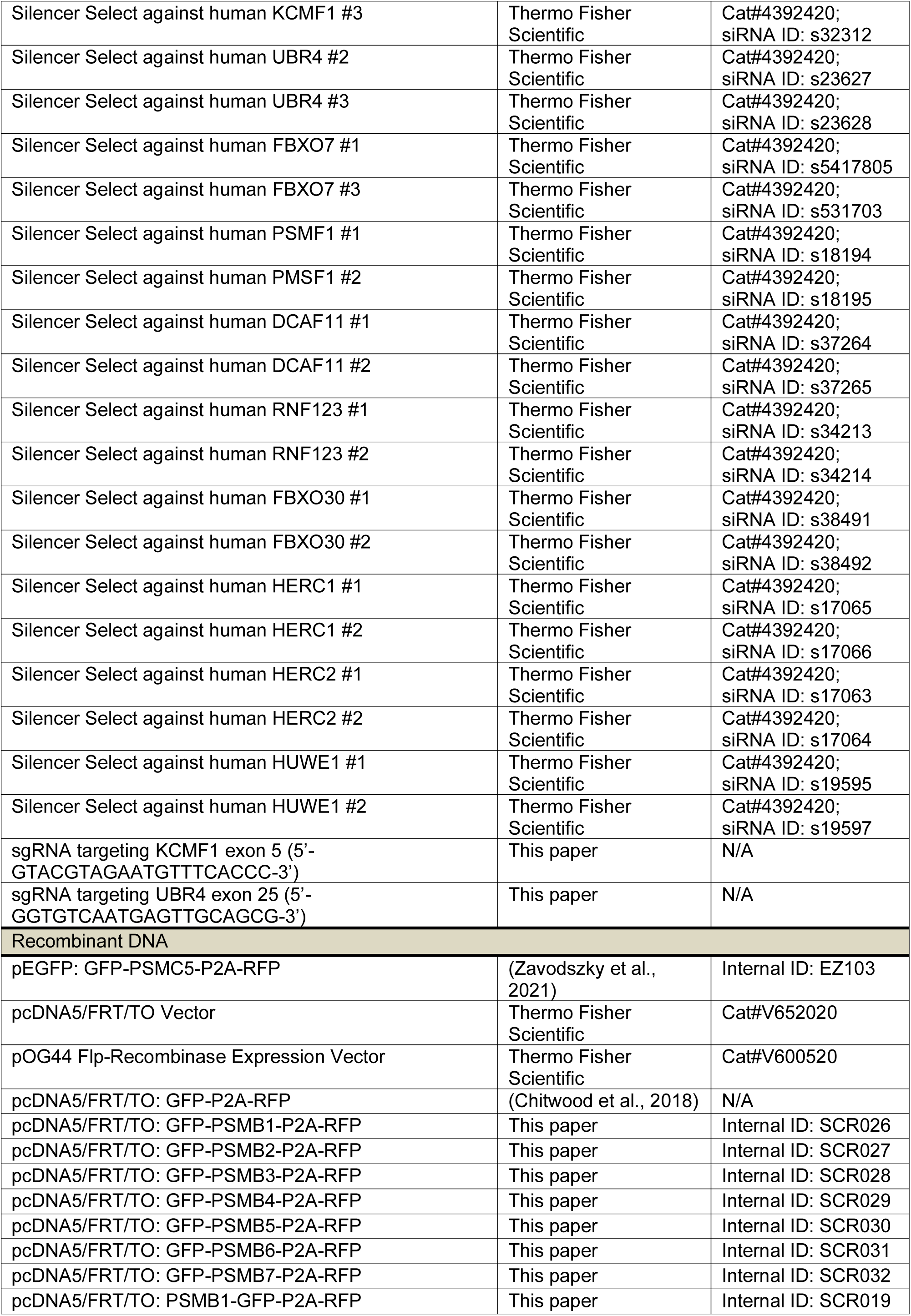
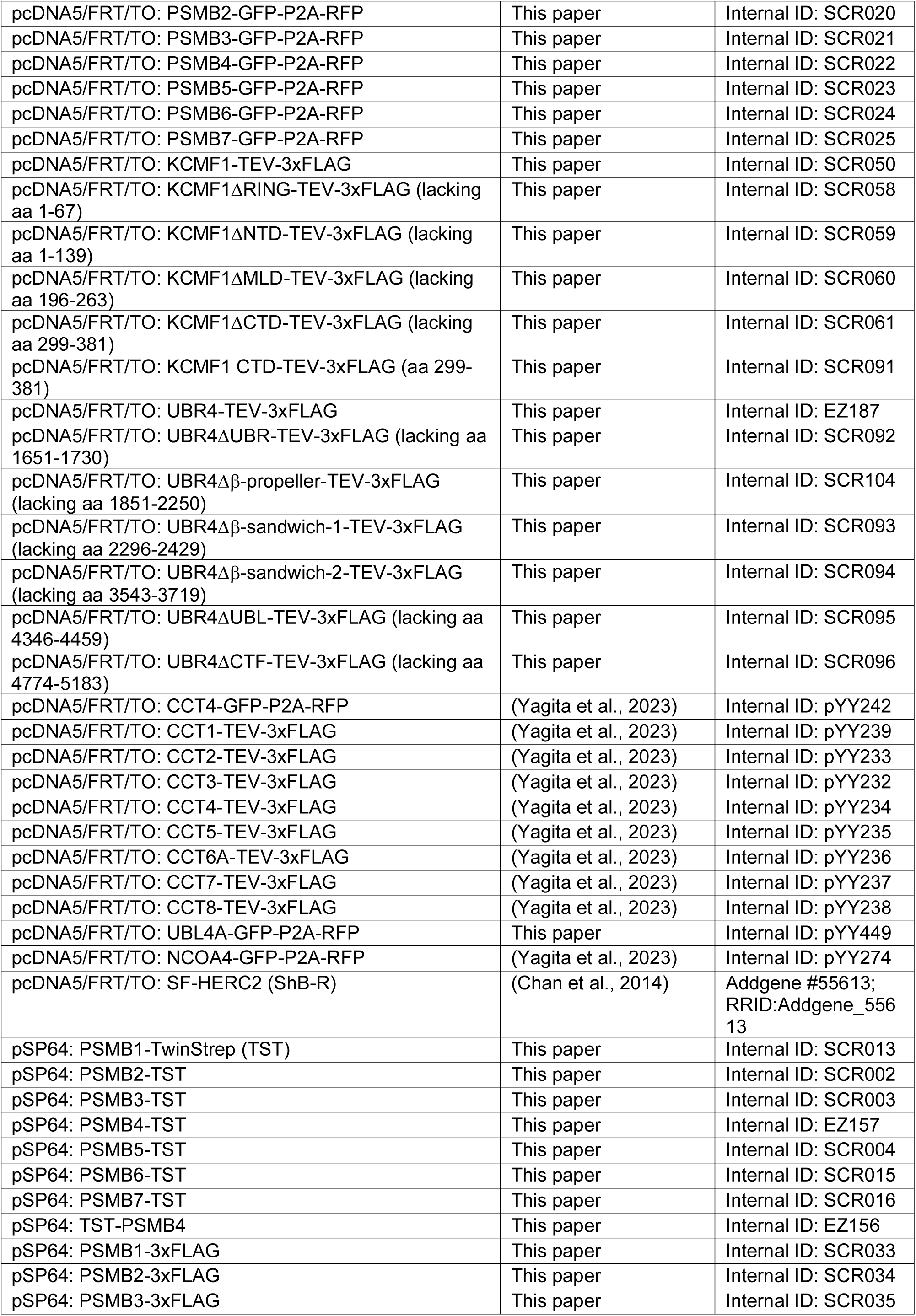
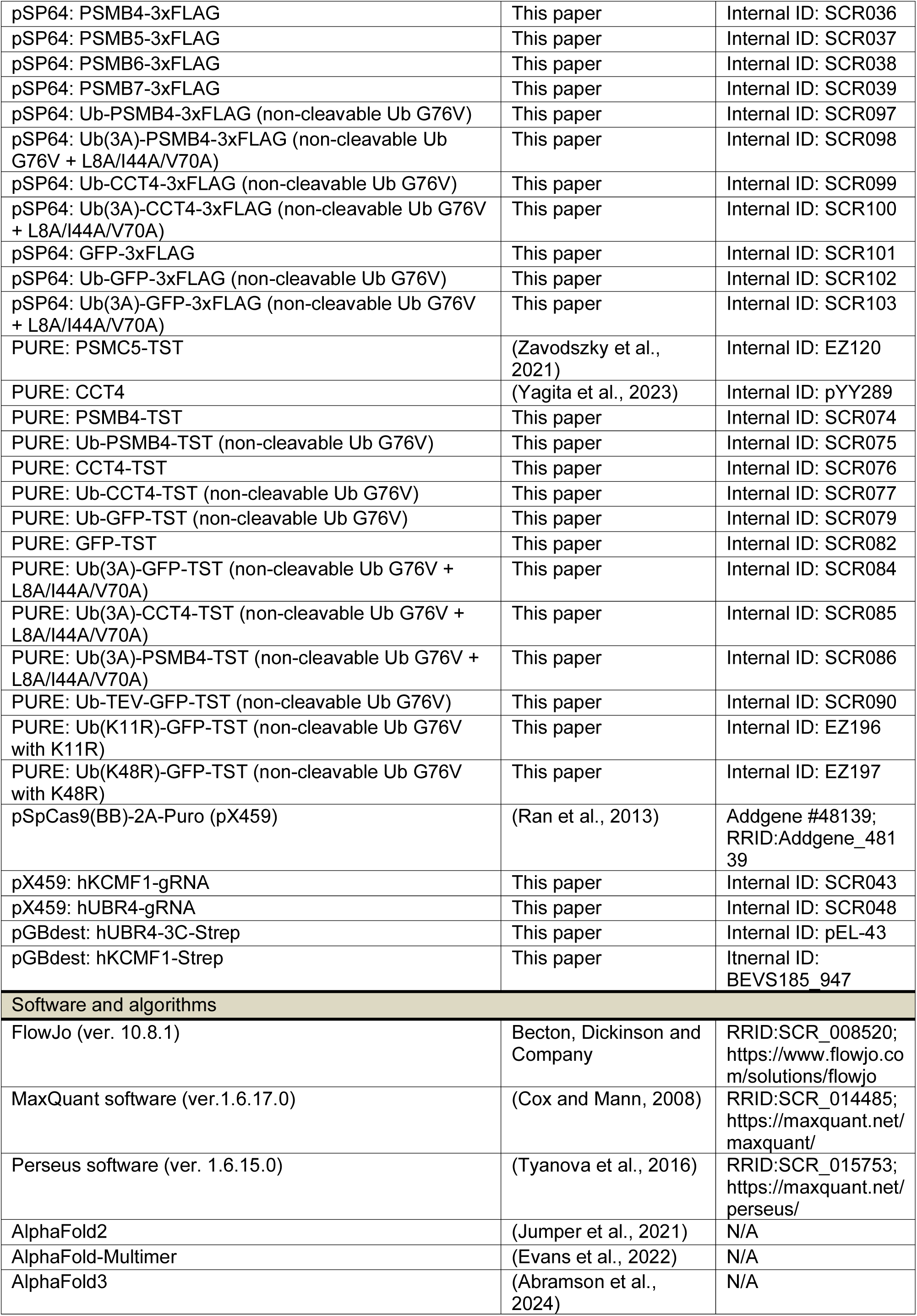
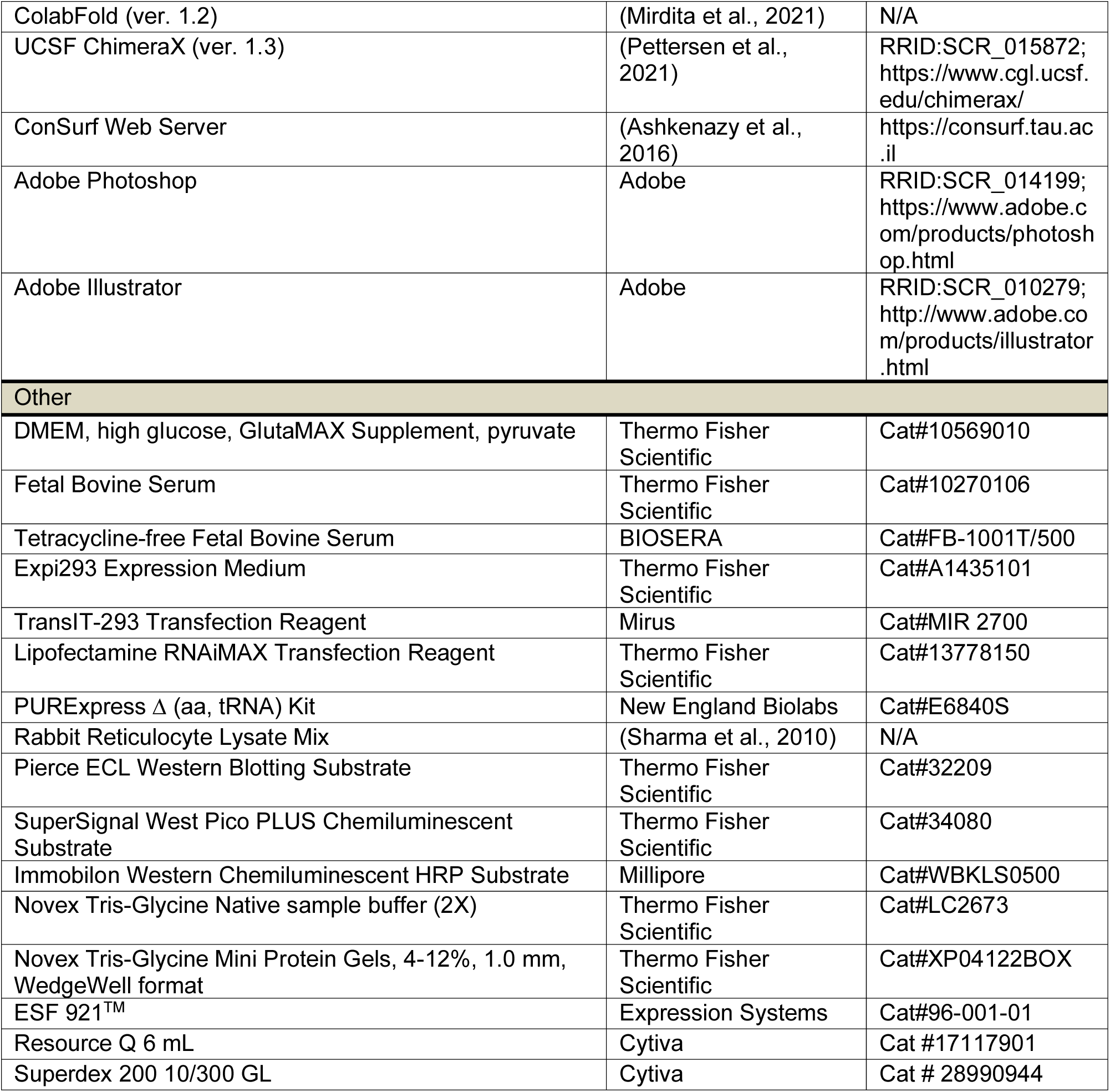

## Materials and methods

### Constructs

All plasmids were constructed using standard molecular cloning techniques, sequenced, and listed in the Key Resources Table (KRT). Constructs for *in vitro* translation in rabbit reticulocyte lysate (RRL) were cloned into pSP64 with an N-terminal or C-terminal twin-strep tag (TST) or C-terminal 3xFLAG tag. Constructs for *in vitro* translation in the PURE system were cloned into the T7-based PURExpress plasmid from New England Biolabs with a C-terminal TST tag. Constructs for mammalian expression were cloned into pcDNA3.1 (Thermo Fisher) or pcDNA5/FRT/TO (Thermo Fisher). The fluorescent reporter constructs were based either on pEGFP or pcDNA5/FRT/TO vectors containing a EGFP-P2A-mCherry cassette. For simplicity, EGFP and mCherry are referred to throughout the paper as GFP and RFP, respectively. KCMF1-FLAG was obtained from GenScript (ohu31023) and the KCMF1 coding region was sub-cloned into a pcDNA5/FRT/TO vector with a TEV cleavage site and a C-terminal 3xFLAG tag. UBR4 for mammalian expression was amplified from HEK293T cDNA and cloned into pcDNA5/FRT/TO with a C-terminal 3xFLAG tag. KCMF1 and UBR4 mutants were then generated by mutagenesis of the respective wild type plasmids. Full length human UBR4 (with a C-terminal 3C site and a STREP-tag) and human KCMF1 (with a C-terminal STREP-tag) were cloned into pGBdest vector using synthetic gene fragments and the GoldenBac assembly system^58^.

### siRNAs and sgRNAs

Pre-designed Silencer Select siRNAs (listed in the KRT) were obtained from Thermo Fisher and resuspended in water to 20 µM. For nontargeting control conditions, negative control siRNA No.1 (Cat#4390843) was used. To make Flp-in TREX KCMF1 and UBR4 KO cells using CRISPR, the guide RNAs listed in the KRT were cloned into the pX459 vector.

### Antibodies

All antibodies used in this study are listed in the KRT with dilutions used for immunoblot.

### Recombinant proteins

Full length human UBR4 with C-terminal strep and strep-tagged human KCMF1 were expressed in High Five cells and purified as follows. Cells were infected at a density of 1,5x 106 cells/ml with UBR4 or KCMF1 viral stock (1:75) and cultured at 27⁰C for 72 hours. Cells were harvested at 600g for 15 minutes, resuspended in 1x PBS and pelleted again at 700g for 10 minutes. Cell pellets were flash frozen in liquid nitrogen and stored in -70 freezer prior to purification. Pellets from UBR4-3C-Strep expression were resuspended in Lysis buffer containing 1x PBS, 0.5mM TCEP, protease inhibitor (EDTA-free cOmplete, Roche) and Benzonase (1:500, 2mg/ml). Cells suspension was lysed using a douncer (Kontes) and centrifuged at 43.000g for 40 minutes (Lynx 6000, Thermo Scientific) to clear the lysate. The protein was purified by Strep-Tactin affinity chromatography using a StrepTrap HP 5ml (Cytiva) column and eluted with 2.5 mM desthiobiotin in the lysis buffer. Subsequently, an anion exchange chromatography step was performed on a Resource Q column (gradient 0 to 1M NaCl in 1x PBS, 0.5mM TCEP). Finally, the protein was concentrated using Vivaspin column (Sartorius). Pellets from KCMF1-Strep expression were resuspended in 50mM Hepes/KOH pH7.5, 500mM KCl, 0.5mM TCEP. The cell lysate was subjected to Strep-Tactin affinity and the KCMF1-containing fractions pooled, concentrated and applied to a Superdex 200 column (Cytiva) to separate remaining impurities by size-exclusion chromatography. Recombinant human HERC2 and ZNRD2 were purified from Expi293F cells as described previously^12,20^. Human His-ubiquitin (WT, K0 and K48R), GST-UBE1, UBE2D1 and UBE2A used for *in vitro* ubiquitination reactions were purchased from Boston Biochem. UBE2A was also purchased from UBPBio (C1101). Other ubiquitin Lys-to-Arg mutants were obtained from the ubiquitin linkage screening kit II from UBPBio (J3210).

### Cell culture

HEK293T and Flp-In T-REx 293 cells were maintained in 10% FBS in DMEM at 37°C with 5% CO_2_. Expi293F cells were grown in Expi293 Expression Medium at 37°C with 8% CO_2_. Flp-In T-REx 293 stable cells were supplemented with 10 µg/mL Blasticidin and 100 µg/mL Hygromycin B. Expression of the integrated construct was induced with 10 ng/µL of doxycycline for 18-48 h depending on the experiment. Where indicated, cells were treated with 100 µg/mL of cycloheximide (CHX), 20 µM of MG132 and/or 1 µM of MLN4924 for 1-6 h. Plasmid transfections were carried out using TransIT-293 transfection reagent (Mirus) according to the manufacturer’s instructions. For siRNA-mediated knockdown, cells were transfected with siRNAs using Lipofectamine RNAiMAX (Thermo Fisher Scientific) and analyzed 72-96 h later.

KCMF1 and UBR4 KO Flp-in T-REx 293 cell lines were generated by CRISPR/Cas9-mediated gene disruption as described previously^59^. Briefly, parental Flp-in T-REx cells were transfected with pX459 containing the guide RNA against KCMF1 or UBR4, which had been selected using the “ChopChop” and “CRSPick” website tools. After 24 h, transfected cells were selected with 2 µg/mL puromycin for another 48 h. Cells were then plated into 96-well plates at a density of 0.5 cells/well to isolate single cell clones. After 2 weeks, single-cell colonies were screened by immunoblotting for successful KO. Relevant phenotypes were verified in multiple independent clones to exclude clone-specific effects.

Flp-In T-REx 293 stable cell lines containing a doxycycline-inducible construct were generated using the Flp-In system (Thermo Fisher Scientific). In brief, cells were co-transfected with the appropriate pcDNA5/FRT/TO-based construct and Flp recombinase (pOG44) in a 1:9 ratio. After 48 h, cells were passaged into media containing 100 µg/mL Hygromycin B and 10 µg/mL Blasticidin S for approximately 10 days to select for stable integration of the plasmid at the FRT site. Expression of the inserted protein was tested by immunoblotting after Dox induction.

### Mammalian in vitro transcription and translation

In vitro transcription by SP6 RNA polymerase was performed as previously described^60^. In brief, 10 ng/µL PCR product served as template for transcription reactions in 40 mM HEPES pH 7.4, 6 mM MgCl_2_, 10 mM reduced glutathione, 20 mM spermidine, 0.1 mM GTP, 0.5 mM ATP, 0.5 mM CAP structure (NEB), 0.4-0.8 U/µL RNAsin (Promega) and 0.4 U/µL SP6 polymerase (Promega). Transcription reactions were incubated at 37°C for 1h and were used for *in vitro* translation reactions without further purification. *In vitro* translations in rabbit reticulocyte lysate (RRL) were performed as described previously^60,61^. Briefly, crude reticulocyte lysate (Green Hectares) was pre-treated with micrococcal nuclease to digest endogenous mRNAs. The RRL was then supplemented with 20 µg/mL total liver tRNA, 40 µM of each amino acid (except methionine), 1 mM ATP, 1 mM GTP, 12 mM creatine phosphate, 40 µg/mL creatine kinase, 1 mM reduced glutathione, 0.3 mM spermidine, and 20 mM HEPES pH 7.4, 10 mM KOH, 50 mM KAc, and 2 mM MgAc_2_. *In vitro* translations were initiated by adding the transcript (at 5% total volume) and either 40 µM unlabelled methionine or 0.5 µCi/µL ^35^S-methionine and incubated at 32°C for 30-60 min. Where indicated, translation was done in the presence of 20 µM TAK-243 E1 inhibitor. In experiments where the ubiquitinated products were affinity-purified, 10 µM of His-ubiquitin was added in the *in vitro* translation reaction. Immediately after translation, samples were placed on ice and any further manipulations were conducted at 0-4°C.

### Affinity purification of in vitro translation products

*In vitro* translation reactions were diluted in physiological salt buffer (PSB: 50 mM HEPES, 100 mM KAc, 2 mM MgAc_2_) at 4°C to a total volume of 1 mL. For TST-tagged or FLAG-tagged PSMB subunits, samples were incubated with 10-15 µL of streptactin Sepharose (Iba Life Sciences) or 15 µL of anti-FLAG M2 affinity gel (Sigma-Aldrich), respectively, for 60 min at 4°C. The beads were then washed 5 times in PSB at 4°C and transferred to a new tube with the 4^th^ wash. Streptactin-bound proteins were either eluted with 50 mM Biotin by incubating on ice for 20 min, or by heating to 95°C in 2X SDS-PAGE sample buffer (100 mM Tris, 2% SDS, 20% glycerol and 20 mM DTT) supplemented with 2 mM biotin. FLAG-tagged proteins were eluted by heating to 95°C in SDS-PAGE sample buffer. For identification of QC candidates by mass spectrometry, the beads containing immunoprecipitated proteins were processed directly as described below.

### Denaturing ubiquitin pulldowns

Ubiquitinated products were recovered through affinity purification of His- or FLAG-tagged Ubiquitin as indicated in the figure legends. Samples were denatured by boiling in 100 mM Tris pH 8.0 with 1% SDS, then diluted 10-fold in pulldown buffer (1x Phosphate Buffer Saline (PBS), 250 mM NaCl, 0.5% Triton X-100, 20 mM Imidazole) and incubated with 10 µL of Ni-NTA agarose (Qiagen) or 10 µL of anti-FLAG M2 affinity gel at 4°C for 1.5 h or overnight. The resin was washed 3 times with pulldown buffer and samples were eluted in SDS sample buffer, supplemented with 50 mM EDTA for Ni-NTA agarose pulldowns.

### Label-free quantitative mass spectrometry

*In vitro* translation and native immunoprecipitation were performed as described above. TST-tagged PSMB subunit constructs (PSMB1-7) served as bait proteins to identify interaction partners whereas a mock translation lacking transcript was used as a negative control. Affinity-purified protein samples on beads were resuspended in 50 µL of 20 mM HEPES, reduced in 10 mM DTT at 56°C for 30 min, and alkylated with 15 mM iodoacetamide in the dark at 22°C. After quenching the alkylation reaction with excess DTT, the samples were then digested with 200 ng of trypsin (Promega) overnight at 37°C. After centrifugation at 10,000 x g for 5 minutes, the supernatant was transferred to a new tube. The beads were washed once with 30 mL of 5% formic acid (FA), and the solution was combined with the corresponding supernatant. The resulting peptide mixtures were desalted using a home-made C18 (3M Empore) stage tip that contained 2 mL of Poros Oligo R3 resin (Thermo Fisher Scientific). Bound peptides were eluted from the stage tip with 30–80% acetonitrile (MeCN) and partially dried in a SpeedVac (Savant). Peptides were separated on an Ultimate 3000 RSLC nano System (Thermo Scientific) fitted with a 75 mm x 25 cm nanoEase C18 T3 column (Waters), using mobile phases buffer A (2% MeCN, 0.1% FA) and buffer B (80% MeCN, 0.1% FA). Eluted peptides were introduced directly via a nanospray ion source into a Q Exactive Plus hybrid quadrupole-Orbitrap mass spectrometer (Thermo Fisher Scientific). The mass spectrometer was operated in data dependent mode. MS1 spectra were acquired from 380–1600 m/z at a resolution of 70K, followed by MS2 acquisitions of the 15 most intense ions with a resolution of 17.5K and NCE of 27%. MS target values of 1e6 and MS2 target values of 5e4 were used. Dynamic exclusion was set for 40 sec. Each condition was analyzed three times to have technical triplicates for statistical analysis. The raw data files were processed for protein identification and quantification with MaxQuant software (version 1.6.17.0)^62^ employing the Andromeda search engine^63^. The data was searched against the *Oryctolagus cuniculus* UniProt FASTA database. Protein quantification was performed using the label-free quantitation (LFQ) algorithm in MaxQuant. MaxQuant output was further processed with Perseus software (version 1.5)^64^. Briefly, potential contaminants, reverse hits, hits only identified by site, and hits with only one unique and razor peptide were filtered out prior to log_2_ transformation of the LFQ intensities. Triplicates were grouped, and the data was filtered to keep proteins with three valid values in at least one group. In order to identify specific interactors of each bait protein, statistical analyses were performed using a two-tailed Student’s *t*-test with Benjamini-Hochberg correction for multiple comparisons. Volcano plots were generated using Prism software.

### Flow cytometry

Cells were transiently transfected with fluorescent reporters for 24-48 h prior to flow cytometry analysis. Where indicated, KCMF1 and UBR4 constructs were co-transfected with the reporters. A total of 1 µg DNA/well was transfected in 6 well plates, whereas 500 ng DNA/well was transfected in 12-well plates. Co-transfection used a 2:2:6 ratio for KCMF1:reporter:empty vector, and an 8:2 ratio for UBR4:reporter. In Flp-In T-REx 293 and its derivative cell lines, expression of stably and transiently transfected constructs was induced with 10 ng/µL doxycycline for 18-24 h prior to flow cytometry analysis. To prepare cells for flow cytometry, cells were washed once with PBS and either resuspended directly or first trypsinized and resuspended in ice-cold PBS containing 10% FBS. Cells were then pelleted by centrifugation and resuspended in ice-cold PBS containing 10% FBS and 1 µg/mL DAPI as a viability marker, prior to passage through a 70-µm cell strainer to ensure a single-cell suspension. Data were collected using a Beckton Dickinson LSRII or LSRFortessa flow cytometer, and subsequently analyzed using FlowJo software. At least 10,000 transfected cells were analyzed, with the majority of experiments containing ∼30,000 transfected cells. Each experiment is internally controlled, and histograms between independent experiments are not directly comparable, as fluorescence intensity values depend on the cytometer model, settings and calibration. In general, however, we used settings such that the control reporter containing untagged fluorophores generated approximately equal GFP and RFP fluorescence intensity values that fell on a diagonal line across a wide range of intensities (see Fig. 1B). Where parallel immunoblotting analysis of cells was desired, an aliquot of cells was harvested and lysed by boiling in 100 mM Tris-HCl pH 8.0, 1% SDS buffer for ∼10 min with occasional vortex mixing to shear genomic DNA.

### Immunoprecipitation from cells

To analyze interactions between 3xFLAG-tagged PSMB subunits and endogenous proteins, HEK293T cells in 6-well plates were transiently transfected with 1 µg of PSMB-3xFLAG constructs using TransIT-293 transfection reagent (Mirus). To analyze interactions between KCMF1-3xFLAG mutants and endogenous proteins, KCMF1 KO TREx 293 cells in 6-well plates were transiently transfected with 500 ng of KCMF1 plasmid supplemented with 500 ng of empty vector, and expression of KCMF1-3xFLAG was induced with 10 ng/µL doxycycline for 18-24h. To analyze interactions between UBR4-3xFLAG mutants and endogenous proteins, UBR4 KO TREx 293 cells in 6-well plates were transfected with 1 µg total DNA. The amount of each UBR4 mutant plasmid was varied to yield comparable expression levels, and was supplemented with empty vector where required. After 48 h, cells were harvested and lysed on ice in native IP buffer (50 mM HEPES-KOH, 100 mM KAc, 2 mM Mg(Ac)_2_, and 0.01% digitonin) supplemented with 1x protease inhibitor cocktail (Sigma-Aldrich). Lysates were incubated on ice for 15-30 min and spun at 15,000 rpm for 10 min at 4°C on a tabletop centrifuge to remove the insoluble fraction. The cytosolic fractions were then incubated with 10 µL of anti-FLAG M2 affinity gel at 4°C for 1-1.5h with end-over-end rotation. The resin was then washed 5 times with native IP buffer and tubes were exchanged with the 4^th^ wash. Bound proteins were eluted either with 0.25 mg/mL 3xFLAG peptide in PSB for 20 min at room temperature with constant gentle mixing or by boiling with SDS sample buffer.

### Immunoblot analysis

Whenever possible, protein concentrations were normalized using absorbance at 280 nm. Samples were mixed with SDS sample buffer, separated by Tris-Tricine SDS-PAGE and electro-transferred to a 0.2 µm nitrocellulose membrane. Membranes were then blocked with 5% milk in PBS-T (PBS containing 0.1% Tween 20) at room temperature for 30-60 min and incubated with the appropriate primary antibodies at 4°C overnight. Membranes were then washed with PBS-T and incubated with HRP-conjugated secondary antibodies for 1h at room temperature, followed by extensive PBS-T washing. Blots were subsequently exposed to Pierce ECL substrate (Thermo Fisher Scientific), SuperSignal West Pico Chemiluminescent Substrate (Thermo Fisher Scientific), or Immobilon Western Chemiluminescent HRP Substrate (Millipore) and chemiluminescent signal was detected by X-ray film or imaged on a ChemiDoc MP Imaging System (Bio-Rad).

### PURE in vitro translation

Translations in the PURE system (Protein synthesis Using Recombinant Elements) were performed using the PURExpress Δ(aa,tRNA) Kit (New England Biolabs). Translation reactions were assembled with 10 ng/µL of the appropriate plasmid DNA, 100 µM mix of 19 amino acids (minus Met), 0.8 U/µL recombinant RNAsin (Promega), 25 µM methionine and 1 µCi/µL ^35^S-methionine. After translation at 37°C for 1h, 20 µL of sample was layered onto a 200 µL 5-25% sucrose gradient. The gradients were prepared in 7 x 20 mm centrifuge tubes (Beckman Coulter, Cat#343775) by successively layering 40 µL each of 25%, 20%, 10% and 5% sucrose (w/v) in PSB. Gradients were allowed to settle for 30-60 min at 4°C. Samples were centrifuged in a TLS-55 rotor (Beckman Coulter) at 55000 rpm at 4°C with slow acceleration and deceleration for 2h and 25 min. After centrifugation, 20 µL fractions were taken and analyzed by SDS-PAGE and autoradiography. Fractions containing soluble protein were combined and used for *in vitro* ubiquitination reactions.

### In vitro ubiquitination reactions

For single-step ubiquitination reactions by the UBR4-KCMF1 complex, radiolabelled substrates produced in the PURE system were mixed with 1 mM ATP, 10 mM creatine phosphate, 40 µg/mL creatine kinase, 10 µM His-ubiquitin (WT or mutants, as indicated), 100 nM GST-UBE1 and 1 µM UBE2A in PSB, with or without ∼150 nM of UBR4-strep and ∼150 nM of KCMF1-strep, as specified. Reactions were incubated at 32°C for 30 min and stopped by boiling at 95°C in 100 mM Tris-HCl pH 8.0 with 1% SDS. Ubiquitinated products were then recovered by His-Ubiquitin pulldown as described above.

For the sequential CCT4 ubiquitination reactions, untagged CCT4 produced in the PURE system was first mixed with the components described above (ATP, creatine phosphate, creatine kinase, His-ubiquitin, GST-UBE1) together with 250 nM UBE2D1 instead of UBE2A, as well as ∼15 nM 3xFLAG HERC2 and ∼800 nM ZNRD2-3xFLAG. After the ubiquitination reaction, samples were diluted 10-fold in native pulldown buffer (50 mM HEPES pH 7.4, 200 mM NaCl, 2 mM Mg(Ac)_2_, 0.5% Triton, 20 mM Imidazole) and incubated with 10 µL Ni-NTA agarose (Qiagen) at 4°C for 1.5 h. Resin was washed four times with native pulldown buffer and transferred to fresh tubes with the 4^th^ wash. A fifth wash was done with PSB without detergent and ubiquitinated samples were eluted in 250 mM Imidazole in PSB for 30 min at 4°C with gentle mixing. The eluted products were used in a second ubiquitination reaction containing FLAG-Ubiquitin, UBE2A, UBR4 and KCMF1 in the concentrations specified above. Reactions were denatured in 100 mM Tris-HCl pH 8.0, 1% SDS and recovered by a denaturing FLAG-IP using 10 µL of anti-FLAG M2 affinity gel incubated at 4°C overnight with end-over-end rotation. Resin was washed 3 times and samples eluted with SDS sample buffer.

### Sucrose gradient fractionation of cytosolic extracts

To analyze the migration of KCMF1 and UBR4 on a sucrose gradient, cell lysates from Flp-in TREx WT and KCMF1 and UBR4 KO cells were prepared by lysing with 0.01% digitonin in PSB as described earlier. 100 µg of cell lysate in a volume of 20 µL was layered onto a 200 µL 5-25% sucrose gradient as described above. Samples were centrifuged in a TLS-55 rotor (Beckman Coulter) at 55000 rpm at 4°C with slow acceleration and deceleration for 45 min. 20 µL fractions were then taken and analyzed by SDS-PAGE and immunoblot.

### Native PAGE

Cell lysates from GFP-PSMB4-P2A-RFP or PSMB4-GFP-P2A-RFP stable cell lines were lysed in PSB with 0.01% digitonin and protease inhibitors, and insoluble material removed by centrifugation, as described above. Protein concentration was calculated from absorbance at 280 nm, and 10-30 µg protein was for each condition was mixed with Novex^TM^ Tris-Glycine Native sample buffer (2X) and loaded on Novex™ Tris-Glycine Mini Protein Gels, 4–12%, 1.0 mm, WedgeWell™ format (Thermo Fisher Scientific). In-gel GFP and RFP fluorescence was then imaged on a ChemiDoc MP Imaging System (Bio-Rad).

### Structural modelling

Structural modelling employing AlphaFold3^42–44^ was used to generate a composite model of the UBR4-KCMF1-UBE2A-CALM complex as follows. In preliminary pair-wise predictions (using AlphaFold-multimer), we determined the sites of high-confidence interaction on UBR4 for the NTD of KCMF1, the CTD of KCMF1, UBE2A, and CALM. High-confidence was taken as PAE values below ∼10 Å at the interface with essentially identical architectures seen in the top five models. With this information, we then performed three separate overlapping predictions using AF3 (see Fig. S6A-S6C): UBR4(1-1831); UBR4(1549-4301) + KCMF1 + CALM; UBR4(4193-5183) + UBE2A.

The regions of overlap for UBR4 were used to align the three predictions, using a bundle of two or three alpha-helices as a guide. This showed that the N-arm and C-arm are relatively heterogeneous in their overall positions, with no evidence for an interaction between them (verified by a pairwise AF3 prediction using the two arms). Sub-domains whose positions were essentially invariant between the five models (such as the beta-propeller domain) were noted and verified to have low PAE scores with scaffold regions of UBR4. Similarly, sub-modules of multiple domains whose relative configurations were high-confidence and essentially invariant across the top five models (e.g., Fig. S6B) were also verified to have low PAE scores between them. By contrast, the position of some subdomains were variable across models (e.g., the UBL domain; see Fig. S6C), and did not have low PAE scores with any other region of UBR4. These observations were used to guide interpretations.

To identify putative ubiquitin-binding site(s) on UBR4, we performed a pairwise prediction with UBR4(251-5183) and ubiquitin (Fig. S6D), which showed a single high-confidence site as judged by low PAE scores and very close alignment of all five models for this module. The site of ubiquitin on a putative charged UBE2A was also predicted by AF3, resulting in five near-identical models in which ubiquitin is positioned at its expected site with G76 abutting the site of charging on UBE2A (C88). These two predictions were used to generate a hypothetical ubiquitin-transfer complex (Fig. 5D) to determine which (if any) lysine on the UBL-bound ubiquitin might be in a position to react with C88 of charged UBE2A. Figure preparation was done using PyMOL.

### Quantification and statistical analysis

Quantification of fluorescent intensities from Native-PAGE gels was done using Fiji. The total intensity values were rectified by subtracting background values. GFP:RFP ratios were calculated with the adjusted values and the ratios were normalized to the control condition. The bar graphs show the mean and the individual ratios from three independent experiments. Statistical analysis was performed in GraphPad Prism using an upaired, two-tailed Student’s *t*-test to compare two individual conditions. The *p*-values (p) are shown in the figures with the following symbols: * p <0.05, ** p <0.01, ** p <0.001, **** <0.0001. For the mass spectrometry data, statistical analysis to identify interactors of each bait protein was performed in Perseus using a two-tailed Student’s *t*-test with Benjamini-Hochberg correction for multiple comparisons.

## Acknowledgements

We thank J. Mark Skehel for performing mass spectrometry as part of the LMB facility, Paul Elliott for useful discussions, and J. Neuhold from the Vienna BioCenter Core Facility in Protein Technology for her support. This work was supported by the Medical Research Council, as part of United Kingdom Research and Innovation (MC_UP_A022_1007 to R.S.H.), LMB-AZ BlueSky project BSF27 (E.Z. and R.S.H.), an Esprit Grant from the Austrian Science Foundation ESP 218-B (P.M.) and Austrian Research Promotion Agency Headquarter grant 852936 (T.C.). The IMP is supported by Boehringer Ingelheim.

## Author contributions

SCR, YY, EZ, and PM performed experiments. PM and RK produced recombinant UBR4 and KCMF1. EZ and RSH conceived the project, parts of which were independently conceived by PM and TC. SCR, EZ and RSH wrote the manuscript, with edits provided by TC and YY. TC, EZ and RSH provided project supervision.

---

**Figure SI.**
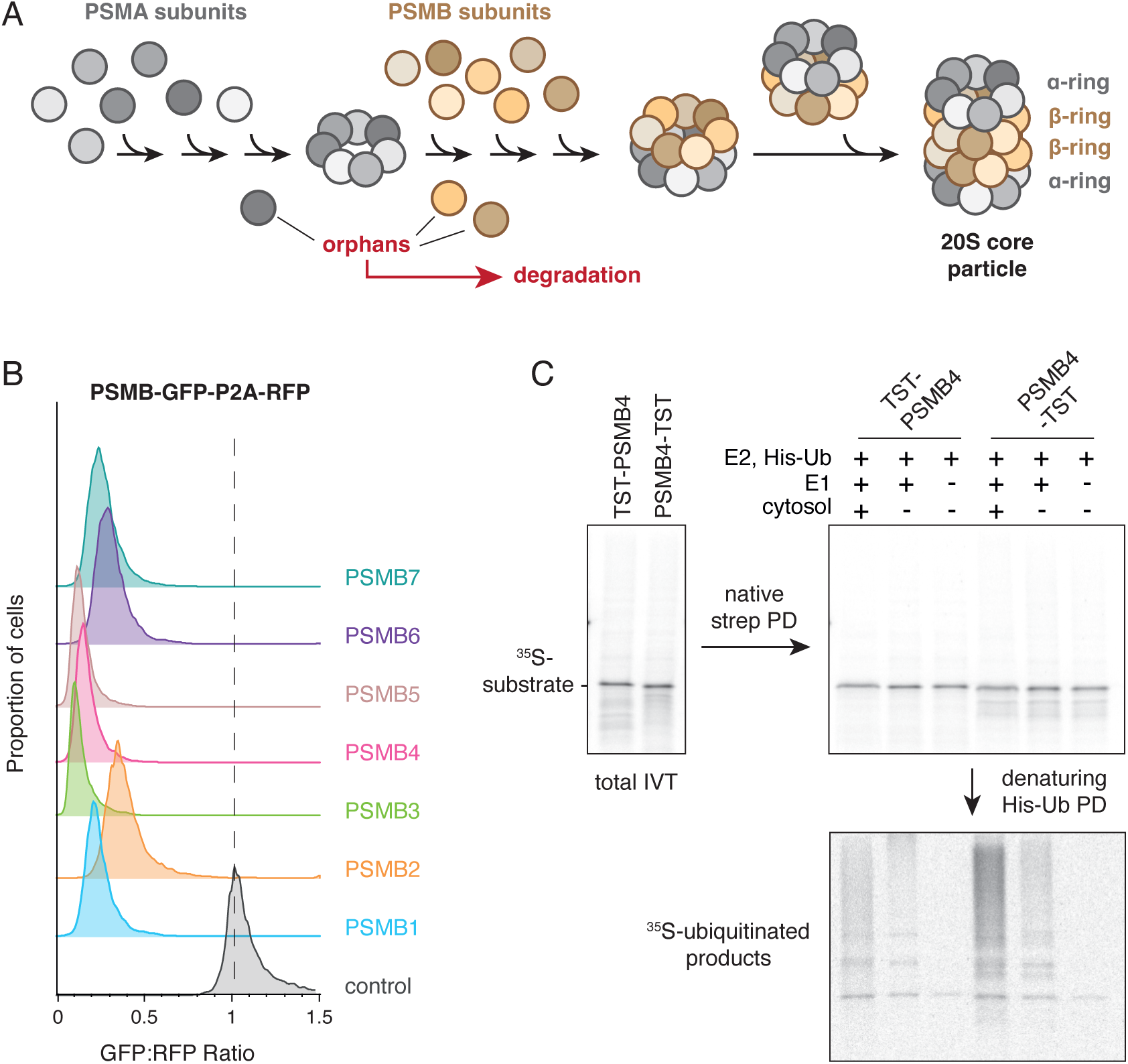
In cell and in vitro analysis of orphaned PSMB subunits, related to Figure 1. (A) Simplified schematic depicting 20S core proteasome assembly and quality control. The hetero-heptameric alpha ring is formed from subunits PSMA1-PSMA7, and the hetero-heptameric beta ring is formed from subunits PSMB1-PSMB7. The 20S core complex is comprised of two rings of each type. Any unassembled subunits (e.g., from imbalanced synthesis), defined as orphans, are degraded. Chaperones, assembly factors and processing steps are not depicted for simplicity. **(B)** Stable-inducible cell lines containing each of the seven PSMBx-GFP reporters and a control GFP reporter (lacking the PSMB insert) were induced with dox for 48Һ and analyzed by flow cytometry. The stability of each reporter was assessed relative to a co-expressed RFP control (see Fig. 1A), with the GFP:RFP ratio being plotted as a histogram. Note that all of the PSMBx reporters are degraded relative to the control. (C) PSMB4 tagged with a N-terminal or C-terminal TwinStrep tag (TST) was translated in rabbit reticulocyte lysate (RRL) with ^35^S-methione and affinity-purified via TST under native conditions. The affinity-purified products were divided into three aliquots and incubated with El, E2 (UBE2D1), His-Ub, ATP and cytosol (RRL), as indicated, then subjected to a pulldown via His-Ub under denaturing conditions. Aliquots of the samples at each step of the process were analyzed by SDS-PAGE and autoradiography: total IVT, the total products of the ubiquitination reaction, and the products recovered by His-Ub pulldown.

**Figure S2.**
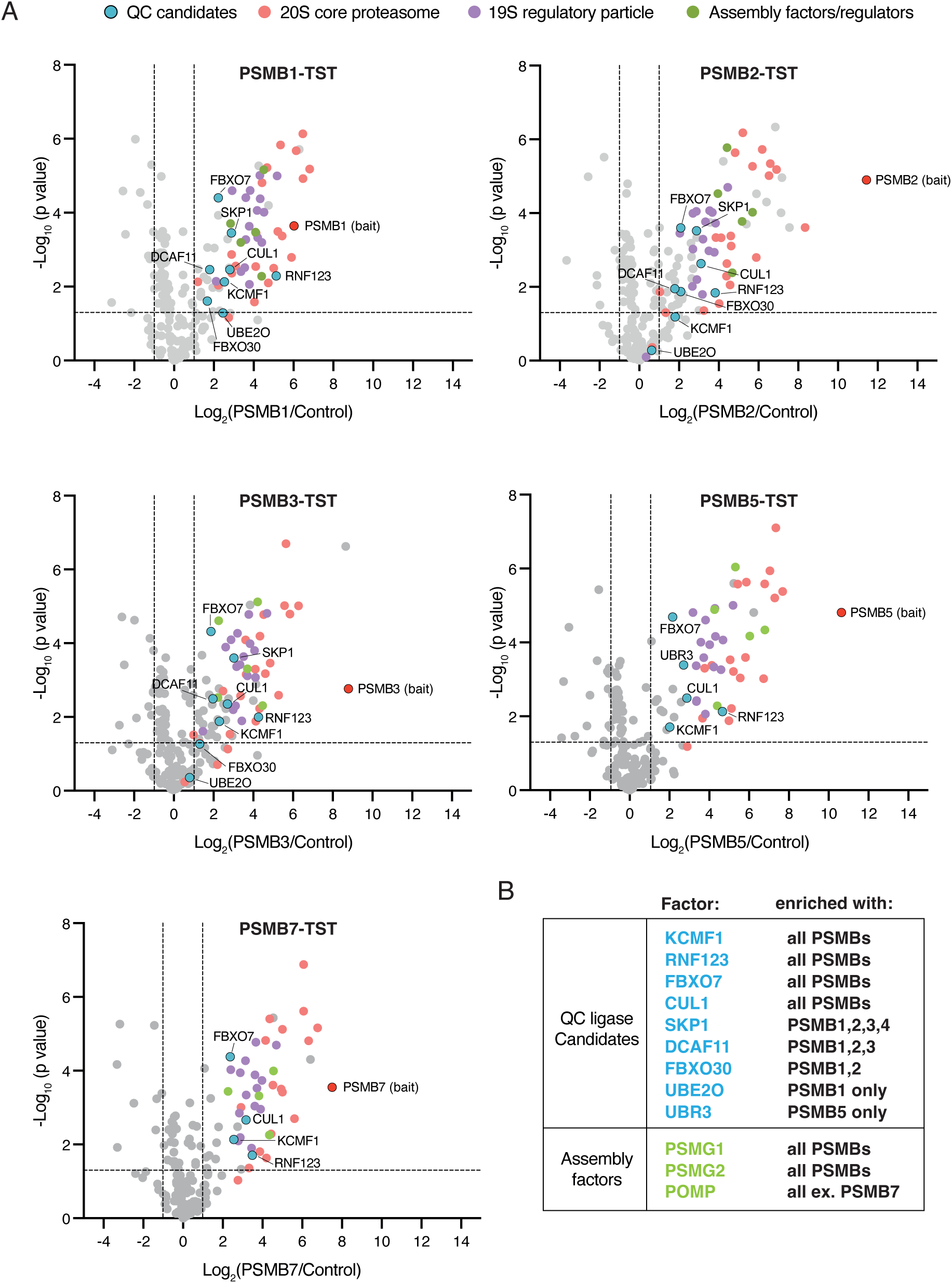
**Identification of quality control candidates of orphaned PSMB subunits, related to Figure 2. (A) C**-terminal TST-tagged PSMB subunits were translated in RRL, affinity-purified under native conditions and analyzed by label free quantitative mass spectrometry. The volcano plots show proteins enriched in each of the PSMB subunit pulldowns compared to a mock translation which served as a negative control. P-values were calculated by two-sided Student’s Z-test with Benjamini Hochberg correction for multiple comparisons. The translation reaction for PSMB6 failed in this experiment, so results for this sample are not shown. **(B)** Table showing QC ligase candidates and assembly factors that were enriched in at least one of the PSMB pulldowns. Recovery of the assembly factors indicates that a proportion of each translation product begins but does not complete assembly, indicative of a population of orphaned subunits in the reaction.

**Figure S3.**
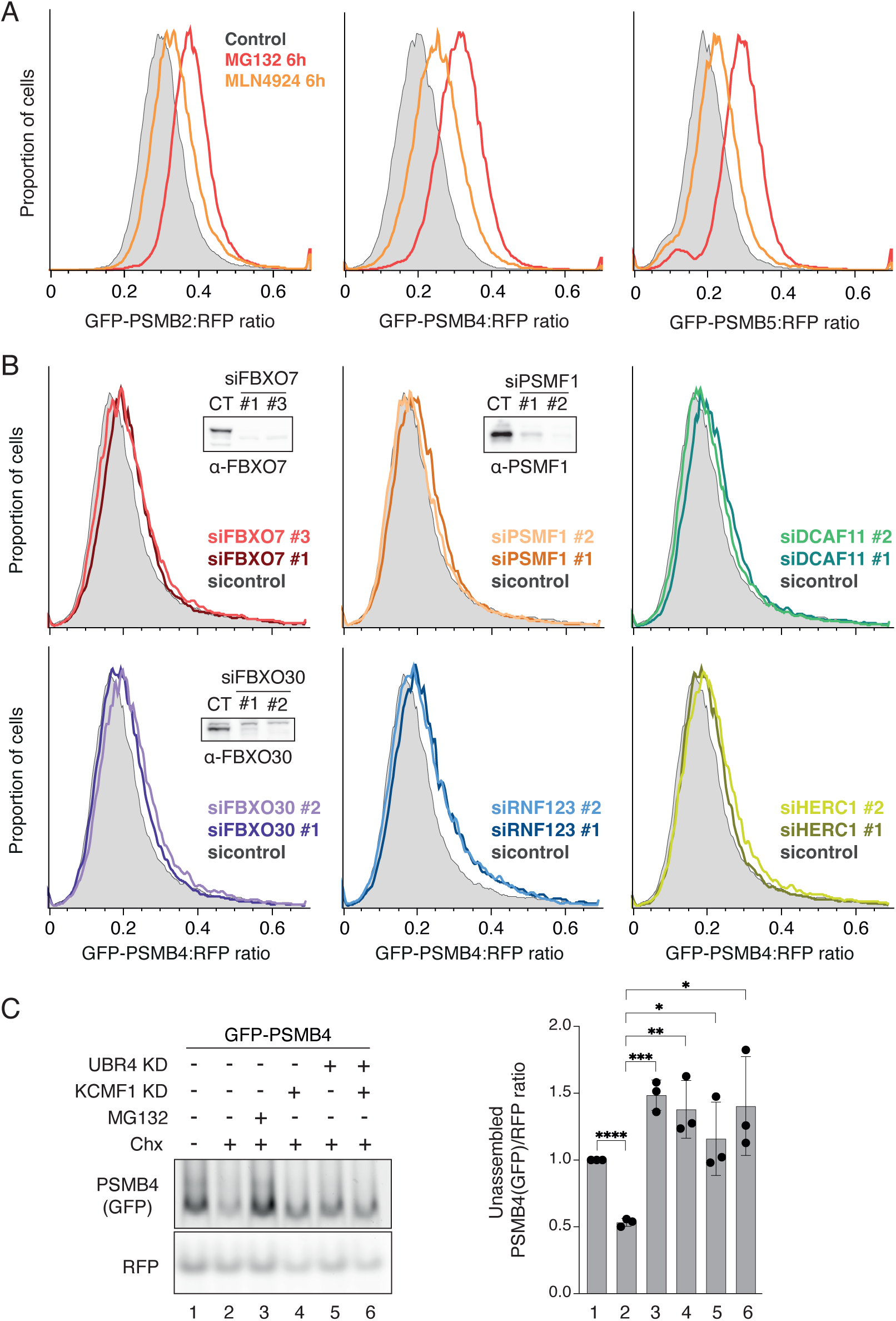
Analysis of quality control candidates in the degradation of orphaned PSMB subunits, related to Figure 2. **(A)** Stable-inducible cell lines containing the indicated GFP-PSMB reporter were induced with dox for 18Һ. After removal of dox, cells were treated with 20 рМ MG132 or 1 pM MLN4924 (a neddylation inhibitor) for 6 hours, then analyzed by flow cytometry. **(B)** The stable-inducible cell line containing the GFP-PSMB4 reporter was transfected with non-targeting control or the specified siRNAs for a total of 72Һ. GFP-PSMB4 reporter expression was induced with dox for the last 18h. Cells were then analyzed by flow cytometry and immunoblot (where suitable antibodies were available). (C) The GFP-PSMB4 reporter cell line was transfected with non-targeting control, KCMF1 or UBR4 siRNAs for a total of 72Һ. GFP-PSMB4 reporter expression was induced with dox after 54Һ for a period of 18h. After removal of dox, cells were treated with 100 pg/mL CHXand20 pMMG132 for 6Һ, as indicated. Cell lysates were then analyzed by Native PAGE and in-gel fluorescence to detect GFP and RFP (left). The ratio of unassembled PSMB4-GFP fluorescence to RFP, relative to untreated control, was quantified from three replicates from two independent experiments and plotted (right). Single, double, triple and quadruple asterisk indicate Student’s t-test p values less than 0.05, 0.01, 0.001 and 0.0001, respectively. Error bars indicate standard deviation.

**Figure S4.**
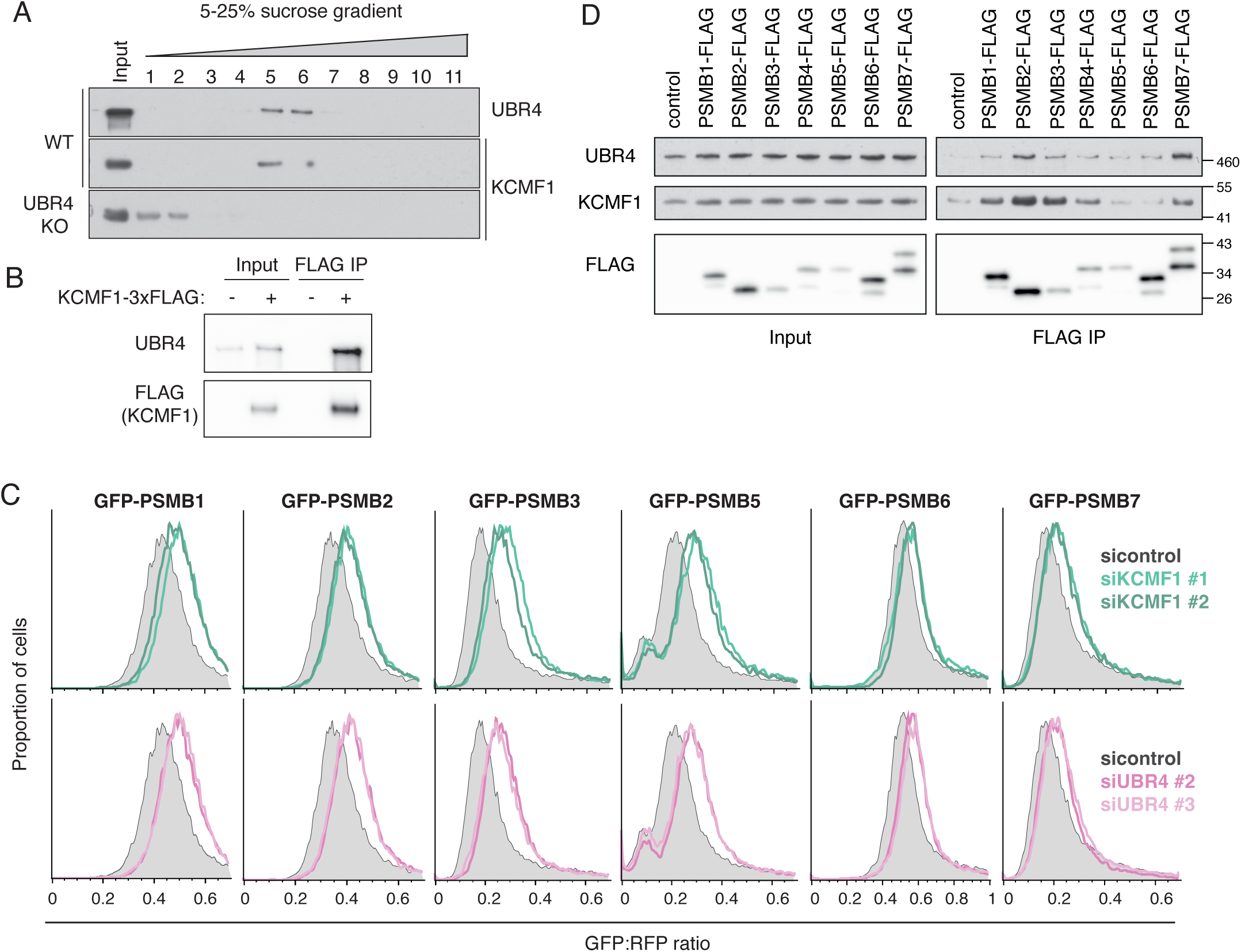
The UBR4-KCMF1 complex is required for the degradation of multiple orphaned PSMB subunits, related to Figure 2. (A) Total cell lysates (Input) from wildtype (WT) or AUBR4 cells were separated on a 5-25% sucrose gradient and analyzed by immunoblot. Note that UBR4 and KCMF1 co-migrated as a large complex in fractions 5-6 in WT cells, but that KCMF1 shifted to fractions 1-2 when UBR4 was knocked out. **(B)** AKCMF1 cells stably integrated with dox-inducible KCMFl-3xFLAG were induced with dox for 18h and subjected to anti-FLAG IP under native conditions. Input and FLAG IP samples were analyzed by immunoblot. **(C)** Cells stably integrated with inducible GFP-PSMBx reporters were transfected with non-targeting control, KCMF1 or UBR4 siRNAs for a total of 72Һ. Expression of GFP-PSMBs was induced with dox for the last 18h and cells were analyzed by flow cytometry. **(D)** HEK293T cells transiently transfected with C-terminally FLAG-tagged PSMB subunits were subjected to anti-FLAG IP under native conditions. Input and FLAG IP samples were analyzed by immunoblot.

**Figure S5.**
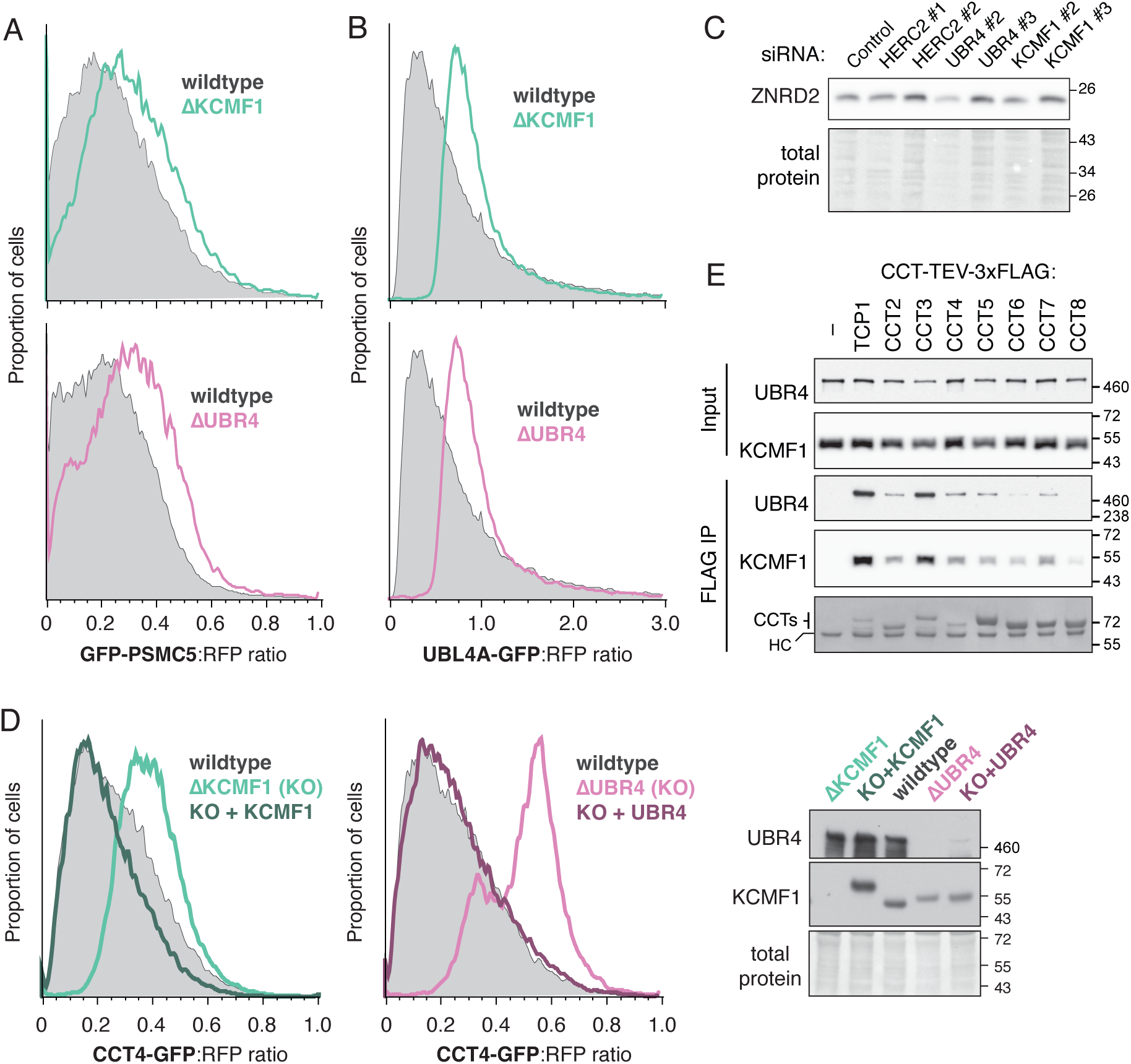
The UBR4-KCMF1 complex is required for the degradation of orphaned PSMC5, UBL4A and CCT4, related to Figure 3. **(A)** Wildtype, AKCMF1 or AUBR4 cells were transiently transfected with GFP-PSMC5 reporter for 48Һ and analyzed by flow cytometry. Note that the steady-state level of orphaned GFP-PSMC5 is higher (relative to the internal RFP control) in both KO cells. **(B)** Same as A but with the UBL4A-GFP reporter. **(C)** Lysates from cells treated with the indicated siRNAs (see Fig. 3A) were analyzed by immunoblot for ZNRD2, whose levels do not change in any of the conditions. Note that lane 4 is slightly underloaded in this blot as indicated by the total protein image. **(D)** Wildtype, AKCMF1 or AUBR4 cells were transiently co-transfected with the CCT4-GFP reporter and KCMFl-3xFLAG or UBR4-3xFLAG constructs, as indicated, for 48Һ. Cells were then analyzed by flow cytometry (left and middle) or immunoblot (right). By comparison to KCMF1, re-expression of UBR4 is relatively low due to reduced overall transfection efficiency (in terms of both copy number and percent of cells) due to the very large plasmid size. Nonetheless, analysis of the transfected cells by flow cytometry indicates complete rescue of the degradation phenotype. **(E)** HEK293T cells transiently transfected with C-terminally FLAG-tagged CCT subunits were subjected to anti-FLAG IP under native conditions. Input and FLAG IP samples were analyzed by immunoblot for UBR4 and KCMF1, or total protein stain for CCTs.

**Figure S6.**
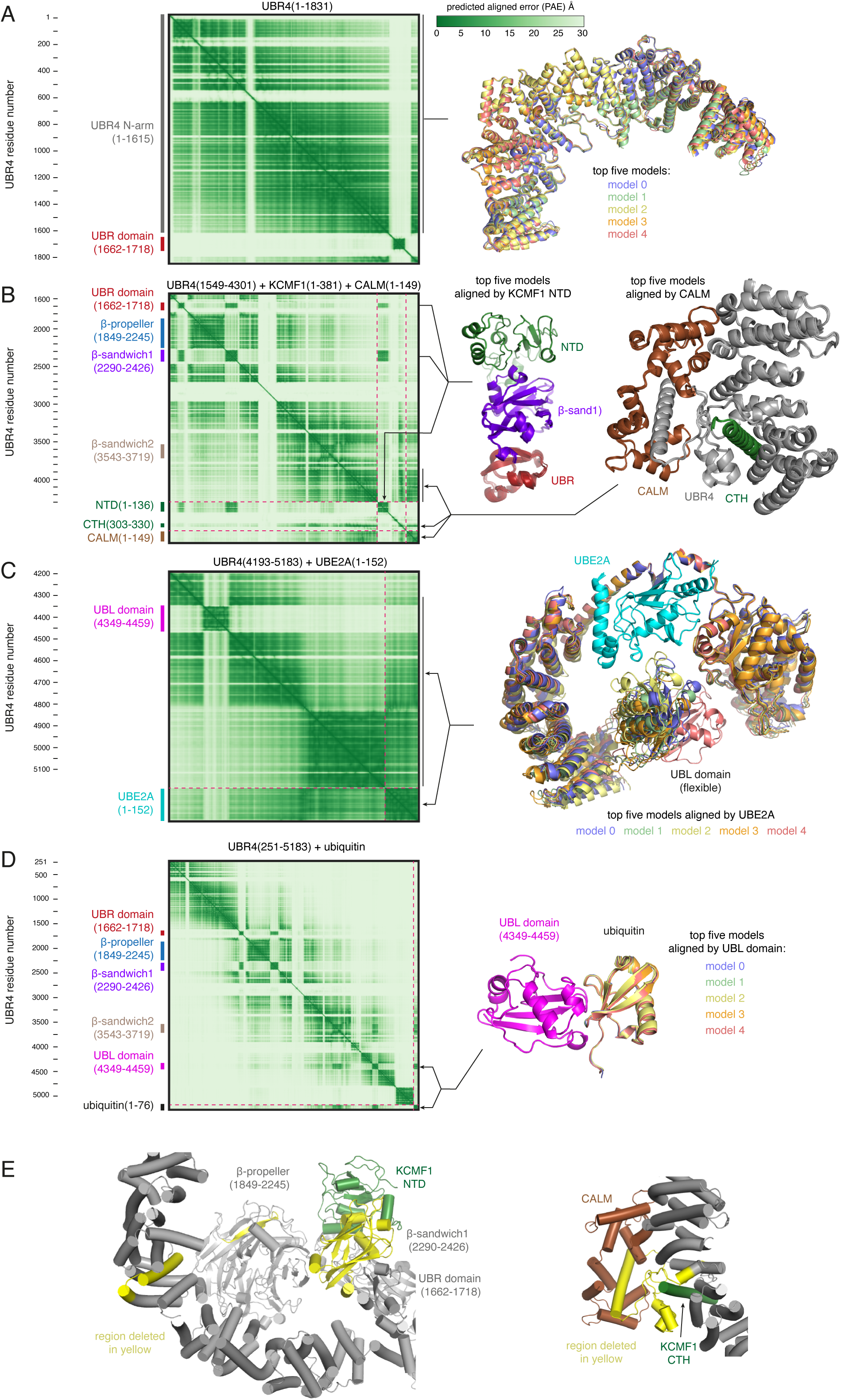
Structural modelling of the UBR4-KCMF1-UBE2A-CALM complex and its interactions with ubiquitin, related to Figure 5. **(A)** AlphaFold3 prediction of UBR4(1-1831). Shown at left is the matrix of predicted aligned error for the top predicted model, annotated on the left side with amino acid numbers and key domains. Shown on the right is an alignment of the N-arm for the top five models. The UBR domain, whose fold is predicted with high confidence, is positioned differently in all the models due to its lack of interaction with the N-arm (evident on the PAE plot). Instead, the UBR domain is tethered by two flexible linkers. **(B)** AlphaFold3 prediction of UBR4(1549-4301) with KCMF1 and CALM. The PAE plot is displayed as in A, with the different proteins and domains highlighted on the left. On the right, alignments of the top five models for two key high-confidence interactions are displayed, with arrows indicating the regions of low PAE between the interacting domains. The KCMF1(NTD)-13-sandwichl-UBR models were aligned on the NTD; the CALM-UBR4-CTH models were aligned on CALM. **(C)** AlphaFold3 prediction of UBR4(4193-5183) with UBE2A displayed as in A. Alignment of the top five models was on UBE2A. Note the apparent flexibility of the UBL domain’s position due to its attached flexible linkers. **(D)** AlphaFold3 prediction of UBR4 (251-5183) and ubiquitin. Note the high-confidence interaction, indicated on the PAE plot, between the UBL domain of UBR4 and ubiquitin. Shown on the right is an alignment of the top five models (aligned on the UBL domain) of this UBL-ubiquitin module. (E) Close-up views of the AlphaFold3-predicted UBR4-KCMF1-CALM composite model highlighting the regions in UBR4 (yellow) that had been previously deleted^35^ to created UBR4 mutants deficient in KCMF1 binding (left) or CALM binding (right). Note that although these deletions would disrupt binding to the interacting domain as intended, both of them also encroach on key structural elements of the core UBR4 scaffold.

**Figure S7.**
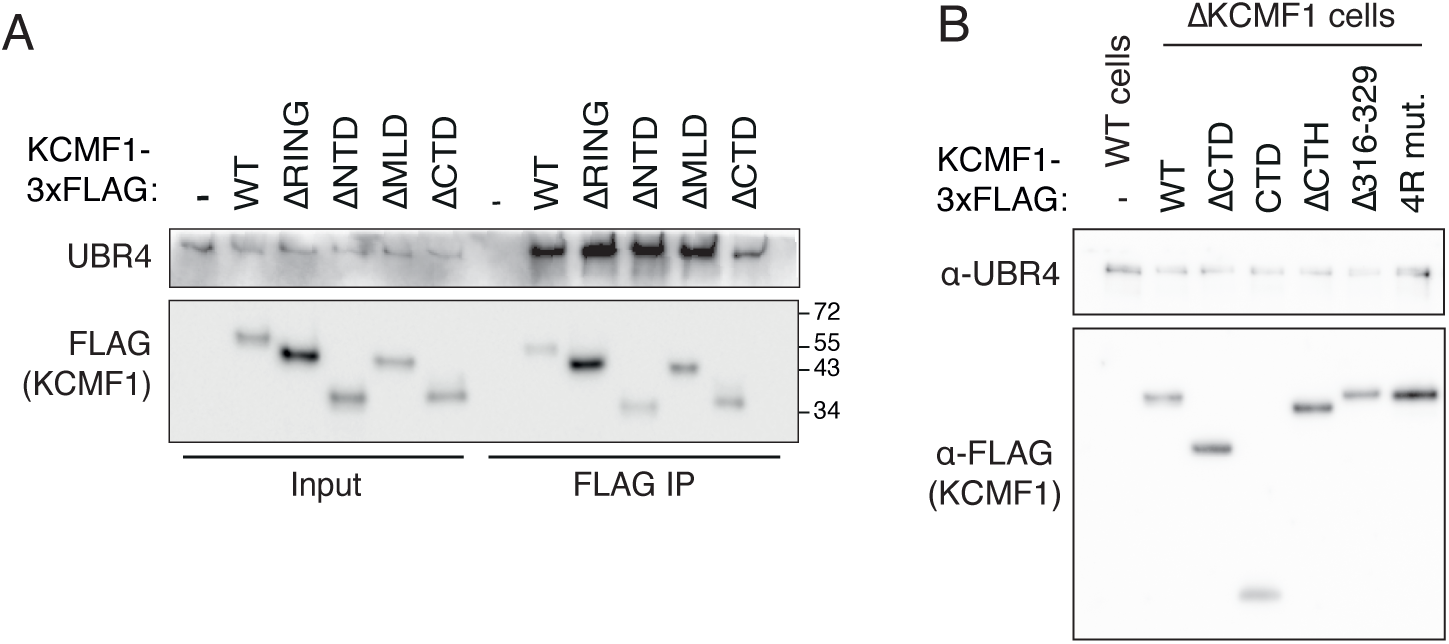
Analysis of KCMF1 deletion mutants, related to Figure 6. **(A)** AKCMFl cells were transiently transfected with the indicated KCMFl-3xFLAG constructs for 48Һ. Cells were then subjected to anti-FLAG IP under native conditions and input and IP samples were analyzed by immunoblot. **(B)** AKCMFl cells were transiently transfected with the indicated KCMFl-3xFLAG constructs for 48Һ. Expression levels of KCMF1 and UBR4 were analyzed by immunoblot.

**Figure S8.**
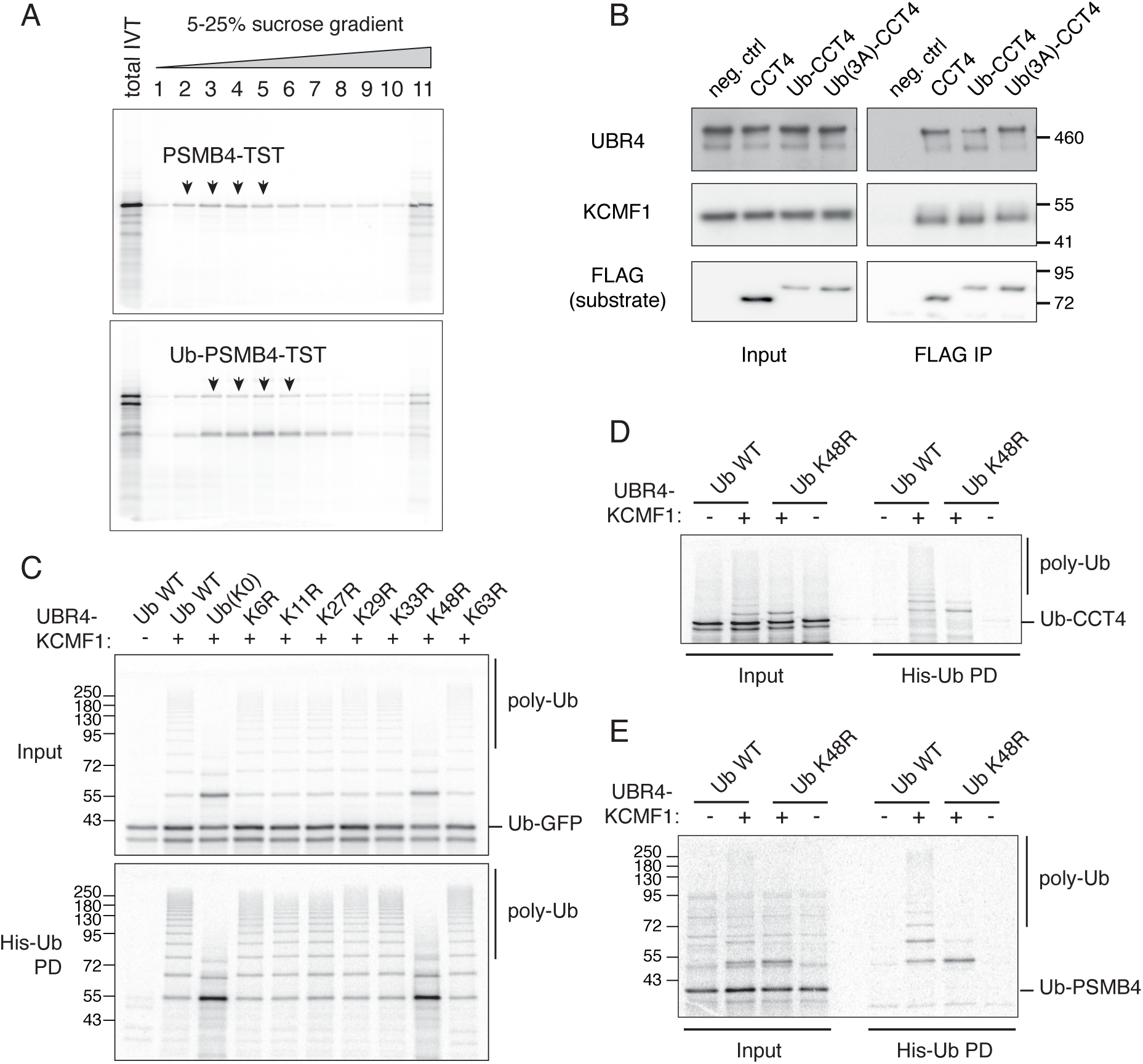
Functional and interaction analysis of the UBR4-KCMF1 complex, related to Figure 7. **(A)** Preparation of ^35^S-methionine-labelled PSMB4 and ubiquitin(G76V)-fused PSMB4 using the PURE translation system. After in vitro translation, the reactions were separated on a 5-25% sucrose gradient, and resulting fractions were analyzed by SDS-PAGE and autoradiography. The arrowheads indicate the soluble fractions that were pooled and used as substrates for subsequent *in vitro* ubiquitination assays. Other ubiquitination substrates were prepared similarly. Further analysis revealed that the prominent smaller molecular weight band seen in the Ub-fused translation reactions is a tRNA-linked peptide generated by out-of-frame translation initiation from a cryptic Shine-Dalgamo sequence in the ubiquitin open reading frame. This peptidyl-tRNA does not interfere in the ubiquitination reactions. **(B)** FLAG-tagged CCT4, Ub-CCT4 and Ub(3A)-CCT4 were translated in RRL in the presence of 20 pM El inhibitor (TAK-243) and subjected to FLAG IP under native conditions. Input and IP samples were then analyzed by immunoblot. (C) ^35^S-methionine-labelled Ub-GFP translated in the PURE system was incubated with El, E2 (UBE2A), His-Ub (WT, KO or the indicated K-to-R mutants), ATP and recombinant UBR4 and KCMF1 as indicated. The samples were then analyzed by autoradiography either directly (Input) or after a His-Ub PD under denaturing conditions. **(D)** ^35^S -methionine-labelled 1Љ-ССТ4 translated in the PURE system was incubated with El, E2 (UBE2A), His-Ub (WT or K48R), ATP and recombinant UBR4 and KCMF1 as indicated. The samples were then analyzed by autoradiography either directly (Input) or after a His-Ub PD under denaturing conditions. (E) As in panel D except using Ub-PSMB4 as the substrate.

## Notes

### Competing Interest Statement

The authors have declared no competing interest.

