## Supplemental Table 1 for "Convergence of orphan quality control pathways at a ubiquitin chain-elongating ligase"

**Table S1 - Proteins recovered with PSMB subunits identified by mass spectrometry. Related to Figure 2A and Figure S2.**

| Uniprot ID | Protein name | Gene name | Protein of interest | Log2(PSMB1/Control) | -Log10(adj. p value)_PSMB1vsCont | Enriched in PSMB1 pulldown | Log2(PSMB2/Control) | -Log10(adj. p value)_PSMB2vsCont | Enriched in PSMB2 pulldown | Log2(PSMB3/Control) | -Log10(adj. p value)_PSMB3vsCont | Enriched in PSMB3 pulldown | Log2(PSMB4/Control) | -Log10(adj. p value)_PSMB4vsCont | Enriched in PSMB4 pulldown | Log2(PSMB5/Control) | -Log10(adj. p value)_PSMB5vsCont | Enriched in PSMB5 pulldown | Log2(PSMB7/Control) | -Log10(adj. p value)_PSMB7vsCont | Enriched in PSMB7 pulldown |
|---|---|---|---|---|---|---|---|---|---|---|---|---|---|---|---|---|---|---|---|---|---|
| G1U3G0 | Ubiquitin conjugating enzy | UBE2O | E3 | 2.45 | 1.29 | enriched | 0.62 | 0.28 | | 0.79 | 0.36 | | -0.87 | 0.35 | | -0.42 | 0.15 | | -0.91 | 0.38 | |
| G1SZL9 | F-box protein 30 | FBXO30 | E3 | 1.67 | 1.61 | enriched | 2.09 | 1.87 | enriched | 1.30 | 1.27 | | 0.00 | 0.00 | | 0.00 | 0.00 | | 0.00 | 0.00 | |
| tG1SIF7 | DDB1- and CUL4-associate | DCAF11 | E3 | 1.79 | 2.46 | enriched | 1.79 | 1.95 | enriched | 1.97 | 2.50 | enriched | 0.00 | 0.00 | | -0.05 | 0.01 | | 0.00 | 0.00 | |
| G1TTU6 | S-phase kinase-associated | SKP1 | E3 | 2.88 | 3.45 | enriched | 2.88 | 3.52 | enriched | 3.03 | 3.60 | enriched | 3.07 | 3.94 | enriched | 1.19 | 0.51 | | 1.03 | 0.43 | |
| G1SHR7 | Potassium channel modula | KCMF1 | E3 | 2.53 | 2.13 | enriched | 1.81 | 1.19 | enriched | 2.30 | 1.88 | enriched | 2.15 | 1.86 | enriched | 2.01 | 1.71 | enriched | 2.55 | 2.13 | enriched |
| G1TTN8 | RING-type domain-contain | RNF123 | E3 | 5.13 | 2.29 | enriched | 3.82 | 1.84 | enriched | 4.26 | 2.00 | enriched | 4.92 | 2.22 | enriched | 4.67 | 2.13 | enriched | 3.49 | 1.71 | enriched |
| G1SUV8 | F-box protein 7 | FBXO7 | E3 | 2.22 | 4.40 | enriched | 2.09 | 3.60 | enriched | 1.87 | 4.32 | enriched | 3.26 | 4.87 | enriched | 2.16 | 4.69 | enriched | 2.38 | 4.38 | enriched |
| G1SHS4 | Cullin 1 | CUL1 | E3 | 2.80 | 2.46 | enriched | 3.11 | 2.63 | enriched | 2.70 | 2.35 | enriched | 3.85 | 2.99 | enriched | 2.87 | 2.49 | enriched | 3.17 | 2.66 | enriched |
| G1SRE9 | E3 ubiquitin-protein ligase | UBR3 | E3 | 0.00 | 0.00 | | 0.00 | 0.00 | | 0.00 | 0.00 | | 0.00 | 0.00 | | 2.71 | 3.39 | enriched | 0.00 | 0.00 | |
| tr\|G1T2L1\|G1T2L1_RABIT | Proteasome subunit alpha | PSMA2 | 20S | 4.67 | 5.22 | enriched | 4.82 | 5.64 | enriched | 4.34 | 4.19 | enriched | 5.04 | 5.64 | enriched | 5.43 | 5.58 | enriched | 4.37 | 5.40 | enriched |
| tr\|G1SQU1\|G1SQU1_RABIT | Proteasome subunit beta ( | PSMB10 | 20S | 2.90 | 2.36 | enriched | 4.63 | 3.11 | enriched | 3.36 | 2.57 | enriched | 2.52 | 1.64 | enriched | 5.21 | 3.21 | enriched | 0.00 | 0.00 | enriched |
| sp\|P28070\|PSB4_HUMAN | Proteasome subunit beta t | PSMB4 | 20S | 4.73 | 2.10 | enriched | 4.57 | 2.05 | enriched | 4.13 | 1.89 | enriched | 11.18 | 3.50 | enriched | 5.11 | 2.21 | enriched | 3.87 | 1.80 | enriched |
| sp\|P28074\|PSB5_HUMAN | Proteasome subunit beta t | PSMB5 | 20S | 5.42 | 3.37 | enriched | 4.60 | 3.38 | enriched | 4.85 | 3.46 | enriched | 5.40 | 3.65 | enriched | 10.65 | 4.81 | enriched | 4.94 | 3.50 | enriched |
| sp\|P49720\|PSB3_HUMAN;tr\|( | Proteasome subunit beta t | PSMB3 | 20S | 4.04 | 1.58 | enriched | 4.02 | 1.54 | enriched | 8.79 | 2.77 | enriched | 4.07 | 1.60 | enriched | 4.98 | 1.88 | enriched | 4.20 | 1.63 | enriched |
| sp\|P49721\|PSB2_HUMAN | Proteasome subunit beta t | PSMB2 | 20S | 5.21 | 3.49 | enriched | 11.43 | 4.90 | enriched | 4.60 | 3.17 | enriched | 4.57 | 3.29 | enriched | 5.82 | 3.59 | enriched | 5.01 | 3.42 | enriched |
| sp\|Q99436\|PSB7_HUMAN | Proteasome subunit beta t | PSMB7 | 20S | 4.10 | 2.54 | enriched | 4.41 | 2.63 | enriched | 2.82 | 1.54 | enriched | 3.60 | 2.18 | enriched | 5.56 | 3.04 | enriched | 7.51 | 3.55 | enriched |
| tr\|G1SDA8\|G1SDA8_RABIT | Proteasome subunit alpha | PSMA1 | 20S | 5.34 | 5.83 | enriched | 5.21 | 6.18 | enriched | 4.53 | 4.78 | enriched | 5.50 | 6.08 | enriched | 5.86 | 5.63 | enriched | 5.00 | 5.12 | enriched |
| tr\|G1SGW4\|G1SGW4_RABIT | Proteasome subunit beta ( | PSMB9 | 20S | 4.42 | 4.81 | enriched | 5.70 | 5.27 | enriched | 3.61 | 4.09 | enriched | 4.08 | 4.28 | enriched | 6.78 | 5.58 | enriched | 4.16 | 4.82 | enriched |
| tr\|G1SIH1\|G1SIH1_RABIT | Proteasome activator subu | PSME2 | 20S | 4.12 | 3.45 | enriched | 3.86 | 3.34 | enriched | 3.64 | 3.16 | enriched | 4.64 | 3.65 | enriched | 3.77 | 3.30 | enriched | 4.52 | 3.61 | enriched |
| tr\|G1SWI7\|G1SWI7_RABIT | Proteasome 20S subunit al | PSMA8 | 20S | 6.48 | 4.92 | enriched | 6.53 | 5.01 | enriched | 5.85 | 4.79 | enriched | 6.65 | 5.05 | enriched | 7.29 | 5.21 | enriched | 6.32 | 4.81 | enriched |
| tr\|G1SZ14\|G1SZ14_RABIT | Proteasome subunit alpha | PSMA3 | 20S | 6.13 | 5.67 | enriched | 6.19 | 5.73 | enriched | 5.57 | 5.02 | enriched | 6.33 | 5.63 | enriched | 7.04 | 5.94 | enriched | 6.07 | 5.61 | enriched |
| tr\|G1T235\|G1T235_RABIT;sp\| | Proteasome endopeptidase | PSMB6 | 20S | 2.75 | 1.16 | enriched | 3.25 | 1.35 | enriched | 2.72 | 1.13 | enriched | 3.51 | 1.45 | enriched | 2.89 | 1.18 | enriched | 3.32 | 1.36 | enriched |
| tr\|G1T4Q9\|G1T4Q9_RABIT | Proteasome subunit beta ( | PSMB5 | 20S | 5.00 | 2.50 | enriched | 4.39 | 2.29 | enriched | 4.32 | 2.23 | enriched | 5.00 | 2.40 | enriched | 3.66 | 1.95 | enriched | 4.44 | 2.28 | enriched |
| tr\|G1T4X8\|G1T4X8_RABIT | Proteasome subunit beta ( | PSMB2 | 20S | 2.89 | 2.87 | enriched | 8.35 | 3.61 | enriched | 2.48 | 2.70 | enriched | 3.38 | 3.25 | enriched | 4.13 | 3.38 | enriched | 2.92 | 3.00 | enriched |
| tr\|G1T519\|G1T519_RABIT | Proteasome subunit alpha | PSMA4 | 20S | 6.81 | 5.18 | enriched | 6.91 | 5.18 | enriched | 6.28 | 5.02 | enriched | 7.13 | 5.25 | enriched | 7.68 | 5.38 | enriched | 6.77 | 5.16 | enriched |
| tr\|G1T670\|G1T670_RABIT;tr\| | Proteasome subunit alpha | PSMA5 | 20S | 5.90 | 2.79 | enriched | 5.89 | 2.80 | enriched | 5.27 | 2.59 | enriched | 6.22 | 2.89 | enriched | 6.73 | 3.02 | enriched | 5.61 | 2.70 | enriched |
| tr\|G1T9V4\|G1T9V4_RABIT | Proteasome subunit alpha | PSMA6 | 20S | 6.47 | 6.13 | enriched | 6.59 | 5.34 | enriched | 5.65 | 6.70 | enriched | 6.61 | 6.37 | enriched | 7.34 | 7.10 | enriched | 6.07 | 6.88 | enriched |
| tr\|G1SU71\|G1SU71_RABIT | Proteasome subunit beta ( | PSMB1 | 20S | 3.10 | 2.56 | enriched | 4.12 | 3.33 | enriched | 4.10 | 3.30 | enriched | 2.15 | 0.46 | | 5.05 | 3.53 | enriched | 2.76 | 1.03 | enriched |
| tr\|G1SGV9\|G1SGV9_RABIT | Proteasome subunit beta ( | PSMB8 | 20S | 2.22 | 2.03 | enriched | 0.65 | 0.36 | | 0.55 | 0.24 | | 2.34 | 2.11 | enriched | 0.00 | 0.00 | | 0.00 | 0.00 | |
| sp\|P20618\|PSB1_HUMAN | Proteasome subunit beta t | PSMB1 | 20S | 6.02 | 3.64 | enriched | 1.33 | 1.30 | | 1.32 | 1.29 | | 2.00 | 1.84 | enriched | 2.24 | 1.98 | enriched | 1.26 | 1.22 | |
| tr\|G1T918\|G1T918_RABIT;tr\| | Proteasome subunit beta ( | PSMB4 | 20S | 0.07 | 0.01 | | 0.00 | 0.00 | | 2.19 | 0.71 | enriched | 3.70 | 2.29 | enriched | 2.19 | 0.51 | | 0.73 | 0.18 | |
| tr\|G1SWK8\|G1SWK8_RABIT | Proteasome subunit beta ( | PSMB7 | 20S | 1.19 | 2.13 | | 1.04 | 1.86 | | 0.99 | 1.52 | | 1.04 | 1.75 | | 1.68 | 2.12 | enriched | 0.00 | 0.00 | |
| tr\|G1SLK2\|G1SLK2_RABIT;tr\|( | AAA domain-containing pro | PSMC5 | 19S | 3.55 | 2.51 | enriched | 2.90 | 2.20 | enriched | 3.10 | 2.31 | enriched | 3.72 | 2.55 | enriched | 3.36 | 2.43 | enriched | 2.88 | 2.19 | enriched |
| tr\|G1TUD6\|G1TUD6_RABIT | 26S proteasome AAA-ATPa | PSMC4 | 19S | 3.61 | 3.27 | enriched | 2.70 | 3.03 | enriched | 3.16 | 3.37 | enriched | 3.61 | 3.56 | enriched | 3.34 | 3.37 | enriched | 2.84 | 2.85 | enriched |
| tr\|B7NZD2\|B7NZD2_RABIT | 26S proteasome non-ATPa | PSMD4 | 19S | 4.18 | 4.06 | enriched | 3.33 | 3.76 | enriched | 3.51 | 3.64 | enriched | 4.08 | 4.03 | enriched | 4.02 | 3.94 | enriched | 3.62 | 3.88 | enriched |
| tr\|G1SFE0\|G1SFE0_RABIT | 26S proteasome regulatory | PSMD14 | 19S | 2.92 | 4.60 | enriched | 2.04 | 3.45 | enriched | 2.61 | 3.89 | enriched | 3.00 | 4.34 | enriched | 3.18 | 4.81 | enriched | 2.41 | 4.03 | enriched |
| tr\|G1SM51\|G1SM51_RABIT;t | 26S proteasome non-ATPa | PSMD13 | 19S | 3.36 | 2.41 | enriched | 2.67 | 2.01 | enriched | 2.99 | 2.20 | enriched | 3.45 | 2.44 | enriched | 3.35 | 2.40 | enriched | 2.78 | 2.09 | enriched |
| tr\|G1SSA2\|G1SSA2_RABIT | 26S proteasome non-ATPa | PSMD2 | 19S | 5.17 | 5.00 | enriched | 4.45 | 4.70 | enriched | 4.69 | 4.81 | enriched | 5.30 | 4.81 | enriched | 5.19 | 5.01 | enriched | 4.70 | 4.69 | enriched |
| tr\|G1SVA3\|G1SVA3_RABIT | Proteasome 26S subunit, n | PSMD11 | 19S | 4.31 | 5.01 | enriched | 3.54 | 4.07 | enriched | 3.78 | 4.78 | enriched | 4.49 | 4.88 | enriched | 4.28 | 4.92 | enriched | 3.67 | 4.77 | enriched |
| tr\|G1SVF2\|G1SVF2_RABIT | 26S proteasome non-ATPa | PSMD1 | 19S | 4.20 | 3.33 | enriched | 3.50 | 2.98 | enriched | 3.70 | 3.12 | enriched | 4.35 | 3.39 | enriched | 4.23 | 3.34 | enriched | 3.62 | 3.04 | enriched |
| tr\|G1SYV0\|G1SYV0_RABIT | 26S proteasome AAA-ATPa | PSMC2 | 19S | 3.60 | 4.40 | enriched | 2.70 | 3.99 | enriched | 2.88 | 4.09 | enriched | 3.85 | 4.61 | enriched | 3.58 | 4.01 | enriched | 2.88 | 3.94 | enriched |
| tr\|G1T3S1\|G1T3S1_RABIT | Proteasome 26S subunit, A | PSMC6 | 19S | 3.83 | 4.60 | enriched | 2.87 | 4.05 | enriched | 3.21 | 4.27 | enriched | 3.87 | 4.76 | enriched | 3.81 | 4.60 | enriched | 3.14 | 4.27 | enriched |
| tr\|G1T6D4\|G1T6D4_RABIT | Proteasome 26S subunit, n | PSMD12 | 19S | 3.78 | 2.06 | enriched | 3.18 | 1.79 | enriched | 3.43 | 1.90 | enriched | 4.12 | 2.18 | enriched | 3.81 | 2.06 | enriched | 3.45 | 1.90 | enriched |
| tr\|G1TLQ8\|G1TLQ8_RABIT | Proteasome 26S subunit, A | PSMC3 | 19S | 3.78 | 3.63 | enriched | 3.21 | 3.29 | enriched | 3.33 | 3.42 | enriched | 3.95 | 3.70 | enriched | 3.72 | 3.59 | enriched | 3.18 | 3.34 | enriched |
| tr\|G1TP15\|G1TP15_RABIT | Proteasome 26S subunit, n | PSMD3 | 19S | 4.40 | 3.19 | enriched | 3.77 | 2.94 | enriched | 4.11 | 3.08 | enriched | 4.65 | 3.28 | enriched | 4.59 | 3.26 | enriched | 3.90 | 2.95 | enriched |
| tr\|G1U115\|G1U115_RABIT | 26S proteasome non-ATPa | PSMD6 | 19S | 4.52 | 4.01 | enriched | 3.84 | 3.73 | enriched | 4.06 | 3.80 | enriched | 4.69 | 4.07 | enriched | 4.68 | 4.07 | enriched | 3.98 | 3.73 | enriched |
| tr\|G1SVT4\|G1SVT4_RABIT | Proteasome 26S subunit, n | PSMD7 | 19S | 2.12 | 2.15 | enriched | 0.34 | 0.10 | | 1.46 | 1.61 | | 2.12 | 2.16 | enriched | 0.97 | 0.42 | | 0.24 | 0.14 | |
| tr\|G1SQL0\|G1SQL0_RABIT | AAA domain-containing pro | PSMC1 | 19S | 4.33 | 4.38 | enriched | 3.68 | 4.02 | enriched | 3.84 | 3.99 | enriched | 4.57 | 4.47 | enriched | 4.31 | 4.16 | enriched | 3.72 | 3.53 | enriched |
| tr\|G1SYM9\|G1SYM9_RABIT | Proteasome maturation pro | POMP | assembly chape | 3.35 | 3.19 | enriched | 5.17 | 3.78 | enriched | 2.23 | 2.53 | enriched | 2.12 | 2.00 | enriched | 6.04 | 4.17 | enriched | 0.27 | 0.13 | |
| tr\|B7NZA6\|B7NZA6_RABIT;tr | Proteasome assembly chap | PSMG1 | assembly chape | 4.09 | 3.47 | enriched | 5.70 | 4.02 | enriched | 3.70 | 3.30 | enriched | 4.43 | 3.59 | enriched | 6.79 | 4.34 | enriched | 3.82 | 3.32 | enriched |
| tr\|G1SN64\|G1SN64_RABIT | Proteasome assembly chap | PSMG2 | assembly chape | 2.84 | 3.71 | enriched | 4.41 | 5.77 | enriched | 2.26 | 4.61 | enriched | 3.02 | 4.97 | enriched | 5.32 | 6.04 | enriched | 2.27 | 3.44 | enriched |
| tr\|G1TCA3\|G1TCA3_RABIT | Proteasome inhibitor PI31 | PSMF1 | assembly chape | 4.40 | 2.28 | enriched | 4.68 | 2.38 | enriched | 4.47 | 2.31 | enriched | 5.58 | 2.67 | enriched | 4.40 | 2.28 | enriched | 4.35 | 2.26 | enriched |
| tr\|G1U8C4\|G1U8C4_RABIT | Proteasome activator subu | PSME1 | assembly chape | 4.51 | 5.16 | enriched | 3.96 | 4.53 | enriched | 4.21 | 5.12 | enriched | 5.18 | 5.56 | enriched | 4.26 | 4.89 | enriched | 4.55 | 3.99 | enriched |
| tr\|G1SCY3\|G1SCY3_RABIT | Ubiquitin protein ligase E3 | UBR4 | | -0.20 | 1.40 | | -0.28 | 1.84 | | -0.13 | 0.93 | | -0.27 | 1.77 | | -0.34 | 2.13 | | -0.08 | 0.54 | |
| sp\|B7NZQ9\|UBL4A_RABIT | Ubiquitin-like protein 4A | UBL4A | | 0.00 | 0.00 | | 1.22 | 1.36 | | 1.27 | 1.40 | | 0.00 | 0.00 | | 0.00 | 0.00 | | 0.00 | 0.00 | |
| sp\|G1SRW8\|SLN14_RABIT | Protein SLFN14 [Cleaved in | SLFN14 | | -0.57 | 3.50 | | -0.48 | 2.47 | | -0.53 | 3.22 | | -0.51 | 3.47 | | -0.54 | 2.50 | | -0.43 | 2.05 | |
| sp\|O19048\|PCBP1_RABIT | Poly(rC)-binding protein 1 ( | PCBP1 | | -1.69 | 4.55 | enriched | -2.57 | 4.99 | enriched | -2.61 | 4.71 | enriched | -2.66 | 5.13 | enriched | -3.05 | 4.41 | enriched | -3.18 | 5.26 | enriched |
| sp\|O19049\|HNRPK_RABIT;tr\| | Heterogeneous nuclear ribo | HNRNPK HNRPK | | 1.35 | 1.77 | | 1.39 | 1.86 | | 1.94 | 2.34 | enriched | 0.99 | 1.28 | | 0.77 | 0.84 | | 1.69 | 2.05 | enriched |
| sp\|O77622\|TCPZ_RABIT | T-complex protein 1 subuni | CCT6 | | 0.90 | 0.93 | | 2.63 | 2.81 | enriched | 2.28 | 2.77 | enriched | 1.68 | 2.23 | enriched | 1.56 | 2.16 | enriched | 1.32 | 1.70 | |
| sp\|P00563\|KCRM_RABIT;tr\|G | Creatine kinase M-type (EC | CKM | | 1.54 | 0.91 | | 1.40 | 0.82 | | 1.32 | 0.77 | | 0.77 | 0.41 | | 1.08 | 0.61 | | 1.96 | 1.15 | enriched |
| sp\|P00919\|CAH2_RABIT | Carbonic anhydrase 2 (EC 4 | CA2 | | 0.51 | 0.25 | | 0.81 | 0.42 | | 0.64 | 0.32 | | 0.59 | 0.29 | | 0.54 | 0.26 | | 0.60 | 0.30 | |
| tr\|G8ZF10\|G8ZF10_RABIT;sp\| | Alpha-globin 1 | HBA1 | | 2.29 | 1.35 | enriched | 1.87 | 1.52 | enriched | 1.73 | 1.51 | enriched | 1.72 | 1.47 | enriched | 1.30 | 1.01 | | 1.11 | 0.90 | |
| tr\|B8K174\|B8K174_RABIT;sp\| | Hemoglobin, beta (Predicte | HBB | | 1.68 | 3.06 | enriched | 1.46 | 2.97 | enriched | 1.42 | 3.67 | enriched | 1.53 | 4.71 | enriched | 1.09 | 4.03 | | 1.39 | 3.25 | |
| sp\|P12530\|LOX15_RABIT;tr\|G | Polyunsaturated fatty acid | ALOX15 | | -0.41 | 1.37 | | -1.23 | 2.99 | | -0.96 | 2.72 | | -0.87 | 2.20 | | -1.63 | 3.51 | enriched | -1.33 | 3.24 | |
| sp\|P19943\|RLA2_RABIT | 60S acidic ribosomal prote | RPLP2 | | -0.57 | 1.77 | | -0.23 | 0.78 | | -0.24 | 0.80 | | -0.33 | 0.72 | | -0.38 | 1.22 | | -0.09 | 0.20 | |
| sp\|P20618\|PSB1_HUMAN | Proteasome subunit beta t | PSMB1 PSC5 | | 6.02 | 3.64 | enriched | 1.33 | 1.30 | | 1.32 | 1.29 | | 2.00 | 1.84 | enriched | 2.24 | 1.98 | enriched | 1.26 | 1.22 | |
| tr\|G1T9L4\|G1T9L4_RABIT;sp\| | CAP18_C domain-containin | CAMP | | -0.60 | 1.55 | | -0.30 | 0.85 | | -1.10 | 0.61 | | -2.37 | 1.05 | enriched | -1.91 | 1.02 | enriched | -0.86 | 0.62 | |
| sp\|P26203\|P15B_RABIT;sp\|P2 | 15 kDa protein B (P15H) (P | P15H | | 0.01 | 0.01 | | 0.43 | 0.22 | | -0.15 | 0.07 | | 1.32 | 1.52 | | 1.04 | 1.11 | | 0.73 | 0.43 | |
| sp\|P29562\|IF4A1_RABIT;tr\|G | Eukaryotic initiation factor | EIF4A1 DDX2A EIF4A | | 0.90 | 1.56 | | 0.85 | 1.54 | | 0.78 | 1.26 | | -0.04 | 0.02 | | -0.15 | 0.12 | | -0.18 | 0.12 | |
| sp\|P30946\|HS90A_RABIT;tr\|A | Heat shock protein HSP 90- | HSP90AA1 HSPCA | | -0.19 | 0.58 | | 0.97 | 2.61 | | 0.98 | 2.59 | | -0.16 | 0.51 | | -0.21 | 0.69 | | -0.21 | 0.67 | |
| sp\|P30947\|HS90B_RABIT | Heat shock protein HSP 90- | HSP90AB1 HSPCB | | -0.49 | 1.77 | | 0.72 | 2.41 | | 0.81 | 2.38 | | -0.37 | 1.09 | | -0.41 | 1.57 | | -0.17 | 0.56 | |
| sp\|P41035\|IF2B_RABIT | Eukaryotic translation initia | EIF2S2 EIF2B | | -0.64 | 0.28 | | -1.83 | 0.96 | | -0.81 | 0.30 | | -1.24 | 0.66 | | -0.77 | 0.36 | | -0.01 | 0.01 | |
| tr\|Q0QEU3\|Q0QEU3_RABIT;t | Glyceraldehyde-3-phosphat | GAPDH | | -3.14 | 1.57 | enriched | -2.13 | 2.05 | enriched | -2.73 | 1.78 | enriched | -3.13 | 2.43 | enriched | -2.15 | 1.74 | enriched | -1.37 | 2.39 | |
| tr\|G1T1Q2\|G1T1Q2_RABIT;sp | Calmodulin | CALM | | -0.18 | 0.51 | | -0.37 | 1.14 | | -0.60 | 1.47 | | -0.46 | 1.37 | | -0.32 | 0.87 | | -0.33 | 0.98 | |
| tr\|G1U6W1\|G1U6W1_RABIT; | Elongation factor 1-alpha | EEF1A1 | | 0.56 | 0.15 | | 0.93 | 0.26 | | 0.86 | 0.23 | | 0.61 | 0.16 | | 1.10 | 0.31 | | 0.91 | 0.25 | |
| sp\|Q28618\|YBOX1_RABIT | Y-box-binding protein 1 (YE | YBX1 YB1 | | -0.40 | 2.68 | | -0.37 | 1.90 | | -0.46 | 2.64 | | -0.52 | 1.74 | | -0.06 | 0.35 | | -0.10 | 0.53 | |
| tr\|A0A140TAV6\|A0A140TAV6 | Hemoglobin subunit gamma | | | -0.39 | 1.48 | | -0.40 | 1.49 | | -0.41 | 1.41 | | -0.58 | 1.99 | | -0.76 | 2.32 | | -0.31 | 1.17 | |
| tr\|B2ZDY6\|B2ZDY6_RABIT;tr\| | CCT-alpha (T-complex prot | TCP1 | | -0.63 | 0.67 | | 0.94 | 1.59 | | 0.45 | 0.58 | | -0.38 | 0.58 | | -1.04 | 1.75 | | -0.37 | 0.62 | |
| tr\|B6V9S8\|B6V9S8_RABIT;tr\| | T-complex protein 1 subuni | CCT7 | | 0.18 | 0.16 | | 1.89 | 2.57 | enriched | 1.23 | 2.09 | | 0.67 | 1.12 | | -0.63 | 0.42 | | 0.00 | 0.00 | |
| tr\|B6V9S9\|B6V9S9_RABIT | CCT-beta (T-complex prote | CCT2 | | -1.13 | 5.30 | | -0.05 | 0.50 | | -0.51 | 3.06 | | -1.09 | 3.75 | | -1.12 | 3.21 | | -1.07 | 3.38 | |
| B7NZQ2 | Ribosomal_L16 domain-con | RPL10 | | -0.30 | 0.87 | | -0.36 | 1.05 | | -0.64 | 1.19 | | -1.20 | 0.85 | | -1.86 | 1.06 | + | -0.33 | 0.93 | |
| tr\|G1TQF7\|G1TQF7_RABIT;tr | TPR_REGION domain-containing protein | | | -0.57 | 2.14 | | -0.41 | 1.68 | | -0.05 | 0.14 | | -0.57 | 2.04 | | -0.19 | 0.71 | | -0.04 | 0.14 | |
| tr\|G1U473\|G1U473_RABIT;tr | 60S ribosomal protein L17 | RPL17 | | 0.16 | 0.06 | | 0.02 | 0.01 | | -0.12 | 0.04 | | 0.07 | 0.02 | | 0.05 | 0.02 | | 0.17 | 0.06 | |
| tr\|G1SCN8\|G1SCN8_RABIT;tr | T-complex protein 1 subuni | CCT3 | | 0.30 | 0.08 | | 2.33 | 1.11 | enriched | 1.83 | 0.85 | | 1.32 | 0.60 | | 1.23 | 0.54 | | 0.60 | 0.22 | |
| tr\|G1SCV4\|G1SCV4_RABIT | DnaJ heat shock protein fa | DNAJA4 | | -0.64 | 2.58 | | -0.46 | 1.98 | | -0.08 | 0.37 | | -0.52 | 2.15 | | -0.32 | 1.32 | | -0.25 | 1.33 | |
| tr\|G1SD04\|G1SD04_RABIT | NudE neurodevelopment pr | NDE1 | | -1.30 | 0.82 | | -1.58 | 0.93 | | -1.03 | 0.38 | | -2.35 | 1.63 | enriched | -0.90 | 0.37 | | -0.85 | 0.56 | |
| tr\|G1SDC7\|G1SDC7_RABIT | Replication protein A3 | RPA3 | | -0.25 | 0.66 | | -1.04 | 0.57 | | -0.35 | 0.97 | | 0.01 | 0.02 | | -0.37 | 0.69 | | -0.34 | 0.57 | |
| tr\|G1SDF2\|G1SDF2_RABIT | Protein-L-isoaspartate O-m | PCMT1 | | 0.01 | 0.03 | | -0.13 | 0.56 | | -0.15 | 0.51 | | -0.10 | 0.48 | | -0.07 | 0.26 | | 0.20 | 1.00 | |
| tr\|G1SDN4\|G1SDN4_RABIT;tr | Synaptotagmin binding cytoplasmic RNA interacting protein | | | 0.68 | 0.44 | | 0.08 | 0.04 | | 1.57 | 1.17 | | 0.36 | 0.14 | | 0.46 | 0.18 | | 0.54 | 0.29 | |
| tr\|G1TJT7\|G1TJT7_RABIT;tr\|( | Ribosomal_L23eN domain-containing protein | | | -0.67 | 3.39 | | -0.64 | 4.53 | | -1.70 | 1.04 | | -3.93 | 4.62 | enriched | -2.76 | 1.46 | enriched | -3.33 | 4.17 | enriched |
| tr\|G1T0C1\|G1T0C1_RABIT;tr\| | Ribosomal protein L15 | | | -0.64 | 3.31 | | -0.63 | 3.08 | | -0.55 | 1.91 | | -0.81 | 3.57 | | -0.72 | 3.18 | | -2.40 | 1.21 | enriched |
| tr\|G1SFK3\|G1SFK3_RABIT | Replication protein A subur | RPA1 | | -1.95 | 5.99 | enriched | -1.77 | 5.52 | enriched | -1.98 | 4.63 | enriched | -1.59 | 5.71 | enriched | -1.56 | 5.43 | enriched | -1.45 | 5.23 | enriched |

| | | | | | | | | | | | | | | | | | | | | | |
|---|---|---|---|---|---|---|---|---|---|---|---|---|---|---|---|---|---|---|---|---|---|
| tr\|G1SFR8\|G1SFR8_RABIT | 40S ribosomal protein S12 | | | 0.25 | 0.22 | | -0.10 | 0.07 | | -0.08 | 0.06 | | 0.03 | 0.02 | | -0.07 | 0.06 | | 0.08 | 0.06 | |
| tr\|G1SFV7\|G1SFV7_RABIT | DNA damage-binding prote | DDB1 | | 0.17 | 0.53 | | 0.31 | 1.21 | | 0.41 | 1.54 | | 0.18 | 0.61 | | -0.01 | 0.04 | | 0.10 | 0.38 | |
| tr\|G1SG29\|G1SG29_RABIT | Skp1_POZ domain-containing protein | | | 1.00 | 0.68 | | 0.72 | 0.47 | | 4.00 | 2.44 | enriched | 1.10 | 0.78 | | -1.15 | 0.38 | | 0.38 | 0.12 | |
| tr\|G1SGX4\|G1SGX4_RABIT | Uncharacterized protein | | | 1.01 | 0.62 | | 1.17 | 0.72 | | -0.66 | 0.27 | | 0.85 | 0.51 | | 0.76 | 0.45 | | 0.06 | 0.02 | |
| tr\|G1TSV2\|G1TSV2_RABIT;tr | 60S ribosomal protein L21 | RPL21 | | 0.10 | 0.04 | | -0.19 | 0.13 | | -0.18 | 0.08 | | 0.05 | 0.03 | | 0.38 | 0.46 | | 0.10 | 0.13 | |
| tr\|G1SHR7\|G1SHR7_RABIT | Potassium channel modulatory factor 1 | | | 2.53 | 2.13 | enriched | 1.81 | 1.19 | enriched | 2.30 | 1.88 | enriched | 2.15 | 1.86 | enriched | 2.01 | 1.71 | enriched | 2.55 | 2.13 | enriched |
| tr\|G1U6S0\|G1U6S0_RABIT;tr | Uncharacterized protein | | | 0.26 | 0.17 | | -0.79 | 0.28 | | -0.72 | 0.28 | | -0.90 | 0.31 | | 0.02 | 0.01 | | 0.39 | 0.25 | |
| tr\|G1SHW8\|G1SHW8_RABIT | Keratin 85 | KRT85 | | 0.00 | 0.00 | | 3.56 | 3.75 | enriched | 0.00 | 0.00 | | 0.00 | 0.00 | | 0.00 | 0.00 | | 0.00 | 0.00 | |
| tr\|G1SHY2\|G1SHY2_RABIT | IF rod domain-containing protein | | | 0.00 | 0.00 | | 5.40 | 3.70 | enriched | 0.00 | 0.00 | | 0.00 | 0.00 | | 0.00 | 0.00 | | 0.00 | 0.00 | |
| tr\|G1SHZ8\|G1SHZ8_RABIT | CCT-theta (T-complex prote | CCT8 | | -0.81 | 2.72 | | 0.78 | 3.01 | | 0.19 | 1.09 | | -0.49 | 1.82 | | -0.57 | 2.20 | | -0.58 | 2.34 | |
| tr\|G1SIB6\|G1SIB6_RABIT | Biliverdin reductase B | BLVRB | | 0.11 | 0.04 | | 1.37 | 1.29 | | 0.78 | 0.51 | | -0.24 | 0.09 | | 0.00 | 0.00 | | 1.65 | 1.60 | enriched |
| tr\|G1SID3\|G1SID3_RABIT | Vacuolar protein sorting 13 | VPS13C | | -0.31 | 2.07 | | -0.18 | 1.26 | | -0.41 | 2.05 | | -0.09 | 0.60 | | -0.13 | 0.91 | | -0.14 | 0.96 | |
| tr\|G1SIF7\|G1SIF7_RABIT | DDB1- and CUL4-associated factor 11 | | | 1.79 | 2.46 | enriched | 1.79 | 1.95 | enriched | 1.97 | 2.50 | enriched | 0.00 | 0.00 | | -0.05 | 0.01 | | 0.00 | 0.00 | |
| tr\|G1SIZ2\|G1SIZ2_RABIT;tr\|G | 40S ribosomal protein S20 | RPS20 | | -0.80 | 1.55 | | -0.56 | 2.72 | | -0.65 | 1.38 | | -0.80 | 1.53 | | -0.52 | 0.88 | | -0.15 | 0.69 | |
| tr\|G1SJB4\|G1SJB4_RABIT;tr\|( | WD_REPEATS_REGION domain-containing protein | | | -0.48 | 1.96 | | -0.37 | 1.51 | | -0.73 | 2.64 | | -0.58 | 1.90 | | -0.62 | 2.38 | | -0.34 | 1.46 | |
| tr\|G1SJQ9\|G1SJQ9_RABIT | HECT domain E3 ubiquitin p | HECTD3 | | 1.34 | 1.85 | | 1.11 | 1.44 | | 0.45 | 0.25 | | 0.00 | 0.00 | | 0.38 | 0.32 | | 0.00 | 0.00 | |
| tr\|G1SK22\|G1SK22_RABIT;sp | 40S ribosomal protein S27a (Ubiquitin carboxyl extension prot | | | 0.55 | 1.97 | | 0.60 | 2.42 | | 0.90 | 2.91 | | 1.02 | 2.83 | | 0.82 | 2.78 | | 0.43 | 1.97 | |
| tr\|G1TG04\|G1TG04_RABIT;tr | 60S ribosomal protein L6 | | | -0.61 | 2.26 | | -0.51 | 3.28 | | -0.68 | 2.15 | | -0.58 | 3.05 | | -0.46 | 1.54 | | -0.38 | 2.78 | |
| tr\|G1SKP2\|G1SKP2_RABIT | Importin 5 | IPO5 | | 0.45 | 2.32 | | 0.12 | 0.47 | | -0.12 | 0.22 | | 0.13 | 0.46 | | -0.76 | 0.60 | | -0.04 | 0.14 | |
| tr\|G1SKW5\|G1SKW5_RABIT | Deleted. | | | 4.26 | 5.27 | enriched | 4.22 | 5.40 | enriched | 3.84 | 5.04 | enriched | 4.64 | 5.44 | enriched | 5.23 | 5.60 | enriched | 4.55 | 5.43 | enriched |
| tr\|G1TY53\|G1TY53_RABIT;tr\| | Ribosomal protein | | | 1.47 | 0.91 | | 1.56 | 0.97 | | 1.60 | 0.99 | | 1.47 | 0.90 | | 1.32 | 0.80 | | 1.17 | 0.70 | |
| tr\|G1SLA0\|G1SLA0_RABIT;tr\| | Uncharacterized protein | | | 1.86 | 1.15 | enriched | 1.97 | 1.23 | enriched | 1.08 | 0.39 | | 2.06 | 1.27 | enriched | 1.67 | 0.78 | | 0.84 | 0.36 | |
| tr\|G1SLK2\|G1SLK2_RABIT;tr\| | AAA domain-containing protein | | | 3.55 | 2.51 | enriched | 2.90 | 2.20 | enriched | 3.10 | 2.31 | enriched | 3.72 | 2.55 | enriched | 3.36 | 2.43 | enriched | 2.88 | 2.19 | enriched |
| tr\|G1TUR3\|G1TUR3_RABIT;tr | Polyadenylate-binding protein (PABP) | | | -0.50 | 1.38 | | -0.39 | 1.05 | | -0.15 | 0.39 | | -0.51 | 1.37 | | -0.39 | 1.00 | | -0.19 | 0.50 | |
| tr\|G1SLM1\|G1SLM1_RABIT | Calcyclin-binding protein | RABGAP1L | | 0.00 | 0.00 | | 1.36 | 0.64 | | 0.00 | 0.00 | | -0.32 | 0.12 | | 0.00 | 0.00 | | 0.00 | 0.00 | |
| tr\|G1SLT8\|G1SLT8_RABIT | Heterogeneous nuclear ribo | HNRNPH3 | | 0.06 | 0.04 | | 0.79 | 1.65 | | 0.57 | 1.32 | | 0.60 | 1.40 | | 0.44 | 0.42 | | 0.90 | 1.96 | |
| tr\|G1SLW3\|G1SLW3_RABIT | Uncharacterized protein | | | 0.00 | 0.00 | | 0.40 | 0.24 | | 0.56 | 0.35 | | 0.37 | 0.22 | | 0.41 | 0.25 | | 0.63 | 0.41 | |
| tr\|G1SMS1\|G1SMS1_RABIT;tr | 26S proteasome non-ATPase regulatory subunit 13 (26S prote | | | 3.36 | 2.41 | enriched | 2.67 | 2.01 | enriched | 2.99 | 2.20 | enriched | 3.45 | 2.44 | enriched | 3.35 | 2.40 | enriched | 2.78 | 2.09 | enriched |
| tr\|G1SMC8\|G1SMC8_RABIT;tr | Uncharacterized protein | | | 1.11 | 1.28 | | -0.15 | 0.12 | | 0.10 | 0.05 | | 1.27 | 1.49 | | 0.73 | 0.78 | | 0.53 | 0.45 | |
| tr\|G1SMK8\|G1SMK8_RABIT | Deleted. | | | 1.05 | 0.42 | | 1.46 | 0.62 | | 1.24 | 0.51 | | 1.32 | 0.55 | | 1.27 | 0.52 | | 1.72 | 0.74 | |
| tr\|G1SMM5\|G1SMM5_RABIT | DnaJ heat shock protein fa | DNAJA2 | | -0.35 | 1.68 | | -0.01 | 0.03 | | 0.13 | 0.67 | | -0.10 | 0.42 | | 0.01 | 0.02 | | -0.17 | 0.85 | |
| tr\|G1SMR7\|G1SMR7_RABIT | 60S ribosomal protein L12 | | | -0.55 | 2.91 | | -0.58 | 1.85 | | -0.64 | 3.00 | | -0.76 | 3.35 | | -0.69 | 2.10 | | -0.44 | 2.75 | |
| tr\|U3KML6\|U3KML6_RABIT;tr | Deleted. | | | -0.75 | 2.77 | | -0.50 | 2.47 | | -0.70 | 3.03 | | -0.72 | 3.20 | | -0.68 | 2.71 | | -0.20 | 1.04 | |
| tr\|G1SP51\|G1SP51_RABIT | 40S ribosomal protein S13 | RPS13 | | 0.39 | 0.25 | | 0.69 | 0.73 | | 0.49 | 0.49 | | -0.22 | 0.12 | | 0.21 | 0.14 | | 1.20 | 1.36 | |
| tr\|G1SPB8\|G1SPB8_RABIT | Nucleosome assembly prot | NAP1L4 | | 2.22 | 3.93 | enriched | 0.70 | 0.53 | | 1.64 | 3.00 | enriched | 0.00 | 0.00 | | 0.00 | 0.00 | | 0.00 | 0.00 | |
| tr\|G1SPS8\|G1SPS8_RABIT | E74 like ETS transcription f | ELF1 | | 0.00 | 0.00 | | 1.99 | 2.78 | enriched | 0.00 | 0.00 | | 0.00 | 0.00 | | 0.00 | 0.00 | | 0.00 | 0.00 | |
| tr\|G1SPU6\|G1SPU6_RABIT | Uncharacterized protein | | | 1.08 | 1.08 | | 1.97 | 1.54 | enriched | 1.40 | 1.16 | | -0.04 | 0.03 | | -0.06 | 0.05 | | 0.13 | 0.10 | |
| tr\|G1T569\|G1T569_RABIT;tr\| | Uncharacterized protein | | | -1.21 | 2.75 | | -0.83 | 1.90 | | -1.13 | 2.05 | | -1.05 | 3.10 | | -1.05 | 3.16 | | -0.40 | 1.42 | |
| tr\|G1SQ96\|G1SQ96_RABIT;tr | Protein phosphatase 6 regu | PPP6R3 | | -0.46 | 3.16 | | -0.21 | 1.39 | | -0.41 | 3.61 | | -0.47 | 2.83 | | -0.39 | 2.30 | | -0.48 | 2.54 | |
| tr\|G1SQL0\|G1SQL0_RABIT | AAA domain-containing protein | | | 4.33 | 4.38 | enriched | 3.68 | 4.02 | enriched | 3.84 | 3.99 | enriched | 4.57 | 4.47 | enriched | 4.31 | 4.16 | enriched | 3.72 | 3.53 | enriched |
| tr\|G1SQZ4\|G1SQZ4_RABIT | RuvB-like helicase (EC 3.6.4 | RUVBL2 | | -0.47 | 2.33 | | -0.41 | 2.23 | | -0.70 | 2.98 | | -0.47 | 2.42 | | -0.37 | 1.89 | | -0.24 | 1.34 | |
| tr\|G1SR03\|G1SR03_RABIT | 15S Mg(2+)-ATPase p97 su | VCP | | -0.08 | 0.59 | | -0.25 | 2.06 | | -0.14 | 1.21 | | 0.23 | 1.57 | | -0.02 | 0.14 | | 0.32 | 2.15 | |
| tr\|G1SR38\|G1SR38_RABIT;tr\| | IF rod domain-containing protein | | | 0.00 | 0.00 | | 7.56 | 3.61 | enriched | 0.00 | 0.00 | | -0.08 | 0.04 | | 0.65 | 0.41 | | 0.00 | 0.00 | |
| tr\|G1SRA8\|G1SRA8_RABIT | Protein-synthesizing GTPase (EC 3.6.5.3) | | | 0.48 | 0.43 | | -0.25 | 0.13 | | -0.40 | 0.19 | | -0.38 | 0.31 | | -0.21 | 0.09 | | 0.55 | 0.31 | |
| tr\|G1SRF5\|G1SRF5_RABIT | Protein phosphatase 6 regu | PPP6R1 | | 1.44 | 1.20 | | 2.21 | 1.79 | enriched | 1.92 | 1.52 | enriched | 0.00 | 0.00 | | 0.00 | 0.00 | | 0.00 | 0.00 | |
| tr\|G1SRF7\|G1SRF7_RABIT | 75 kDa glucose-regulated p | HSPA9 | | -0.40 | 1.83 | | -1.52 | 0.73 | | -2.27 | 1.27 | enriched | -1.84 | 0.59 | | -0.41 | 1.60 | | -0.15 | 0.65 | |
| tr\|U3KPK3\|U3KPK3_RABIT;tr\| | Diaphanous related formin | DIAPH2 | | 4.20 | 1.10 | enriched | 6.40 | 4.30 | enriched | 4.21 | 0.96 | enriched | 6.26 | 4.14 | enriched | 2.24 | 0.43 | | 6.42 | 4.30 | enriched |
| tr\|G1TPV0\|G1TPV0_RABIT;tr\| | Uncharacterized protein | | | -0.42 | 1.88 | | -0.41 | 2.16 | | -3.10 | 1.11 | enriched | -4.05 | 4.31 | enriched | -0.57 | 2.65 | | -2.03 | 1.10 | enriched |
| tr\|G1STW0\|G1STW0_RABIT | 60S ribosomal protein L7a | | | -0.64 | 4.98 | | -0.59 | 4.80 | | -0.43 | 1.95 | | -0.51 | 2.00 | | -0.53 | 3.39 | | -0.33 | 2.42 | |
| tr\|G1SU30\|G1SU30_RABIT | BCL2-associated athanogen | BAG6 | | 0.34 | 0.21 | | 2.16 | 1.67 | enriched | 2.62 | 1.92 | enriched | -0.70 | 0.42 | | -1.02 | 0.55 | | -1.87 | 1.26 | enriched |
| tr\|G1TRC0\|G1TRC0_RABIT;tr\| | Uncharacterized protein | | | -0.16 | 1.55 | | 0.01 | 0.12 | | -0.05 | 0.63 | | -0.02 | 0.20 | | 0.18 | 1.89 | | 0.25 | 2.04 | |
| tr\|G1TUB1\|G1TUB1_RABIT;tr | Uncharacterized protein | | | 0.27 | 0.06 | | -0.42 | 0.08 | | 0.72 | 0.19 | | 0.57 | 0.15 | | 0.66 | 0.18 | | 0.77 | 0.21 | |
| tr\|G1SVB0\|G1SVB0_RABIT;tr | 40S ribosomal protein S7 | | | 0.50 | 0.96 | | 0.00 | 0.00 | | 0.25 | 0.14 | | 0.11 | 0.24 | | 0.01 | 0.01 | | 0.37 | 0.40 | |
| tr\|G1SVE6\|G1SVE6_RABIT | AD domain-containing protein | | | -2.56 | 4.59 | enriched | -1.98 | 1.51 | enriched | -1.26 | 2.28 | | -3.01 | 4.48 | enriched | -1.15 | 1.22 | | -1.23 | 0.77 | |
| tr\|G1SVR2\|G1SVR2_RABIT | YTH N6-methyladenosine R | YTHDF3 | | 0.48 | 0.23 | | -1.29 | 0.59 | | -0.88 | 0.36 | | -0.84 | 0.38 | | 0.16 | 0.05 | | -0.88 | 0.42 | |
| tr\|G1SVW5\|G1SVW5_RABIT | 60S ribosomal protein L4 | | | -0.49 | 2.28 | | -0.67 | 3.34 | | -0.67 | 3.29 | | -0.50 | 1.85 | | -0.57 | 1.93 | | -0.23 | 1.06 | |
| tr\|G1SW76\|G1SW76_RABIT | Calcium binding and coiled | CALCOCO1 | | -1.16 | 0.43 | | -0.93 | 0.29 | | -0.67 | 0.22 | | -0.66 | 0.19 | | -0.40 | 0.12 | | 0.35 | 0.12 | |
| tr\|G1SWS0\|G1SWS0_RABIT;t | IF rod domain-containing protein | | | 0.00 | 0.00 | | 5.90 | 4.52 | enriched | 0.00 | 0.00 | | 0.00 | 0.00 | | 0.00 | 0.00 | | 0.00 | 0.00 | |
| tr\|G1SX32\|G1SX32_RABIT | Serine/threonine-protein p | PPP6C | | 1.33 | 1.73 | | 1.80 | 2.15 | enriched | 1.47 | 1.81 | | 1.32 | 1.75 | | 1.06 | 1.14 | | 0.57 | 0.33 | |
| tr\|G1SY58\|G1SY58_RABIT | Iron-responsive element-bi | IREB2 | | -1.13 | 0.94 | | -0.34 | 0.62 | | -0.89 | 0.51 | | -0.64 | 1.12 | | -0.15 | 0.24 | | 0.08 | 0.12 | |
| tr\|G1SYJ6\|G1SYJ6_RABIT;tr\|G | Ribosomal_L18_c domain-containing protein | | | -0.45 | 2.64 | | -0.45 | 2.26 | | -0.69 | 2.69 | | -0.83 | 2.34 | | -0.51 | 2.22 | | -0.46 | 1.79 | |
| tr\|G1SYL8\|G1SYL8_RABIT | Elongin B | ELOB | | 0.04 | 0.02 | | -0.61 | 0.43 | | 4.14 | 2.89 | enriched | -0.10 | 0.06 | | 0.46 | 0.34 | | 0.50 | 0.36 | |
| tr\|G1SYR7\|G1SYR7_RABIT | RING-type E3 ubiquitin tra | MKRN1 | | -0.68 | 2.92 | | -0.61 | 2.74 | | -0.41 | 1.79 | | -0.64 | 2.95 | | -0.63 | 3.08 | | -0.18 | 0.92 | |
| tr\|G1SYV0\|G1SYV0_RABIT | 26S proteasome AAA-ATPa | PSMC2 | | 3.60 | 4.40 | enriched | 2.70 | 3.99 | enriched | 2.88 | 4.09 | enriched | 3.85 | 4.61 | enriched | 3.58 | 4.01 | enriched | 2.88 | 3.94 | enriched |
| tr\|U3KNW6\|U3KNW6_RABIT; | 60S ribosomal protein L14 | RPL14 | | 0.00 | 0.00 | | 0.03 | 0.08 | | 0.09 | 0.11 | | -0.29 | 0.82 | | -0.08 | 0.25 | | -0.07 | 0.25 | |
| tr\|G1SZT8\|G1SZT8_RABIT | GATOR complex protein SE | SEC13 | | 0.45 | 0.33 | | 0.59 | 0.45 | | -0.61 | 0.31 | | 0.45 | 0.32 | | 0.71 | 0.53 | | 0.85 | 0.69 | |
| tr\|G1TPV3\|G1TPV3_RABIT;tr\| | S10_plectin domain-containing protein | | | 0.43 | 1.09 | | 0.74 | 1.49 | | -0.13 | 0.16 | | 0.63 | 0.72 | | 0.93 | 1.46 | | 0.00 | 0.00 | |
| tr\|G1T1C5\|G1T1C5_RABIT | Uncharacterized protein | | | -0.90 | 0.69 | | -0.61 | 0.65 | | -0.42 | 0.43 | | -0.83 | 0.61 | | -0.73 | 0.67 | | 0.14 | 0.12 | |
| tr\|G1U472\|G1U472_RABIT;tr | Uncharacterized protein | | | 2.77 | 2.51 | enriched | 2.71 | 2.49 | enriched | 2.51 | 2.35 | enriched | 2.42 | 2.26 | enriched | 2.59 | 2.39 | enriched | 2.75 | 2.48 | enriched |
| tr\|G1T1V9\|G1T1V9_RABIT | Heat shock protein family A | HSPA2 | | -0.11 | 0.15 | | -0.42 | 0.27 | | 0.09 | 0.14 | | -0.74 | 0.49 | | -0.25 | 0.27 | | 0.40 | 0.74 | |
| tr\|G1T2L1\|G1T2L1_RABIT | Proteasome subunit alpha type | | | 4.67 | 5.22 | enriched | 4.82 | 5.64 | enriched | 4.34 | 4.19 | enriched | 5.04 | 5.64 | enriched | 5.43 | 5.58 | enriched | 4.37 | 5.40 | enriched |
| tr\|G1T2U1\|G1T2U1_RABIT | Amidophosphoribosyltrans | PPAT | | -0.84 | 3.41 | | -0.66 | 3.42 | | -1.02 | 3.80 | | -0.70 | 2.62 | | -0.47 | 3.49 | | 0.00 | 0.01 | |
| tr\|G1T345\|G1T345_RABIT;tr\| | E3 ubiquitin-protein ligase | UBR1 | | -0.18 | 0.65 | | -0.22 | 0.79 | | -0.29 | 1.10 | | 0.25 | 0.90 | | -0.04 | 0.12 | | 0.10 | 0.31 | |
| tr\|G1T3D9\|G1T3D9_RABIT;tr\| | 40S ribosomal protein S8 | | | -0.47 | 2.78 | | -0.36 | 2.60 | | -0.51 | 3.06 | | -0.46 | 2.83 | | -0.48 | 2.16 | | -0.20 | 1.50 | |
| tr\|G1T3U5\|G1T3U5_RABIT | DnaJ heat shock protein fa | DNAJB4 | | -0.05 | 0.26 | | 0.16 | 0.93 | | 0.08 | 0.42 | | 0.22 | 1.13 | | 0.21 | 1.23 | | 0.49 | 2.52 | |
| tr\|G1T3X6\|G1T3X6_RABIT | Tripartite motif containing | TRIM58 | | 0.03 | 0.18 | | 0.02 | 0.07 | | -0.11 | 0.54 | | -0.29 | 1.88 | | -0.17 | 1.21 | | 0.01 | 0.02 | |
| tr\|G1T3Y8\|G1T3Y8_RABIT | 60 kDa chaperonin (EC 5.6.1.7) (60 kDa heat shock protein, mi | | | 0.00 | 0.00 | | 0.46 | 0.33 | | 2.75 | 2.50 | enriched | 2.83 | 2.66 | enriched | 1.27 | 0.84 | | 0.00 | 0.00 | |
| tr\|G1T4P0\|G1T4P0_RABIT | Keratin 32 | KRT32 | | 0.00 | 0.00 | | 7.14 | 4.01 | enriched | 0.00 | 0.00 | | 0.00 | 0.00 | | 0.00 | 0.00 | | 0.00 | 0.00 | |
| tr\|G1T520\|G1T520_RABIT | DnaJ heat shock protein fa | DNAJC7 | | -0.54 | 1.55 | | -0.36 | 1.85 | | -0.19 | 0.91 | | -0.40 | 1.58 | | 0.00 | 0.00 | | -0.21 | 0.75 | |
| tr\|G1U4I5\|G1U4I5_RABIT;tr\|( | 60S ribosomal protein L9 | | | 0.28 | 1.15 | | 0.08 | 0.15 | | 0.30 | 0.99 | | -0.06 | 0.15 | | 0.09 | 0.37 | | 0.19 | 0.68 | |
| tr\|G1T695\|G1T695_RABIT | Nucleosome assembly prot | NAP1L1 | | 2.26 | 1.97 | enriched | 1.97 | 1.74 | enriched | 2.62 | 2.23 | enriched | -0.08 | 0.03 | | 0.00 | 0.00 | | -0.35 | 0.13 | |
| tr\|G1T6D1\|G1T6D1_RABIT;tr\| | 60S ribosomal protein L23 | | | -0.07 | 0.03 | | 0.70 | 0.44 | | 0.57 | 0.45 | | 0.71 | 0.58 | | 0.42 | 0.25 | | 1.05 | 0.89 | |
| tr\|G1T6W5\|G1T6W5_RABIT | Keratin 34 | KRT34 | | 0.00 | 0.00 | | 7.18 | 4.96 | enriched | 0.00 | 0.00 | | 0.00 | 0.00 | | 0.00 | 0.00 | | 0.00 | 0.00 | |
| tr\|G1T6W7\|G1T6W7_RABIT | Catalase (EC 1.11.1.6) | CAT | | -0.85 | 2.89 | | -0.77 | 2.60 | | -0.57 | 2.22 | | -0.81 | 2.03 | | -0.86 | 2.58 | | -0.15 | 0.50 | |
| tr\|G1T7I3\|G1T7I3_RABIT | Acetyl-CoA carboxylase 1 (E | ACACA | | 0.45 | 3.49 | | 0.44 | 3.45 | | 0.01 | 0.11 | | 0.28 | 2.88 | | 0.09 | 0.67 | | 0.35 | 3.12 | |
| tr\|G1T7R2\|G1T7R2_RABIT | Tyrosine 3-monooxygenase | YWHAE | | 0.05 | 0.08 | | -0.12 | 0.27 | | 0.15 | 0.17 | | -0.40 | 1.24 | | 0.00 | 0.00 | | 0.00 | 0.00 | |
| tr\|U3KM36\|U3KM36_RABIT;t | Ubiquitin carboxyl-termina | USP15 | | -1.29 | 2.83 | | -0.92 | 2.52 | | -1.04 | 2.68 | | -1.15 | 2.60 | | -0.68 | 1.97 | | -0.18 | 0.49 | |
| tr\|G1T810\|G1T810_RABIT | Cullin 2 | CUL2 | | 0.38 | 0.55 | | -0.19 | 0.21 | | 0.04 | 0.04 | | 0.35 | 0.49 | | 0.39 | 0.40 | | 0.61 | 0.45 | |
| tr\|G1T874\|G1T874_RABIT | Zinc finger ZZ-type and EF- | ZZEF1 | | 1.95 | 1.33 | enriched | 1.66 | 1.15 | | 2.17 | 1.45 | enriched | -0.15 | 0.05 | | -0.33 | 0.12 | | -0.29 | 0.14 | |
| tr\|G1T8P4\|G1T8P4_RABIT | Ribosomal protein lateral s | RPLP1 | | 0.40 | 0.98 | | 0.51 | 1.19 | | 0.39 | 0.85 | | -0.13 | 0.07 | | 0.34 | 0.70 | | 0.49 | 1.02 | |
| tr\|G1T8S4\|G1T8S4_RABIT | Clustered mitochondria pro | CLUH | | 0.00 | 0.00 | | 0.00 | 0.00 | | 2.21 | 1.94 | enriched | 0.00 | 0.00 | | 0.00 | 0.00 | | 0.00 | 0.00 | |
| tr\|G1T8V4\|G1T8V4_RABIT | DnaJ heat shock protein fa | DNAJB2 | | 0.00 | 0.00 | | 1.65 | 1.47 | enriched | 1.77 | 1.54 | enriched | 0.71 | 0.30 | | 1.86 | 1.64 | enriched | 0.00 | 0.00 | |
| tr\|G1T9F3\|G1T9F3_RABIT | Karyopherin subunit beta 1 | KPNB1 | | 1.95 | 1.97 | enriched | 0.51 | 0.26 | | 0.96 | 1.06 | | 1.29 | 1.35 | | 0.79 | 0.61 | | 0.52 | 0.30 | |
| tr\|G1T9M9\|G1T9M9_RABIT | Heat shock protein family A (Hsp70) member 8 | | | -0.25 | 1.94 | | -0.08 | 0.67 | | -0.09 | 0.61 | | -0.13 | 1.01 | | 0.03 | 0.09 | | 0.14 | 0.98 | |
| tr\|G1TA47\|G1TA47_RABIT;tr\| | 40S ribosomal protein S4 | | | -0.24 | 1.32 | | -0.04 | 0.14 | | -0.25 | 1.34 | | -0.37 | 1.43 | | -0.33 | 1.09 | | -0.23 | 0.77 | |
| tr\|G1TAF8\|G1TAF8_RABIT | Stress induced phosphopro | STIP1 | | -0.62 | 2.38 | | 0.12 | 0.51 | | 0.60 | 2.57 | | -0.61 | 2.12 | | -0.63 | 2.17 | | -0.30 | 1.48 | |
| tr\|G1TB40\|G1TB40_RABIT | Replication protein A2 | RPA2 | | -2.44 | 3.42 | enriched | -3.66 | 2.72 | enriched | -2.49 | 3.42 | enriched | -2.21 | 3.88 | enriched | -3.42 | 1.42 | enriched | -3.30 | 1.92 | enriched |
| tr\|G1TBL6\|G1TBL6_RABIT | Clathrin heavy chain | CLTC | | -2.15 | 1.30 | enriched | -2.13 | 2.31 | enriched | -1.88 | 1.97 | enriched | -2.70 | 1.16 | enriched | -1.99 | 2.57 | enriched | -0.95 | 1.97 | |

| | | | | | | | | | | | | | | | | | | | | | |
|---|---|---|---|---|---|---|---|---|---|---|---|---|---|---|---|---|---|---|---|---|---|
| tr\|G1TBS2\|G1TBS2_RABIT | RuvB-like helicase (EC 3.6.4 | RUVBL1 | | -0.04 | 0.06 | | 0.17 | 0.29 | | -0.01 | 0.02 | | 0.11 | 0.18 | | 0.10 | 0.15 | | 0.17 | 0.29 | |
| tr\|G1TCB7\|G1TCB7_RABIT | Ubiquitin carboxyl-terminal | UCHL5 | | 0.94 | 1.50 | | 2.89 | 3.33 | enriched | 0.19 | 0.16 | | 1.10 | 1.84 | | 0.30 | 0.30 | | -0.26 | 0.17 | |
| tr\|G1TCS8\|G1TCS8_RABIT | RAB1A, member RAS onco | RAB1A | | 0.75 | 0.39 | | -0.29 | 0.08 | | -1.61 | 0.85 | | -0.10 | 0.03 | | 0.06 | 0.02 | | -0.79 | 0.25 | |
| tr\|G1TCT3\|G1TCT3_RABIT | Aspartate carbamoyltransf | CAD | | 6.29 | 5.71 | enriched | 6.84 | 6.33 | enriched | 8.65 | 6.62 | enriched | 6.97 | 6.32 | enriched | 6.23 | 4.81 | enriched | 0.63 | 1.51 | |
| tr\|G1TCW1\|G1TCW1_RABIT | Transferrin receptor protei | TFRC | | 1.50 | 2.26 | enriched | 1.51 | 2.29 | enriched | 1.49 | 2.17 | enriched | 1.49 | 2.10 | | 2.17 | 2.74 | enriched | 1.79 | 2.47 | enriched |
| tr\|G1TD41\|G1TD41_RABIT;tr\| | Heterogeneous nuclear ribo | HNRNPH1 | | -0.99 | 2.31 | | -0.99 | 2.73 | | -1.04 | 2.66 | | -1.23 | 3.13 | | -1.09 | 2.74 | | -0.38 | 1.38 | |
| tr\|G1TDB3\|G1TDB3_RABIT | 40S ribosomal protein S25 | | | 0.28 | 0.13 | | 0.60 | 0.30 | | 0.37 | 0.17 | | 0.39 | 0.18 | | -0.13 | 0.05 | | 0.66 | 0.34 | |
| tr\|G1TDI0\|G1TDI0_RABIT | Peptidase_M24 domain-containing protein | | | -0.92 | 0.27 | | -1.35 | 0.51 | | -0.66 | 0.20 | | -0.87 | 0.27 | | -0.85 | 0.29 | | 0.68 | 0.25 | |
| tr\|G1TDJ2\|G1TDJ2_RABIT | Mitochondrial import inner | TIMM8B | | 1.89 | 0.57 | | 1.15 | 1.61 | | 1.09 | 1.03 | | 1.58 | 1.56 | | 0.00 | 0.00 | | 0.00 | 0.00 | |
| tr\|G1U1G9\|G1U1G9_RABIT;t | 60S ribosomal protein L30 | | | -0.04 | 0.13 | | 0.02 | 0.05 | | -0.29 | 0.97 | | -0.26 | 1.28 | | -0.36 | 1.65 | | -0.07 | 0.31 | |
| tr\|G1TEU6\|G1TEU6_RABIT;tr | E3 ubiquitin-protein ligase | UBR2 | | -0.23 | 1.23 | | -0.05 | 0.20 | | -0.20 | 1.12 | | 0.28 | 1.45 | | 0.13 | 0.58 | | 0.16 | 0.84 | |
| tr\|G1TEW4\|G1TEW4_RABIT | Ubiquitinyl hydrolase 1 (EC | USP9X | | 1.67 | 2.18 | enriched | 0.86 | 1.90 | | 1.29 | 1.91 | | 0.16 | 0.17 | | 1.29 | 2.40 | | -0.24 | 0.60 | |
| tr\|G1TFM5\|G1TFM5_RABIT;t | Ribosomal protein S5 | RPS5 | | 0.43 | 1.82 | | 0.57 | 2.10 | | 0.48 | 1.30 | | 0.75 | 2.56 | | 0.48 | 1.29 | | 0.69 | 2.49 | |
| tr\|G1TFX7\|G1TFX7_RABIT | YTH N6-methyladenosine R | YTHDF2 | | -0.77 | 2.00 | | -0.47 | 3.23 | | -0.60 | 1.67 | | -0.57 | 2.20 | | -1.30 | 1.33 | | -0.17 | 0.58 | |
| tr\|G1TIB4\|G1TIB4_RABIT | 40S ribosomal protein S28 | RPS28 | | -1.11 | 0.74 | | -0.44 | 3.07 | | -1.33 | 1.00 | | -2.93 | 1.08 | enriched | -3.32 | 2.94 | enriched | -2.48 | 3.12 | enriched |
| tr\|G1TK56\|G1TK56_RABIT | Deleted. | | | 0.34 | 0.13 | | 0.59 | 0.25 | | 0.21 | 0.08 | | 0.40 | 0.16 | | 0.55 | 0.23 | | 0.57 | 0.24 | |
| tr\|G1TL06\|G1TL06_RABIT;tr\|( | Ribosomal protein L3 | RPL3 | | -1.33 | 1.14 | | -0.81 | 2.76 | | -0.67 | 2.65 | | -0.93 | 2.35 | | -0.70 | 2.51 | | -0.53 | 2.29 | |
| tr\|G1U5C0\|G1U5C0_RABIT;tr | 40S ribosomal protein S6 | | | 0.76 | 0.53 | | -0.60 | 0.22 | | 0.58 | 0.38 | | 0.73 | 0.50 | | 0.94 | 0.68 | | 0.97 | 0.70 | |
| tr\|G1TMS5\|G1TMS5_RABIT | Chaperonin containing TCP1 | CCT5 | | 0.30 | 0.15 | | 2.64 | 2.22 | enriched | 1.93 | 1.76 | enriched | 0.63 | 0.36 | | 0.59 | 0.37 | | 0.58 | 0.33 | |
| tr\|G1TMU2\|G1TMU2_RABIT | Uncharacterized protein | | | -0.23 | 1.08 | | -0.49 | 1.62 | | -0.45 | 1.57 | | -0.58 | 2.27 | | -0.44 | 1.70 | | 0.19 | 0.85 | |
| tr\|G1TN62\|G1TN62_RABIT | 40S ribosomal protein S19 | | | -0.36 | 1.96 | | -0.35 | 1.87 | | -0.42 | 2.11 | | -0.54 | 2.39 | | -0.43 | 2.15 | | -0.23 | 1.04 | |
| tr\|G1TNM3\|G1TNM3_RABIT;t | 40S ribosomal protein S3 (EC 4.2.99.18) | | | -0.65 | 0.92 | | -0.60 | 0.86 | | -0.72 | 1.02 | | -0.62 | 0.85 | | -0.60 | 0.87 | | -0.35 | 0.46 | |
| tr\|G1TNS4\|G1TNS4_RABIT;tr | Tubulin alpha chain | | | 0.35 | 1.03 | | 0.74 | 2.95 | | 0.87 | 3.29 | | 0.19 | 0.76 | | 0.25 | 0.85 | | 0.37 | 1.16 | |
| tr\|G1TNY1\|G1TNY1_RABIT | Mitochondrial import inner membrane translocase subunit | | | 4.74 | 4.20 | enriched | 0.00 | 0.00 | | 0.83 | 0.47 | | 0.00 | 0.00 | | 0.00 | 0.00 | | 0.00 | 0.00 | |
| tr\|G1TQM9\|G1TQM9_RABIT;t | 40S ribosomal protein SA (; | RPSA | | -0.80 | 2.24 | | -0.84 | 2.20 | | -0.76 | 2.19 | | -0.88 | 2.44 | | -0.81 | 1.91 | | -0.55 | 1.53 | |
| tr\|G1TR82\|G1TR82_RABIT | Tubulin alpha chain | TUBA4A | | 1.42 | 0.82 | | 1.71 | 2.37 | enriched | 2.94 | 1.49 | enriched | 1.39 | 2.00 | | 0.88 | 0.50 | | 1.24 | 1.18 | |
| tr\|G1TU13\|G1TU13_RABIT;tr | 40S ribosomal protein S17 | | | -0.14 | 0.29 | | -0.17 | 0.38 | | -0.25 | 0.53 | | -0.15 | 0.33 | | -0.13 | 0.33 | | -0.20 | 0.56 | |
| tr\|G1TSH8\|G1TSH8_RABIT;tr\| | CSD domain-containing protein | | | -0.69 | 0.15 | | -0.28 | 0.05 | | 1.98 | 0.73 | | 1.31 | 0.25 | | -0.15 | 0.03 | | 0.02 | 0.00 | |
| tr\|G1TZB5\|G1TZB5_RABIT;tr\| | 60S ribosomal protein L8 | | | -0.99 | 1.94 | | -0.78 | 1.59 | | -0.84 | 1.62 | | -1.02 | 1.43 | | -0.97 | 1.69 | | -0.77 | 1.58 | |
| tr\|G1TT67\|G1TT67_RABIT | VLIG-type G domain-containing protein | | | 0.18 | 0.93 | | -0.07 | 0.27 | | 0.15 | 0.42 | | -1.21 | 1.53 | | -1.30 | 1.23 | | -0.44 | 0.47 | |
| tr\|G1TTN8\|G1TTN8_RABIT | RING-type domain-containing protein | | | 5.13 | 2.29 | enriched | 3.82 | 1.84 | enriched | 4.26 | 2.00 | enriched | 4.92 | 2.22 | enriched | 4.67 | 2.13 | enriched | 3.49 | 1.71 | enriched |
| tr\|G1TTU6\|G1TTU6_RABIT | S-phase kinase-associated protein 1 | | | 2.88 | 3.45 | enriched | 2.88 | 3.52 | enriched | 3.03 | 3.60 | enriched | 3.07 | 3.94 | enriched | 1.19 | 0.51 | | 1.03 | 0.43 | |
| tr\|G1TUB8\|G1TUB8_RABIT | 60S ribosomal protein L11 | ELOA | | -1.11 | 0.61 | | -0.75 | 0.58 | | -1.22 | 0.52 | | -1.48 | 0.62 | | -0.68 | 0.44 | | -1.00 | 0.66 | |
| tr\|G1TUT9\|G1TUT9_RABIT | 40S ribosomal protein S2 | | | 2.82 | 1.28 | enriched | 3.00 | 1.36 | enriched | 2.58 | 1.17 | enriched | 2.80 | 1.26 | enriched | 2.67 | 1.22 | enriched | 2.93 | 1.33 | enriched |
| tr\|G1TXF6\|G1TXF6_RABIT;tr\| | 60S ribosomal protein L27 | RPL27 | | 2.69 | 3.28 | enriched | 2.61 | 3.19 | enriched | 2.02 | 1.76 | enriched | 1.90 | 1.27 | enriched | 0.00 | 0.00 | | 1.13 | 0.56 | |
| tr\|G1U0I7\|G1U0I7_RABIT | Ataxin 2 like | ATXN2L | | -1.34 | 4.22 | enriched | -0.88 | 3.43 | | -1.20 | 4.13 | | -1.30 | 3.58 | | -0.77 | 2.31 | | -0.56 | 2.61 | |
| tr\|G1U0Q2\|G1U0Q2_RABIT | 40S ribosomal protein S15 | RPS15 | | 0.14 | 0.07 | | -0.06 | 0.02 | | -0.33 | 0.19 | | -0.45 | 0.27 | | -0.44 | 0.24 | | 0.00 | 0.00 | |
| tr\|G1U0V2\|G1U0V2_RABIT | Alpha-synuclein | SNCA | | -0.06 | 0.12 | | 0.01 | 0.02 | | 0.20 | 0.44 | | 0.34 | 0.78 | | 0.16 | 0.37 | | 0.63 | 1.45 | |
| tr\|G1U410\|G1U410_RABIT;tr | Tubulin beta chain | TUBB4A | | 1.45 | 0.90 | | 1.24 | 0.76 | | 1.01 | 0.60 | | 0.69 | 0.37 | | 0.87 | 0.50 | | 1.11 | 0.67 | |
| tr\|G1U482\|G1U482_RABIT | 40S ribosomal protein S3a | RPS3A | | 1.11 | 0.27 | | 2.43 | 0.68 | enriched | 2.05 | 0.81 | enriched | 3.42 | 2.50 | enriched | 1.05 | 0.28 | | 1.19 | 0.40 | |
| tr\|G1U5N5\|G1U5N5_RABIT | Lactamase_B domain-containing protein | | | 0.83 | 0.50 | | 0.98 | 0.61 | | 0.89 | 0.54 | | 1.31 | 0.84 | | 1.17 | 0.74 | | 1.05 | 0.65 | |
| tr\|G1U6N2\|G1U6N2_RABIT;t | Acidic ribosomal phosphop | 36B4 | | -0.46 | 1.82 | | -0.55 | 2.22 | | -0.76 | 1.86 | | -0.57 | 1.34 | | -0.84 | 1.26 | | -0.43 | 1.54 | |
| tr\|G1U6X4\|G1U6X4_RABIT | Methionine aminopeptidas | METAP2 | | 0.21 | 0.14 | | 0.00 | 0.00 | | 0.55 | 2.73 | | 0.00 | 0.00 | | 0.00 | 0.00 | | 0.00 | 0.00 | |
| tr\|G1U7L1\|G1U7L1_RABIT | 60S ribosomal protein L28 | | | -1.04 | 3.48 | | -0.81 | 2.97 | | -0.93 | 2.83 | | -1.04 | 2.17 | | -0.73 | 1.33 | | -0.61 | 2.07 | |
| tr\|G1U8P2\|G1U8P2_RABIT | Ribosomal protein L19 | | | -0.60 | 0.26 | | 0.77 | 0.53 | | -1.53 | 1.04 | | -0.48 | 0.22 | | -0.10 | 0.03 | | -1.28 | 0.93 | |
| tr\|G1U9T8\|G1U9T8_RABIT;tr | T-complex protein 1 subuni | CCT4 | | -0.90 | 2.81 | | 0.23 | 1.12 | | -0.12 | 0.46 | | -0.61 | 2.00 | | -1.03 | 3.31 | | -0.72 | 2.40 | |
| tr\|G8ZF20\|G8ZF20_RABIT;tr\| | Alpha-globin 1 (Alpha-globi | HBA1 HBA2 | | 1.20 | 2.47 | | 1.02 | 3.60 | | 0.78 | 2.54 | | 1.04 | 2.29 | | 0.65 | 1.32 | | 1.09 | 4.06 | |
| tr\|P79370\|P79370_RABIT | Rsc protein | rsc | | 2.03 | 3.04 | enriched | 1.00 | 1.96 | | 1.79 | 2.60 | enriched | 1.16 | 1.50 | | 1.39 | 2.46 | | 1.16 | 1.65 | |
| tr\|Q0QEW4\|Q0QEW4_RABIT | Ribosomal protein L18 (Fra | RPL18 | | -0.39 | 3.33 | | -0.44 | 2.73 | | -0.54 | 3.08 | | -0.46 | 1.95 | | -0.63 | 2.72 | | -0.14 | 1.16 | |
